# Monoclonal nephritic factors reveal insights into C3 convertase dynamics and dysregulation

**DOI:** 10.64898/2026.06.11.731599

**Authors:** Seth J. Welsh, Christopher T. Culek, Zhen Xu, Héctor Merinero, Jannik Sichau, Daniel Walls, Renee Goodfellow, Dingwu Shao, Cobey Donelson, Kameron Kruger, Sarah Roberts, Nicole C. Meyer, Angela F. M. Nelson, Anna R. Carmen, Sydney S. Jellison, Stephanie N Cook, Monica D. Hall, Rebecca Franklin, Tina Liu, Jill Hall, Lauren O. Fergus, Nicholas J Schnicker, Christoph Q. Schmidt, Yuzhou Zhang, Carla M. Nester, Richard J.H. Smith

## Abstract

C3- and C5-nephritic factors are potent but poorly understood autoantibodies that dysregulate complement convertases. To date, their underlying mechanisms of action, epitopes, and sequences remain unknown. To address this knowledge gap, we immune profiled B cells from a nephritic factor-positive C3 glomerulopathy patient and identified the first monoclonal C3- and C5-nephritic factors. We present the structure of a C3-nephritic factor bound to a C3 convertase with the convertase protease domain unexpectedly rotated and inhibited. This rotation advances our understanding of complement convertase progression and decay, explains disease-associated convertase variants, and reveals the molecular mechanism by which nephritic factors can either activate or inhibit convertase activity. We also detail heterogeneity within and between C3- and C5-nephritic factors in terms of convertase binding, stabilizing capacity, regulator inhibition, fluid-phase activation, and disease contribution. These findings improve stratification of patients with C3 glomerulopathy, redefine basic C3 convertase dynamics, and provide insights into antibody-mediated modulation of the complement system.

## Main Text

The complement system is a dynamic immune network of humoral and cell-surface proteins that targets pathogens, maintains tissue homeostasis, and modulates adaptive immunity^1–3^.

Consequently, complement dysregulation is linked to infections, autoimmunity, and myriad inflammatory diseases ranging from neurodegeneration to cancer^4–6^. Regardless of the initiating event, almost all complement activity converges onto the highly dynamic and tightly regulated multimolecular enzymatic complex known as the alternative pathway C3 convertase. The C3 convertase cleaves C3 into the anaphylatoxin C3a and the opsonin C3b, thereby promoting rapid pathway amplification, immune complex processing, and localized increases in C3b density that are a prerequisite for C5 priming and the transition to a C5 convertase^7–9^. The subsequent activation of C5 into C5a and C5b results in proinflammatory cellular chemotaxis and initiation of the terminal pathway membrane attack complex^10^.

In some individuals, this process is dysregulated by genetic and/or acquired drivers, as occurs in the complement-mediated disease C3 Glomerulopathy (C3G)^11–13^. In C3G, overactivation of the alternative pathway of complement is most frequently driven by heterogenous autoantibodies known collectively as nephritic factors or “Nefs”^14,15^. Nefs bind unidentified epitopes on complement convertases and promote convertase formation and stability through unknown mechanisms^16–21^. The challenges in Nef study are significant and reflect both the polyclonal heterogeneity of Nefs as well as the dynamic, extremely labile, and fleeting half-life (∼90s *in vivo*) of their multimolecular convertase targets^22,23^. As evidence, a mysterious complement-activating “nephritic factor” was first described nearly 60 years ago, yet to date, a monoclonal Nef and its epitope remain unknown^24–26^.

To address this knowledge gap, we performed single-cell immune profiling of B cells isolated from a Nef-positive C3G patient. From the resulting B cell receptor (BCR) dataset, we identified the first monoclonal sequences for two classes of convertase-targeting Nefs: properdin-independent (C3Nefs) and properdin-dependent (C5Nefs). We functionally characterized both classes, mapped the binding epitope for a C3Nef using cryo-EM, and showed that C3Nef binding competes with the positive regulator of complement, properdin. Unexpectedly, the C3Nef-bound convertase displayed a 180° upward rotation of its protease domain, Bb, into a novel “Bb-up” position not observed in any current structures. The Bb-up position, when stabilized by C3Nef, resulted in convertases that were paradoxically inhibited. In contrast, C5Nefs stabilized convertase in the Bb-down position, maintaining its catalytic competence. Clinical biomarker data support this mechanistic dichotomy revealing that C3Nefs and C5Nefs possess distinct characteristics that underlie different C3G disease subtypes. Incidentally, the dynamic rotation of Bb captured by C3Nef also provides a structural mechanism for convertase variants associated with decay-accelerator inhibition and offers a new model of regulator-mediated decay. These multifaceted results define mechanistic distinctions between C3Nefs and C5Nefs and provide novel insights into C3 convertase dynamics and complement modulation by immunoglobulins.

Given the overwhelming success of two recently FDA-approved convertase-targeting drugs, Empaveli and Fabhalta^27,28^, our findings suggest a potential path toward antibody-based complement-modulating therapies that exploit native convertase dynamics.

### Identification of C3- and C5-nephritic factors from C3G patient B cells

To identify candidate Nefs, we performed single-cell BCR sequencing of peripheral blood mononuclear cells (PBMCs) isolated from a C3G patient co-positive for both C3Nefs and C5Nefs and whose complement profile was consistent with Nef-driven disease (Fig.1A, and fig. S1). PBMCs collected nine months apart were FACS-enriched for antigen-experienced B-lineage cells and sorted into two groups prior to single-cell immune profiling. Group 1 (“Memory B”) consisted entirely of memory B cells (IgD^−^CD27^+^CD21^+^), whereas Group 2 (“Pooled”) combined circulating plasma cells and plasmablasts (CD27^Hi^CD38^Hi^) with marginal zone-like cells (IgD^+^CD27^+^) and atypical memory B cells (IgD^−^CD21^−^) (fig. S2, A and B; table S1).

**Fig. 1.**
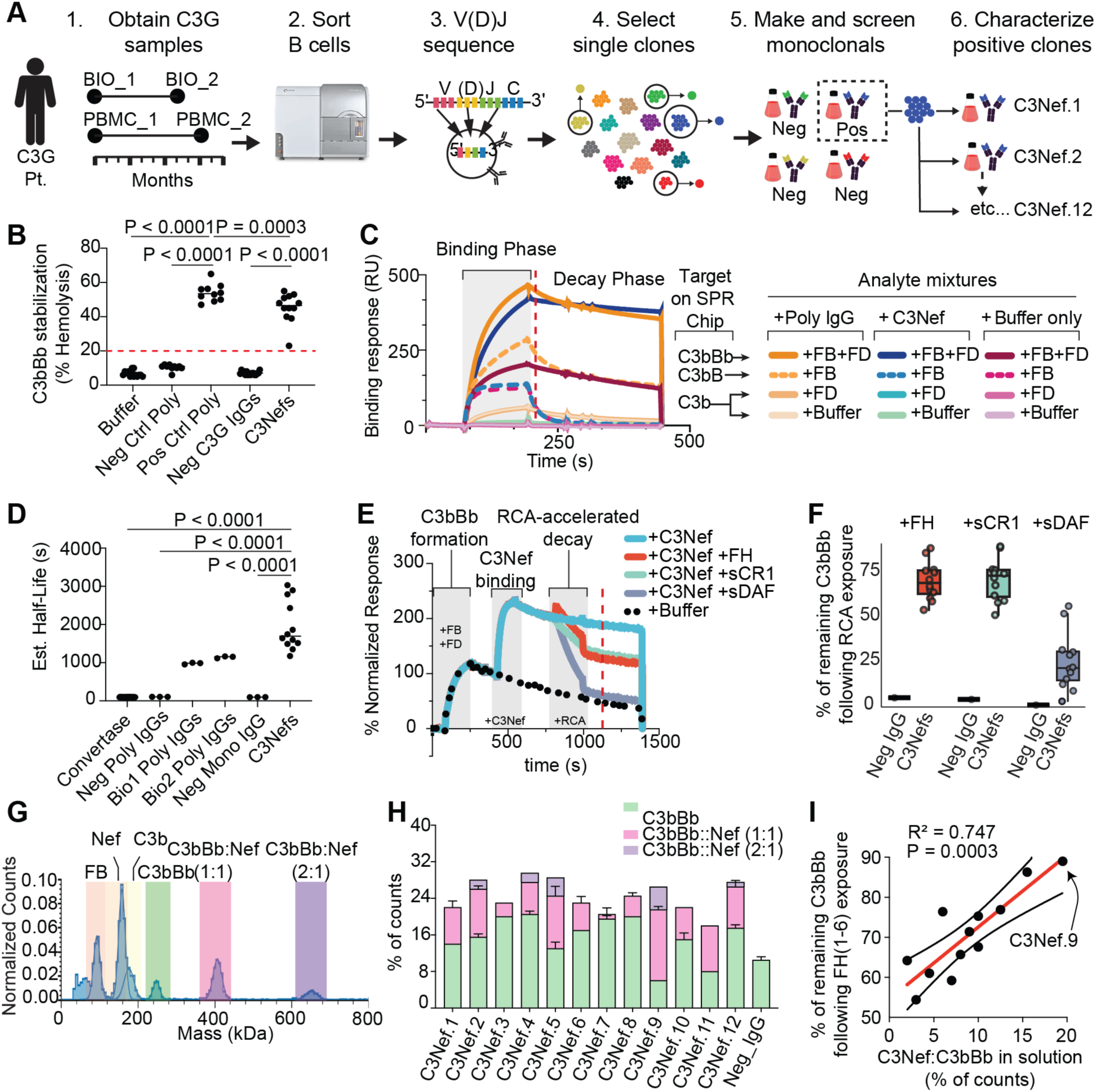
Identification of monoclonal nephritic factors. (**A**) Workflow for patient-derived C3Nef identification. (**B**) Hemolytic assay showing the convertase-stabilizing properties for 12 monoclonal C3Nefs compared with buffer, polyclonal neg controls (n=10), polyclonal pos controls (n=10), and non-Nef monoclonal IgGs from the same C3G patient (n = 16). Ordinary one-way ANOVA with Tukey’s multiple comparison test. Dotted red line indicates clinical cutoff for Nef positivity. (**C**) Representative SPR sensorgram overlay showing the differences in the binding and decay of recombinant complement convertase proteins to C3b-coated chip in the presence of buffer, C3Nef.9, or polyclonal serum IgGs obtained from the same C3G patient and timepoint as the C3Nefs. Vertical dashed red line indicates report point. (**D**) Jitter plot showing the estimated half-life decay values for controls (n=3) compared with monoclonal C3Nefs. Each C3Nef dot is the average of 3 technical replicates. *P* values from one-way ANOVA with Tukey’s multiple comparison test. (**E**) Representative SPR sensorgram overlay showing the ability of C3Nef to inhibit RCA-mediated decay of C3bBb by factor H domains 1-6 (FH); soluble complement receptor 1 domains 1-3 (sCR1); and soluble decay accelerating factor domains 1-4 (sDAF). Vertical dashed red line indicates report point. (**F**) Jitter plot showing the report point results from **E** for of all 12 C3Nefs. (**G**) Representative histogram overlay of mass photometry (MP) peak counts showing C3Nef.9 binding to C3bBb convertase in solution. (**H**) Bar graph quantifying the peak % counts for C3bBb alone (green), monovalent C3Nef binding to C3bBb (1:1; pink), or bivalent C3Nef binding to C3bBb (2:1; purple). (**I**) Correlation between the ability of each C3Nef to bind C3bBb in solution (y axis; % of counts) and its ability to inhibit Factor H domains 1-6 (FH)-mediated C3bBb decay. Red line denotes simple linear regression with 95% CI (dotted black lines). Two-tailed Pearson’s R squared and P values listed.

Following sequencing, we used enclone software to cluster BCR sequences informatically into distinct clonotypes ^29^. A clonotype is a cluster of unique B cells with similar V(D)J region sequences predicted to have arisen from a common progenitor. In total, we captured 5,730 individual cells with productive V(D)J rearrangements that clustered into 5,274 unique clonotypes (fig. S2, B to D).

From our BCR dataset, we applied non-mutually exclusive ranking criteria to select 27 clonotypes for initial Nef screening. Briefly, our ranking criteria used evidence-based inference from Nef literature and prioritized clonotypes harboring IgG constant regions, high levels of expression, and clonal persistence across time, among other features (see Methods). From each selected clonotypic cluster we first chose a single unique clone to express as paired variable heavy (VH) and variable light (VL) chains. Given that most Nef assays utilize IgG fractionated serum, and because the antibody variable domain is primarily responsible for Nef convertase-stabilizing properties ^30,31^, we expressed all clones in a common IgG1 constant heavy region (TWIST Bioscience, Inc) (Fig. 1A; table S2). Light chain constant regions were assigned either a shared kappa or shared lambda to match each clonotype’s original VL isotype (table S3). In addition to our 27 patient-derived clones, we also tested a previously reported monoclonal IgG with C3Nef-like function generated *ex vivo* by cloning the BCR from pokeweed mitogen-stimulated PBMCs obtained from a patient with membranoproliferative glomerulonephritis ^32^. In total, our initial hemolytic screen tested 28 unique monoclonals for C3 convertase-stabilizing activity in the presence and absence of the positive regulator of complement activity, properdin. From this screen, we identified one clone with properdin-independent convertase-stabilizing properties (Clone 1) and a second clone that showed a modest increase in convertase stabilization after the addition of properdin (Clone 4) (fig. S2, E to G). Based on these results, we returned to the respective parental clonotype clusters and generated all remaining subclones, ultimately yielding 14 cells (12 unique VH+VL pairs) that were properdin-independent clones (henceforth “C3Nefs”), and 8 cells (6 unique VH+VL pairs) that were properdin-dependent clones (henceforth “C5Nefs”) (fig. S3; tables S2 and S3).

### Properdin independent C3Nefs bind and stabilize the C3bBb convertase

To confirm that our properdin-independent monoclonals possessed Nef-associated properties, we completed a series of biophysical assays. Importantly, all 12 C3Nef clones tested positive in a hemolytic C3 convertase stabilizing assay used for clinical C3Nef detection (Fig. 1B, and fig. S3, A and B). Second, for comparison, we showed that polyclonal IgGs from the patient’s contemporaneous serum bound broadly to C3b, pro-convertase C3bB, and fully formed convertase C3bBb by surface plasmon resonance (SPR) (Fig. 1C). In contrast, each of the 12 unique monoclonal C3Nefs showed minimal reactivity to C3b and C3bB but strong, selective binding to C3bBb (Fig. 1C; fig. S3F, S4 and S5). Despite shared clonality, the 12 C3Nefs differed markedly in their ability to bind and stabilize C3bBb as measured by estimated half-life values (Fig. 1D; fig. S3G, S6 to S9). Third, we generated C3bBb convertase on a chip, stabilized it with C3Nef, and then challenged it with three known regulators of complement convertases: factor H domains 1-6 (FH); soluble complement receptor 1 domains 1-3 (sCR1); and soluble decay accelerating factor domains 1-4 (sDAF) (Fig. 1E; fig. S10). All 12 C3Nefs reduced FH-and sCR1-mediated decay, while sDAF-mediated decay was substantially less affected (Fig. 1F; fig. S11). Lastly, mass photometry confirmed that all 12 C3Nefs formed C3Nef:C3bBb complexes in solution, with five binding bivalently to two C3bBb convertase complexes (Fig. 1, G and H). The capacity of each C3Nef to form a complex with C3bBb in solution was highly correlated with its ability to inhibit convertase decay by blocking the fluid-phase regulator FH as measured by SPR (Fig. 1I). Together, these data demonstrate that the 12 monoclonal autoantibodies isolated from the C3G index patient bind and stabilize the C3bBb convertase and function as C3Nefs.

### C3Nefs are distinct from C5Nefs

In addition to the 12 C3Nefs, we identified 6 unique C5Nefs whose optimal convertase binding and stabilizing activity required properdin (fig. S3, C to E; tables S2 and S3). Sequence analysis showed that C3Nefs and C5Nefs belonged to distinct clonotypes arising from separate V(D)J recombination events with markedly different Ig CDRs (Fig. 2A). Specifically, C3Nefs used the heavy gene variable region IGHV3-21 recombined to IGHJ4 and paired with a kappa light chain (IGKV1-17 → IGKJ1). By contrast, C5Nefs used IGHV5-51 recombined to IGHJ3 and paired with a lambda light chain (IGLV3-19 → IGLJ2) (table S2). With respect to class-switch recombination, only one C5Nef had switched to an IgG1 constant region; all other C3Nefs and C5Nefs were expressed as an IgG3 *in vivo*.

**Fig. 2.**
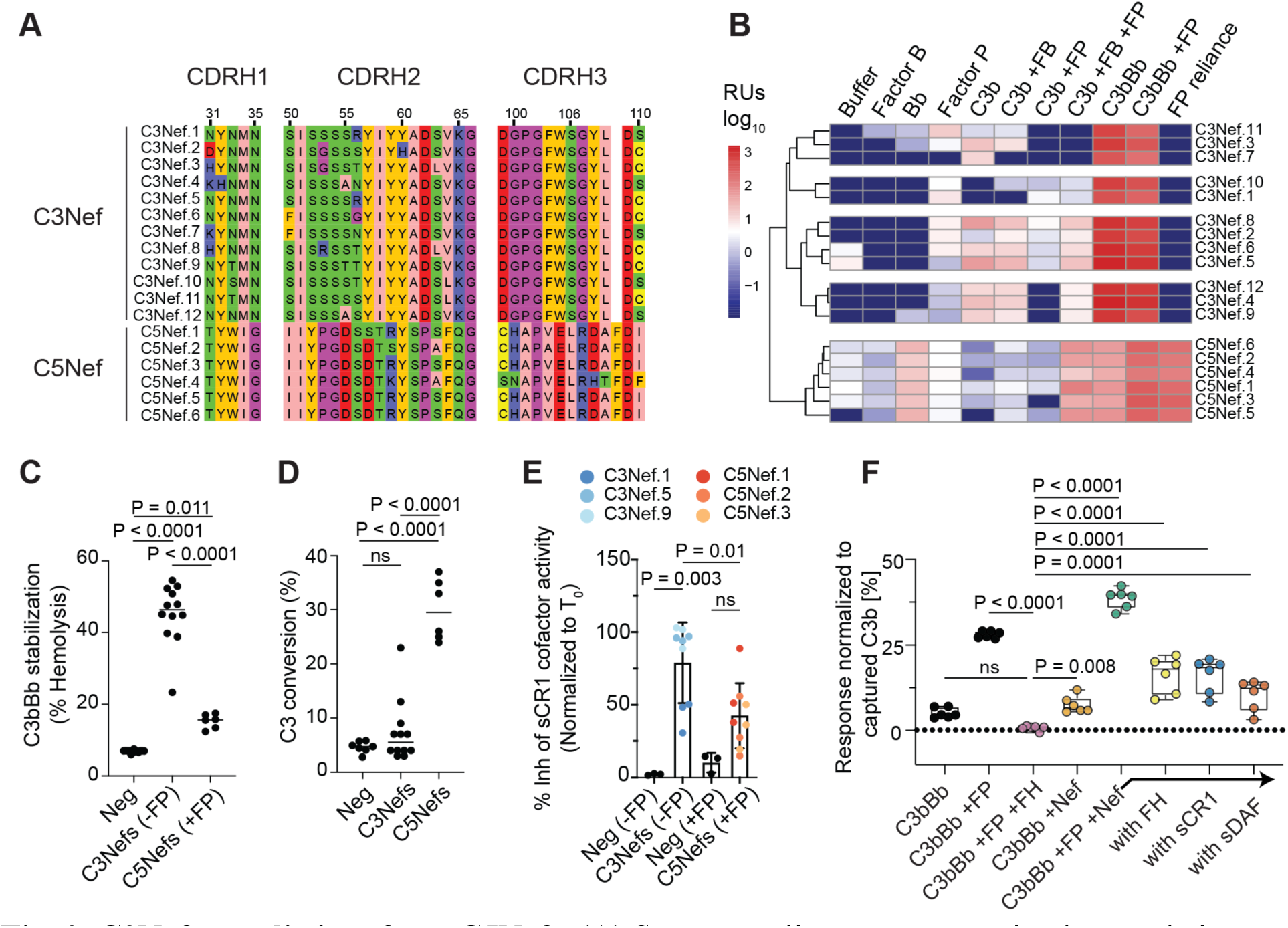
C3Nefs are distinct from C5Nefs. (**A**) Sequence alignment comparing heavy chain complementarity determining regions (CDRs) 1-3 for C3Nefs (top) and C5Nefs (bottom) using Kabat numbering. (**B**) Hierarchical clustering heatmap for all 12 C3Nefs and all 6 C5Nefs exposed to various analyte mixtures (top). SPR binding RU values shown in log_10_ scale. Individual Nef ids listed on the right. “FP reliance” calculated as: (C3bBb + FP RUs) – (C3bBb RUs). Values show the mean from two independent runs. FB, factor B; FP, factor P/properdin; FD, factor D; RU, resonance units. (**C**) Jitter plot showing differences between C3Nefs and C5Nef’s ability to stabilize membrane bound C3bBb convertase. FP, properdin. (**D**) Jitter plot showing differences between C3Nefs and C5Nef’s ability to induce fluid-phase activation of complement. (**E**) Jitter plot bar graph overlay showing the ability of C3Nefs and C5Nefs to inhibit sCR1(domains 15-17) cofactor-mediated cleavage of C3b into iC3b in solution. (**F**) Jitter plot showing the ability of C5Nefs to bind a properdin-stabilized C3bBb convertase and block RCA-mediated decay. FP, factor P; FH, factor H domains 1-3; sCR1, soluble complement receptor 1 domains 1-3; sDAF, soluble decay accelerating factor domains 1-4.

To confirm nephritic-factor activity and compare the binding profiles of C3Nefs with C5Nefs, we performed extensive SPR characterization for all 18 monoclonal antibodies which yielded multiple insights. First, C3Nefs and C5Nefs exhibited distinct binding profiles (Fig. 2B). Second, properdin inhibited C3Nef binding to convertase but enhanced C5Nef binding (fig. S12 to S14). Third, C3Nefs showed no reactivity toward C4/C4b or C5/C5b6 (fig. S15A). Fourth, C3Nef bound weakly to C3b and the C3bB proconvertase; in contrast, C5Nefs preferentially recognized Bb and multimeric Bb-properdin complexes and showed almost no reactivity to factor B, suggesting the Ba domain might block access to the composite epitope recognized by C5Nefs (Fig. 2B). The strongest response for all C3Nefs and C5Nefs remained the fully formed convertase.

Unsupervised clustering of the SPR binding profiles cleanly separated C3Nefs from C5Nefs but yielded groupings that differed slightly from DNA-sequence phylogeny, indicating that similar binding phenotypes can arise via different somatic-hypermutation pathways (Fig. 2B, fig. S15, B and C). Functionally, *in vitro* analysis showed that C3Nefs and C5Nefs dysregulate complement by distinctly different mechanisms: C3Nefs acted independently of properdin and preferentially stabilized membrane-bound convertases, albeit in a paradoxically non-functional conformation when bound (Fig. 2C; details below). In contrast, C5Nefs were properdin dependent and activated C3 consumption in the fluid phase (Fig. 2D). C3Nefs were also highly effective and better than C5Nefs at blocking the cofactor activity of sCR1 domains 15-17; however, neither Nef significantly prevented the cofactor activity inherent in FH domains 1-6 (Fig. 2E; fig. S15, D to G). To test whether C5Nefs could protect convertases from regulator-accelerated decay only, we built a properdin-stabilized C3bBb convertase on a chip, added C5Nef, and then exposed the C3bBbP:C5Nef complex to RCAs (Fig. 2F; fig. S16). In this setup, properdin alone was unable to prevent complete RCA-accelerated decay of the C3bBbP complex (fig S17, A to D); however, the addition of C5Nefs stabilized the C3bBbP complex and inhibited decay by FH and sCR1 but less so by sDAF (Fig. 2F; fig S17E).

Collectively, these monoclonal data confirm that C3Nefs and C5Nefs both share an ability to inhibit various RCAs yet differ in their sequences, binding profiles, and stages of convertase complex engagement– stabilizing convertases that are either inactive (C3Nefs) or active (C5Nefs) (details below).

### Cryo-EM structure of C3Nef bound to the C3bBb convertase complex

While multiple C3 convertase structures exist, the conformational state recognized by Nefs is unknown. To identify the C3Nef epitope on C3bBb, we performed cryogenic electron microscopy (cryo-EM) utilizing subclone C3Nef.9, which formed the most stable C3Nef:C3bBb complex in solution (Fig. 1, G and H). Unexpectedly, our early attempts to prolong the C3Nef:C3bBb interaction using well-documented convertase-stabilizing methods such as Ni^2+^, factor B mutants (p.D279G or p.K350N), or the addition of staphylococcal complement inhibitor (SCIN) protein all reduced C3Nef binding in solution (fig. S18) ^33–37^. Ultimately, grids were prepared using a magnesium-supplemented mixture of recombinant C3b, factor B, factor D, and C3Nef.

From these grids, we obtained two density maps. The first was a high resolution 3-Å reconstruction of the 1:1 C3Nef:C3bBb complex, consisting of the C345c domain of C3b, the Von Willebrand (vWA) and serine protease (SP) domains of Bb, and one Fab domain of the C3Nef IgG (Fig. 3A; fig. S19 and S20; tables S4 and S5). The second was a lower resolution reconstruction from the same data that showed additional density for the macroglobulin (MG) domains of C3b (fig. S20, D and E, tables S4 and S5). An atomic model built into the 3-Å map showed C3Nef contacted both the C345c domain of C3b and the vWA domain of Bb (Fig. 3A to C; table S6). In this trimeric structure, the α1 and α3 helices of C345c drove the interaction with the C3Nef VH domain engaging all three CDRH loops: Specifically, CDRH3 residues P120, G121, F122 and W123 contacted C345c residues F1659, S1655, L1529 and P1662 on one side of the α3 helix, while CDRH1 N50, and CDRH2 S71, S73, and Y76 contacted C345c V1658, F1659, and E1654/G1650, respectively, on the opposite side of the α3 helix (Fig. 3B). While C345c contacted both heavy and light chains of the C3Nef, Bb contacted only the CDRL2 loop (Fig. 3, C and D; fig. S21). V(D)J sequence comparison of the 12 C3Nefs revealed modest heterogeneity within CDRH1, CDRH2, and CDRL3 regions that likely contributed to differences in binding and function observed among the C3Nefs. Nevertheless, 11 of the 16 contact residues were completely conserved across all 12 C3Nefs, suggesting a shared epitope (fig. S22, A and B). In support of this, robust binding for all C3Nefs required the fully formed C3bBb convertase (Fig. 3E; fig. S13).

**Fig. 3.**
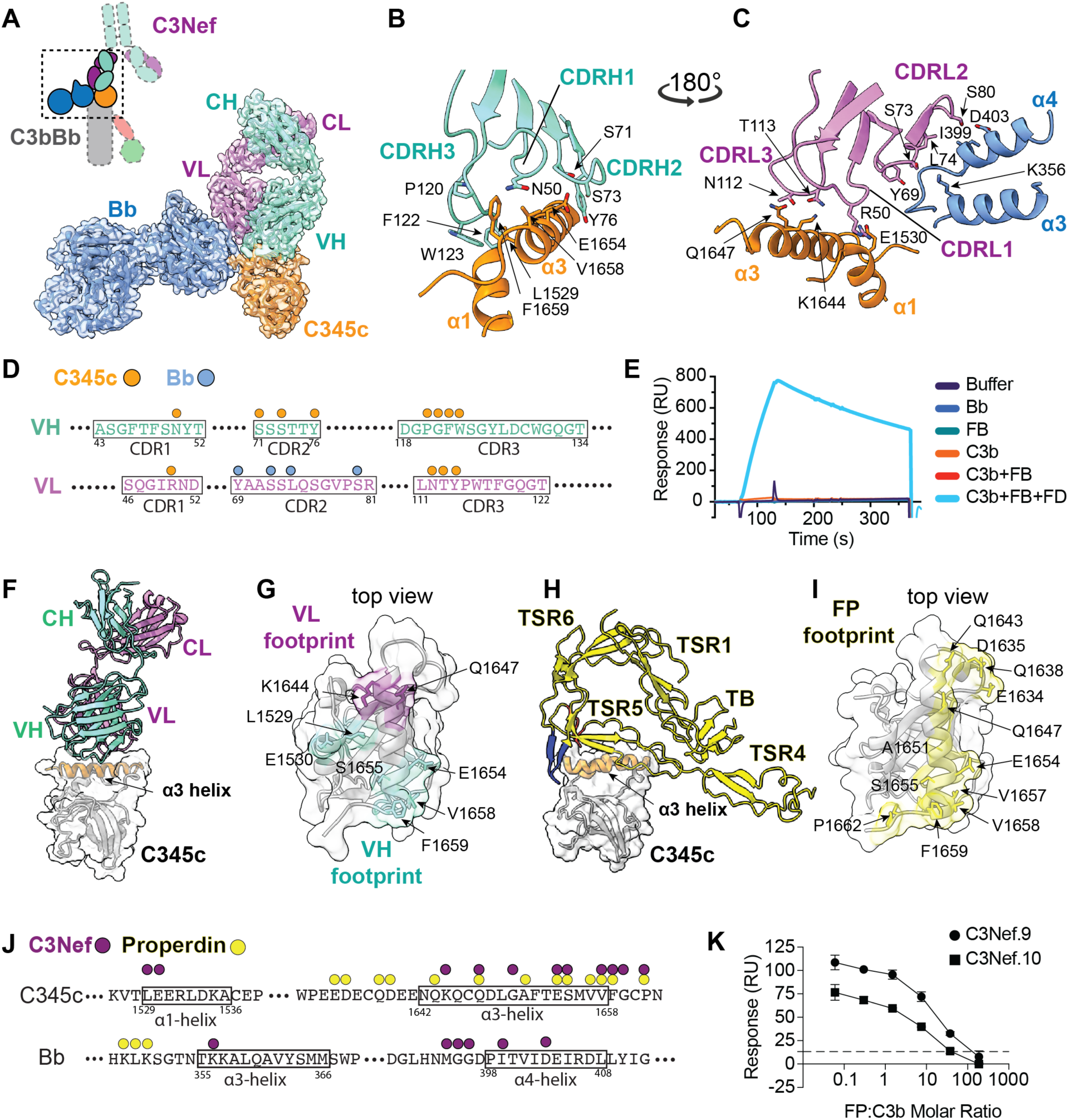
C3Nefs bind the C345c domain of C3b and the Von Willebrand domain of Factor B in competition with properdin. (**A**) Upper left: Diagram of a C3Nef-bound C3bBb convertase depicting the high-resolution structures captured by cryo-EM (dashed box). Main structure: Cryo-EM map at 2.99 Å resolution with fitted model of C3Nef stabilized C3bBb complex. C3Nef Fab domain region heavy chain in aquamarine, Fab light chain in purple, C345c domain of C3b in orange and Bb in blue. (**B**) Zoomed-in model showing interaction of C3Nef heavy chain with C345c domain of C3b. Not all interactions are shown. (**C**) Zoomed-in model of C3Nef light chain interaction with C345c domain of C3b and the Von Willebrand domain type A of Bb. Not all interactions are shown. (**D**) Linear model showing individual interaction sites between the VH and VL CDR regions of the C3Nef with the C3b C345c domain (orange) or the Bb domain of FB (blue). (**E**) Representative SPR sensorgram showing the interaction of C3Nef.9 with C3bBb is highly specific to the fully formed convertase (Analyte: C3b +FB +FD). (**F**) Side view of C3Nef Fab binding the α_3_-helix of C3b’s C345c domain. (**G**) Top-down view showing the footprint of VH and VL binding on the apical α3 and α1 helices of C345c domain of C3b (heavy chain footprint in aquamarine, light chain footprint in purple). (**H**) Side view of properdin heterodimer TB-TSR1:TSR4-6 binding to the α_3_-helix of C3b’s C345c domain. Properdin’s “thumb” and “finger” shown in red and blue, respectively ^44^. **(I)** Top-down view showing the footprint of properdin (yellow) on the C345c domain of C3b. (**J**) Linear schematic showing select amino acid residues of C345c (top) or Bb (bottom) that contact C3Nef (purple dots) or properdin (yellow dots). (**K**) Reduction in C3Nef binding to C3bBb complexes generated in the presence of increasing concentrations of properdin. n = 2 technical replicates. For all panels: CH, constant heavy chain; CL, constant light chain; VH, variable heavy chain; VL, variable light chain; CDR, complementary determining region; CDRH, complementary determining region heavy; CDRL, complementary determining region light; FB, factor B; FD, factor D; RU, resonance units; TSR, thrombospondin type1 repeat; TB, transforming growth factor-beta binding protein-like domain; FP, factor P/properdin.

### C3Nef binding competes with properdin

Comparison of our C3Nef:C3bBb complex with current properdin-bound convertase models ^38–44^ revealed analogous binding sites for C3Nef and properdin on the apical α3 helix of the C345c domain of C3b (Fig. 3, F to I; fig. S23, A to B). In this interaction, C3Nef heavy chain CDRs functionally mimicked the ‘thumb’ and ‘finger’ of properdin thrombospondin repeats (TSRs) 5 and 6 (also known as “stirrups”) (fig. S23, C to F). Footprint analysis of C345c revealed overlap between C3Nef and properdin interaction surfaces with properdin contacting a larger region across the α3 helix than C3Nef (Fig. 3, F to J). However, C3Nef made unique contacts with the α1 helix of C345c and displayed markedly greater interaction with Bb than properdin, contacting both the α3 and α4 helices of Bb (Fig. 3J). In support of competitive binding, orthogonal SPR assays demonstrated that pre-incubation of C3bBb with increased properdin reduced subsequent C3Nef binding (Fig. 3K). Interestingly, some of the key C345c residues contacted by both C3Nef and properdin are also part of the epitope recognized by a recently identified single-domain VHH nanobody capable of inhibiting alternative pathway and C5 convertase activity ^45^.

### C3Nef epitope is associated with 180° rotation of Bb

Surprisingly, comparison of Bb’s orientation in our C3Nef:C3bBb structure with previously reported properdin(±SCIN):C3bBb structures, revealed an unexpected ∼180-degree rotation of Bb around a perpendicular axis relative to C345c that was centered on the metal ion-dependent adhesion site (MIDAS) (Fig. 4, A to C) ^34,38,43^. We refer to this new orientation as the “Bb-up” configuration. To examine if the Bb-up configuration was necessary for Nef binding we titrated the convertase inhibitor SCIN into a convertase-forming analyte mixture prior Nef exposure.

**Fig. 4.**
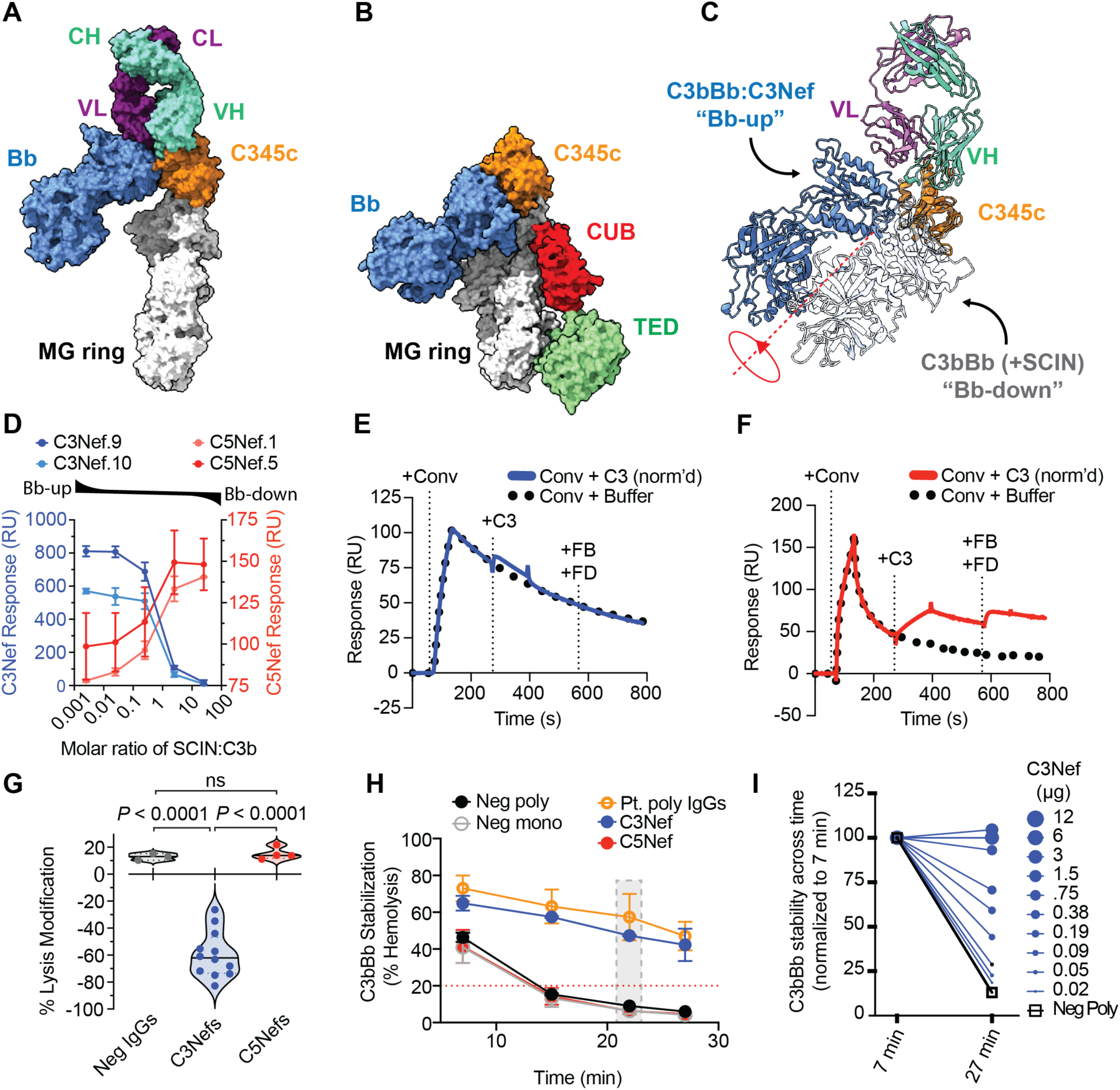
C3Nefs bind C3bBb in Bb-up configuration, inhibiting the C3 convertase. (**A**) Cryo-EM surface area model showing 3.74 Å resolution for C3Nef-stabilized C3bBb complex with partial resolution for MG ring. C3Nef Fab region heavy chain is colored in aquamarine, light chain in purple, C3b C345c in orange, MG ring in grey and Bb in blue. (**B**) Structure of SCIN-stabilized C3bBb complex (PDB 2WIN) ^34^. SCIN is not shown. (**C**) 180° reorientation of Bb in the C3Nef structure compared with 2WIN, aligned on the C345c domain. (**D**) SPR titration data showing that SCIN-induced Bb-down configuration results in decreased binding of C3Nefs (blue) but increased binding of C5Nefs (red). (**E**) SPR sensorgram overlay showing the ability of a C3Nef-stabilized C3bBb convertase to bind, but not cleave, free C3. See Methods for details. (**F**) SPR sensorgram showing the ability of C5Nef-stabilized C3bBbP(mono) convertase to cleave C3; the resulting C3b opsonizes the SPR chip and forms new C3bBb convertase upon addition of +FB+FD. (**G**) Guinea pig hemolytic assay data showing C3Nef-induced inhibition of lysis. (**H**) Hemolytic assay showing the ability of index patient polyclonal IgGs and monoclonal C3Nef.5 to stabilize the C3bBb convertase as measured by prolonged hemolysis across time. Shaded grey box indicates timepoint and lysis values reported out clinically. Dotted red line indicates clinical cutoff for positive result (20% lysis). n = 2 technical replicates. Neg mono, C3Nef, and C5Nef tested at 0.75µg. Polyclonal IgG at 300µg. Neg poly at 150µg. (**I**) Increases in monoclonal C3Nef.5 concentration results in increased C3bBb stability across time. For all panels: CH, constant heavy chain; CL, constant light chain; VH, variable heavy chain; VL, variable light chain; MG, macroglobulin; CUB, complement C1r/C1, Uegf, Bmp1 domain; TED, thioester-containing domain; SCIN, staphylococcal complement inhibitor; RU, resonance units; Conv, C3bBb convertase; FB, factor B; FD, factor D; poly, polyclonal; mono, monoclonal.

SCIN facilitated the Bb-down configuration, which reduced C3Nef binding but enhanced C5Nef binding (Fig. 4D). Structural comparison of Bb-up with known convertase models aligned on the MG ring showed a 9-Å shift in the C-terminus of the C345c α3 helix in Bb-up as compared to the 2WIN model (fig. S24, A to B) ^34,43^. Notably, this region of C345c contains the C-terminal residue N1663, which possesses rotational freedom and is central to Mg^2+^ coordination in the MIDAS region of the C3bBb complex. While Bb-up maintained the overall MIDAS architecture, there was a significant relocation of the MIDAS Mg^2+^-coordinating residues S278, S280, and T353 of Bb relative to the known structures of 2WIN and 9U62 (fig. S24, C to E). This rotational repositioning of Bb greatly reduced the contact surface of Bb on C345c (fig. S25).

### C3Nef-binding inhibits the C3 convertase

The displacement of Bb in the Bb-up model raised the question of whether a C3Nef-bound convertase remains functionally active. To assess the plausibility that a C3Nef-stabilized convertase could bind its C3 substrate, we first aligned our Bb-up model with the recently published model of C3 substrate bound to properdin-stabilized C3bBb (fig. S26A) ^43^. With this alignment, we observed a small amount of steric hindrance between Bb-up’s Bb and the overlayed C3 (fig. S26B); however, given the flexibility of the C345c domain, it was unclear whether this minor overlap would prevent C3 binding *in vivo*. More prominent was the rotational displacement of the serine protease domain of Bb from its ligand – the scissile loop of the C3a ANA domain (fig. S26, C and D), suggesting that Bb-up may prevent C3 cleavage. To test C3Nef-induced convertase inhibition directly, we immobilized C3Nef on a SPR chip, built C3bBb convertases on the C3Nef paratopes, and then exposed the Nef-stabilized C3bBb convertases to C3 (fig. S27). This approach ensured that all convertases exposed to free C3 were bound by C3Nef. Under these conditions, the C3Nef:C3bBb complexes bound, but could not cleave, free C3 (Fig. 4E; fig. S28). By contrast, the same assay using C5Nef-bound convertases in the presence of monomerized properdin (“P(mono)”; fig. S29) ^41^, resulted in C5Nef-bound convertases cleaving free C3 into C3b (Fig. 4F; fig. S30), with the newly cleaved C3b opsonizing the SPR chip and supporting the generation of additional C3bBb convertase upon addition of FB and FD. As orthogonal confirmation, we performed a hemolytic assay in which guinea pig RBCs were incubated with normal human serum (NHS) supplemented with increasing concentrations of C3Nefs or C5Nefs. Here, C5Nefs did not affect NHS-mediated RBC lysis; however, C3Nefs produced a dose-dependent inhibition of lysis (Fig. 4G; fig. S31). In aggregate, these data indicate that C3Nefs bind convertases in the Bb-up orientation, rendering them catalytically inactive, while C5Nefs bind convertases in the presence of properdin in the Bb-down orientation, thereby permitting ongoing convertase activity.

To explore the dynamics of C3Nef-driven complement dysregulation we used a hemolytic-based C3 convertase stabilization assay in which a low-density of C3 convertase is built on the surface of sheep erythrocytes to which patient-purified IgGs are added. The readout of convertase activity is the percentage of hemolysis recorded 20-30 min after IgG addition, by which time unbound C3bBb convertases have naturally decayed. Addition of either polyclonal IgGs or monoclonal C3Nefs obtained contemporaneously from the index patient both prevented convertase decay and extended lysis across time (Fig 4H). When we tested monoclonal C3Nef.5 across a 500-fold range, increasing C3Nef concentrations produced progressively greater convertase stability (Fig. 4I) yielding a two-phase concentration-response curve: at low concentrations (phase I), increases in C3Nef levels increased lysis; however, beyond a threshold (phase II), further increases in C3Nef levels decreased lysis (fig. S32A). Time-course analysis revealed distinct decay slopes for these two phases: in phase I, the decay slope of C3Nef-bound convertase matched that of unbound convertase decay, whereas in phase II, the C3Nef-bound slope was flat, consistent with target saturation resulting in enhanced convertase stabilization paired with catalytic inhibition (fig. S32, B and C). These same patterns were observed using increasing concentrations of patient purified polyclonal IgGs (fig. S32, D and E). In aggregate, these data indicate that increasing C3Nef concentration generates more “paused” C3bBb convertases that persist across time, and above a critical concentration, excess C3Nef inhibits all convertases present. This biphasic inhibitory effect is unique to C3Nefs as C5Nefs showed only monotonic concentration-dependent increases in lysis (fig. S3D).

### C3Nefs drive the clinical profile of the C3G patient

To assess how monoclonal Nefs impact C3G disease progression, we compared the single-cell transcriptomic BCR profiles of C3Nef- and C5Nef-expressing B cells with the longitudinal complement biomarker data from the index patient. As expected, the clinically detected polyclonal C3Nefs declined across time, a trend paralleled by a reduction in the frequency of C3Nef-positive B cells and the per-cell C3Nef transcript count (Fig. 5A to C). Although few C3Nef-expressing B cells persisted at the second timepoint, those captured expressed C3Nefs with slightly higher estimated half-life binding affinities, consistent with affinity maturation (fig. S33A). Per cell, C5Nef transcript levels were near-baseline whereas C3Nef transcript levels were ∼50-1,000-fold higher, indicating that C5Nefs were likely retained as surface BCRs in contrast to C3Nefs, which were likely secreted into circulation as antibodies (Fig. 5D). Supporting this distinction, C5Nef-expressing cells were more frequently derived from the memory B-cell compartment than C3Nef-expressing cells (28% vs. 14%) (table S2) and clinical polyclonal C5Nef levels for the index patient fell just below clinical cutoff in the second timepoint (fig. S1). To measure the potential of each Nef subtype for fluid phase dysregulation, we quantified changes in complement biomarker profiles in normal human serum incubated at 37° for 30 min following the addition of monoclonal C3Nefs, C5Nefs, or negative control IgGs. Consistent with the immunofixation electrophoresis data, C5Nefs were capable of driving fluid phase activation as evidence by increased levels of Ba, Bb, C3d, and sC5b9 (Fig. 5E). Effects on properdin were not statistically different between C3Nefs and C5Nefs but trended lower with C5Nefs (fig. S33B). In aggregate, these findings show that monoclonal Nefs reproduced the complement dysregulation profile of the patient from whom they were derived.

**Fig. 5.**
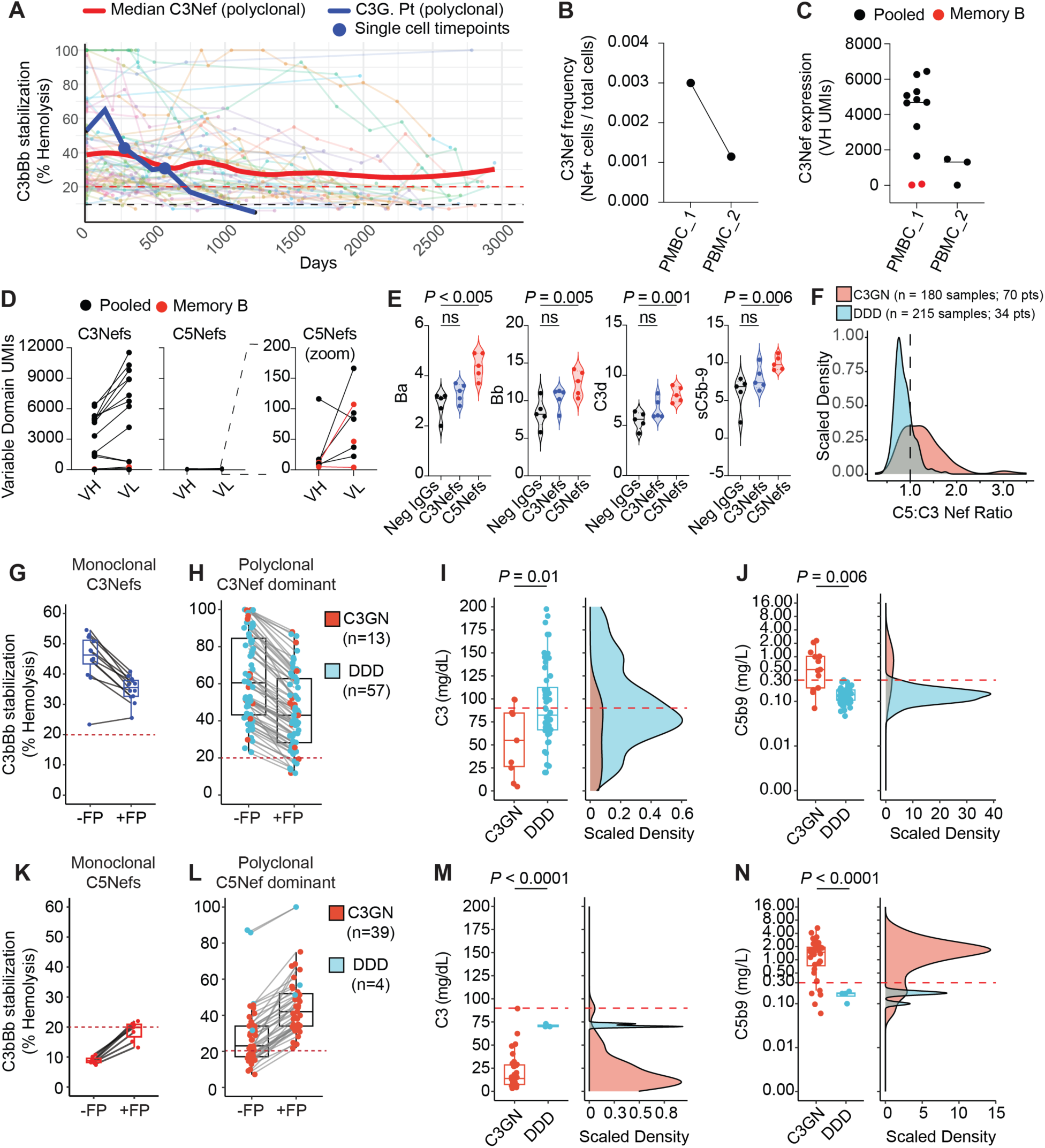
C3Nefs drive DDD, C5Nefs drive C3GN. (**A**) Clinical polyclonal C3Nef values for 394 C3G patients across time. Blue line highlights index C3G patient from whom monoclonal Nefs were identified. Enlarged blue dots show timepoints of PBMC collection. Thick red line shows smoothed median for C3Nef scores for all patients. Horizontal dotted red line indicates the clinical cutoff for positive result (20%). Horizontal dotted black line indicates the limits of assay detection (∼10%). (**B**) Frequency of individual C3Nef-expressing B lineage cells from timepoints 1 and 2 as a percentage of total cells captured at each timepoint. (**C**) Reduced C3Nef expression across time as shown by decreases in unique molecular identifier transcript levels for VH chain of C3Nefs. (**D**) C3Nefs are highly expressed, C5Nefs are not, as determined by UMI transcript levels for paired VH and VL chains. (**E**) C5Nefs activate complement in human serum as determined by increases in biomarkers Ba, Bb, C3d, and sC5b-9 from normal human serum supplemented with monoclonal Neg IgGs, C3Nefs, or C5Nefs. (**F**) Scaled density plot showing the C5Nef:C3Nef ratio for C3GN (red) or DDD (blue) Nef-positive C3G samples. (**G**) Hemolytic assay showing the convertase-stabilizing ability for all 12 monoclonal C3Nefs in the absence or presence of properdin. n = 3 technical replicates per C3Nef. (**H**) Same assay as in (**G**) but filtered for C3Nef-dominant samples (C3Nef ≥ C5Nef +10%) in C3G consensus cohort. (**I-J**) Jitter plot (left) with corresponding scaled density plots (right) showing C3 (**I**) and sC5b9 (**J**) levels for C3Nef-dominant C3G Consensus Cohort samples separated into DDD and C3GN subtypes. (**K**) Hemolytic assay showing the convertase-stabilizing ability for all 6 monoclonal C5Nefs in the absence or presence properdin. n = 3 technical replicates per C5Nef. (**L**) Same assay as (**K**) but filtered for C5Nef-dominant samples (C5Nef scores ≥ C3Nef + 10%) in C3G Consensus Cohort. Note the high representation of C3GN samples based upon C5Nef-dominant filtering. (**M-N**) Jitter plot (left) and corresponding scaled density plot (right) showing C3 (**M**) or sC5b9 (**N**) levels for those same C5Nef-dominant samples. Dotted red lines in (**G-N**) indicate a positive clinical result. For all panels: PBMC, peripheral blood mononuclear cells; UMI, unique molecular identifier; VH, variable heavy domain; VL, variable light domain; C3GN, C3 glomerulonephritis; DDD, dense deposit disease; FP, factor P/properdin.

### C3Nefs drive DDD, C5Nefs drive C3GN

C3G is subdivided by electron microscopy into dense deposit disease (DDD), characterized by highly electron-dense, ribbon-like deposits within the lamina densa of the glomerular basement membrane, or C3 glomerulonephritis (C3GN), which has amorphous, fluffy deposits in the mesangial and subendothelial cells. While previous studies have linked C3Nefs with DDD and C5Nefs with C3GN the basis for this association and the mechanisms driving each subtype remain unclear ^46,47^. To examine this relationship, we retrospectively analyzed two C3G patient cohorts: The first was a “C3G Consensus Cohort” consisting of a small but comprehensively vetted group of confirmed DDD or C3GN patients who had undergone biopsy-verified immunofluorescence and electron microscopy screening, and who had received extensive in-house biomarker and genetic testing. The second more expansive dataset, termed “C3G Large Cohort”, included within it the Consensus Cohort but with the addition of C3G patients who were given a DDD or C3GN diagnosis from their referring health care provider, and for whom biomarker data were available (fig. S34, A to B). For our analyses we treated each sample (single blood draw) independently, filtering on Nef status, properdin level, or diagnosed subtype.

Beginning with the Consensus Cohort, we first stratified C3Nef- or C5Nef-positive samples by their histologic subtype (DDD versus C3GN) and compared C5Nef:C3Nef ratios. Consistent with prior reports, most DDD samples displayed higher levels of C3Nefs, while C3GN samples trended toward higher levels of C5Nefs (Fig. 5F).

One highly informative insight afforded by monoclonal analysis was that in the convertase stabilization assay used for clinical Nef detection, all 12 C3Nefs produced a false-positive C5Nef readout (Fig. 5G). In the false-positive readout, we observed that the addition of properdin reduced the hemolytic score of monoclonal C3Nefs by ≈10% as a reflection of the competition between C3Nefs and properdin (Fig. 5G, fig. S3A). On this basis, we filtered Nef-positive C3G patient samples into two groups: samples for which C3Nef ≥ C5Nef + 10% (defined as “C3Nef-dominant”) and samples having C5Nef ≥ C3Nef + 10% (defined as “C5Nef-dominant”).

Strikingly, segregation of the Consensus Cohort samples by Nef dominance effectively separated clinical subtypes: the C3Nef-dominant group was composed predominantly of DDD samples that displayed moderately low C3 levels paired with normal levels of properdin, C5 and sC5b9, whereas the C5Nef-dominant group was overwhelmingly C3GN samples with severely reduced C3, low levels of C5 and properdin, and elevated sC5b9 (Fig. 5, G to N, fig. S35, A to D).

Examination of the C3G Large Cohort revealed that these DDD- and C3GN-specific biomarker signatures were only evident when examining Nef-positive samples (fig. S36, A to E), as the signatures were lost or weakened when the analysis was performed on Nef-negative C3G samples (fig. S36 F to J). Similar to the Consensus Cohort, filtering of the C3G Large Cohort based on Nef-dominance remained an effective way to segregate C3G subtypes, with the C5Nef-dominant-filter being particularly effective at capturing samples with C3GN phenotypes (fig. S36, K to T).

Previous reports have shown properdin to be low in C3GN patients, yet the mechanisms driving this pattern remain unclear ^48–51^. Given our finding that properdin inhibited C3Nef binding, but was required for C5Nef binding, we examined Nef-positive samples from both our Large Cohort and Consensus Cohort stratified by circulating properdin levels. Here, Nef-positive C3G samples with normal properdin levels displayed a disproportionately high number of DDD subtype samples with elevated C3Nefs (fig. S37, A and B). In contrast, Nef-positive C3G samples with low properdin were highly enriched for C3GN subtype and displayed C5Nef dominance (fig. S37, C and D). Collectively, these *ex vivo* clinical observations recapitulate our monoclonal findings: C3Nefs compete with properdin, limit C3 consumption, and have low sC5b-9 in circulation; in contrast C5Nefs consume properdin, deplete circulating C3, and elevate circulating sC5b-9.

## Discussion

A monoclonal Nef and its epitope have eluded scientists for decades. Here, we addressed these knowledge gaps by identifying multiple monoclonal C3Nefs and C5Nefs, thereby revealing a transient, conformation-dependent, bimolecular epitope for a C3Nef on the alternative pathway convertase. Using the Nefs as tools, we showed how C3Nefs and C5Nefs exploit two distinct convertase configurations to produce divergent functional outcomes. We also presented a cryo-EM structure of a C3bBb alternative pathway convertase bound by C3Nef that revealed an unexpected 180° rotation of the protease domain centered on the MIDAS region. We labeled this previously undocumented C3 convertase configuration Bb-up (Movie S1).

The conformational distinction between Bb-down and Bb-up provides mechanistic explanation for several observations related to both Nef binding and basic complement convertase regulation. First was the role of properdin: C5Nefs bound the Bb-down state and required properdin. In contrast, C3Nefs bound the Bb-up state in which Bb sterically clashes with properdin (fig. S38), and in which C3Nef and properdin compete for the same epitope on C345c (Fig. 3), thus explaining the mutual exclusivity between C3Nef and properdin binding, and between C3Nefs and C5Nefs binding to the same convertase. Second was the identification of a composite epitope for C3Nefs. We showed that C3Nefs have negligible binding to individual convertase components Bb and C3b, and that robust binding required convertase assembly and progression to reveal conformationally the bimolecular epitope recognized by C3Nefs, which bridges Bb and C345c. Lastly, the Bb-up configuration reconciles the effects of several documented pathogenic variants in factor H [FH(p.R53C), FH(p.Q81P)] and factor B [FB(p.K323E), FB(p.K323Q), FB(p.S367R), FB(p.D371G)] that are spatially remote in all current Bb-down models but are in close proximity in Bb-up, providing a structural basis explaining their shared phenotype of impaired regulator-mediated decay ^52–56^ (fig. S39A-D; Movie S2).

A critical implication of our findings is that the full conformational range of Bb rotation is greater than appreciated as prior structures captured the C3bBb convertase with stabilizing molecules such as SCIN and properdin. In contrast, our structure was obtained in the absence of both molecules, permitting visualization of a Bb-up state. Extending these findings to conformational dynamics and convertase control, we propose that complement regulators first engage the lower MG ring and position their upper SCR domains adjacent to C345c. In this position, the upper SCRs can capture a naturally rotating Bb and facilitate its permanent displacement by overextending its normal trajectory (fig. S39; Movie S2). By contrast, properdin appears to stabilize convertases by impinging on the natural rotation of Bb around the MIDAS axis rather than tethering Bb to C3b as currently believed (fig. S38; Movie S1). These concepts align with emerging data and offer a unifying framework for convertase progression, stabilization by properdin, and RCA-mediated inhibition ^40^.

The unique properties of C3Nefs and C5Nefs we presented reconcile overarching clinical biomarker patterns. C3Nef-stabilized convertases, although inactive when bound, have a significantly increased half-life and are protected from endogenous regulators compared to normal convertases. These C3Nef:C3bBb complexes adhere naturally to the glomerular endothelial glycocalyx, and upon Nef release, the activated convertase drives localized alternative and terminal pathway activity in the glomeruli ^57^. As a consequence, C3Nef positivity is associated with modest reductions in C3, local membrane-bound convertase stabilization, and phenotypes compatible with DDD. By contrast, C5Nef-bound convertases remain catalytically active in circulation and produce profound hypocomplementemia and elevated soluble C5b-9, as is seen with C3GN histology ^46,47,57,58^. Clinically, our data identified a previously unappreciated source of diagnostic error: C3Nef-dominant samples can yield false-positive C5Nef calls. We therefore propose that diagnostic workflows incorporate quantitative C3Nef/C5Nef ratios in conjunction with adjunct biomarkers to improve patient stratification and potentially guide selection of anti-complement therapies (fig. S40).

Beyond C3G, the approach described here demonstrates a new path for discovery of complement-modulating biologics. Conventional phage-display and biopanning strategies struggle with the extreme lability of C3bBb and attempt to rigidify the convertase with Ni²⁺, SCIN, or FB mutants. However, these techniques abrogate physiologically relevant conformations and as such are ineffective in capturing C3Nefs (fig. S18). By contrast, we used unbiased single cell sequencing to recover human-derived monoclonal antibodies that recognize convertase states that both inhibit (C3Nef) and potentiate (C5Nef) convertase function. These monoclonals provide valuable mechanistic probes and may show promise as tools for rational design of complement therapeutics that exploit convertase dynamics *in vivo*.

While these results represent a significant step forward in our understanding of C3G, Nef biology, and basic complementology, we acknowledge that we have mapped the conformational epitope for only one C3Nef clonotype and have not resolved the full repertoire for all monoclonals identified. Additional high-resolution structures and mutational mapping will be required to generalize the Bb-up/Bb-down paradigm across Nef classes, and this work is ongoing. Functional studies in animal models and longitudinal patient cohorts will be important to validate the proposed pathogenic mechanisms, and translating monoclonal Nefs into safe therapeutics will require careful engineering to avoid off-target complement effects.

The heterogeneity of nephritic factors has made it difficult to achieve both a consensus nomenclature and testing methodology. Here, for convention and readability, we intentionally chose the labels “C3Nef” and “C5Nef” in an effort to clearly distinguish the two separate classes. However, our data reveal inconsistencies in both labels given that the “C3Nefs” described herein actually prevent C3 cleavage and that the “C5Nefs”, while driving C5 activation in fluid phase, clearly target a properdin-stabilized C3 convertase and are poor activators of the membrane attack complex. In light of these findings, the alternatively used terminology of “properdin dependent” vs. “properdin independent”, paired with new modifiers of “convertase-pausing” vs. “convertase-activating”, might offer better mechanistic clarity into Nef classification. Additional studies and peer input will be necessary to reach nomenclature consensus moving forward.

In conclusion, the identification of the first *in vivo* monoclonal nephritic factors made it possible to capture previously undocumented physiologically relevant convertase conformations. This work reconciles clinical observations in complement biology, refines our mechanistic understanding of convertase regulation, and opens a practical path toward new diagnostics and complement-modulating immunotherapeutics that leverage native convertase dynamics.

## Materials and Methods

### Study population

All patient samples were obtained between January 1^st^, 2016, and July 1^st^, 2025. All complement biomarker assays were performed in house by the Molecular Otolaryngology and Renal Research Laboratories (MORL), University of Iowa. For analyses, patients were divided into two overlapping cohorts. The first was the “C3G Large Cohort” that consisted of 3,097 unique samples obtained from 1,263 C3G patients. Nested within this dataset was a second group, termed “C3G Consensus Cohort”, that consisted of 1,407 samples from 252 C3G patients with biopsy-proven C3G selected from our highly curated C3G registry. Inclusion in the C3G Consensus Cohort was based on the availability of sufficient sera and plasma samples to complete all assays multiple times, as well as complete histopathologic data (light microscopy, immunofluorescence, and EM) required to confirm a C3G diagnosis. All C3G Consensus Cohort patients had biopsy-proven disease consistent with the description provided in the C3G consensus article ^11^, and all qualifying biopsies were confirmed by the research team as well as an independent pathologist. For each analysis, samples were filtered by C3Nef or C5Nef status, circulating properdin levels, or DDD/C3GN diagnosis as documented by their primary health care provider (for C3G Large Cohort), or as verified by MORL (for C3G Consensus Cohort).

C3G Consensus Cohort samples obtained while patients were receiving eculizumab, ravulizumab, pegcetacoplan, iptacopan were excluded from all analyses. The majority of non-C3G Consensus Cohort samples that constitute the bulk of the C3G Large Cohort dataset were provider-diagnosed and were not filtered to remove genetic drivers, drug treatment, or C4Nefs status. The University of Iowa Institutional Review Board approved all procedures, and all patients gave informed consent before donating samples.

### Peripheral blood mononuclear cell sorting

Blood was obtained with consent from the C3G patient. PBMCs were isolated from whole blood using Cytiva Ficoll-Paque PLUS per manufacturer’s instructions, resuspended at ∼20×10^6^ cells/mL in cold 90% heat-inactivated FBS + 10% DMSO, and frozen in liquid nitrogen until needed. Upon use, cells were thawed and washed 2x in 10mL warm RPMI buffer + 10% heat-inactivated FBS. Cells were then resuspended in 100µl of 1x Annexin V binding buffer (Invitrogen; 00-0055-56), stained as listed in table S1 for 20 min at 4°C in the dark. Following staining, cells were washed 2x in 1mL and resuspended in 1-2mL of Annexin V buffer +1% heat-inactivated FBS and then sorted. Antigen-experienced B cells were isolated by FACS purification using a Cytek Aurora CS. B-lineage cells were identified by gating on Live Cells > Singlets > Dump^−^ (dump markers: CD2^+^ CD3^+^ CD14^+^ CD16^+^ CD36^+^) > CD19^+^ and CD38^+^ cells.

Specific B-lineage subpopulations were sorted into two groups for 10X Chip K loading: Group 1 consisted entirely of memory B cells (CD19+CD38+/IgD^−^/CD27^+^CD21^+^); and Group 2 “Pooled” was a combination of circulating plasma cells/plasmablasts (CD19+CD38+/CD27^Hi^CD38^Hi^), marginal zone-like cells (CD19+CD38+/IgD^+^CD27^+^), and atypical memory B cells (CD19+CD38+/IgD^−^CD21^−^). All cells were kept on ice, sorted and collected at 4°C. Sorted cells were used immediately for 10X processing. All FACS sorting and analysis were performed at the University of Iowa Flow Cytometry Core.

### Library preparation and sequencing

After FACS sorting, B cells were washed, counted, and resuspended in cold PBS +0.01% BSA for immediate single cell processing using the following kits from 10X Genomics: (1) Chromium Next Gem Single Cell 5’ Reagent Kit v2 (Dual Index) PN-1000263; (2) Chromium Single Cell Human BCR Amplification Kit PN-1000253; (3) Chromium Next GEM Chip K Single Cell Kit PN-1000286; (4) Library Construction Kit PN-1000190; (5) Dual Index Kit TT Set A PN-1000215. Briefly, following the protocols outlined by 10X in the CG000331 Rev E manual, sorted B cells were processed to generate gel-beads-in-emulsion, barcoded cDNA, BCR-amplified VDJ libraries, and gene expression libraries. When possible, lanes were loaded with the maximum recommended number of cells per lane. Following library construction, BCR V(D)J libraries were pooled and sequenced on an Illumina NovaSeq 6000 on a SP flow-cell with a run length of 100 cycles. Raw BCL files from the NovaSeq were demultiplexed and converted to FASTQ files using Illumina bcl2fastq2 version 2.20. The FASTQ files were processed by the Cell Ranger version 7.1.0 V(D)J pipeline to assemble sequences based on the GRCh38 V(D)J reference.

### Clone selection for initial screening

Following single-cell sequencing, clonotypes were generated using enclone ^29^. Single clone sequences were selected from candidate clonotypes for monoclonal production and functional screening. Based on Nef literature ^24,25,59–65^ we chose several criteria (discussed below) to guide our filtering and clonotype selection. These literature-informed benchmarks were not mutually exclusive and selected clonotypes/clones fulfilled most criteria.

#### Presence in PBMCs

While long-lived plasma cells take up residence in bone marrow and intestinal niches, we hypothesized that in Nef-positive C3G patients some fraction of Nef-producing plasma cells, short-lived plasmablasts, or B cells circulate as PBMCs ^59^.

#### Isotype

The majority of clinical Nef-based assays include an IgG isolation or enrichment step prior to Nef testing. Thus, we were confident that Nefs exist in the IgG fraction; more specifically, Nefs are of the IgG3 subtype ^24,25,60–62,65^.

#### Secretion

By definition, Nefs are circulating autoantibodies. Therefore, we prioritized BCR sequences displaying distinctly high transcript levels (as measured by unique molecular identifier count), consistent with active antibody expression/section.

#### Origin

We prioritized sequences from our “pooled” fraction as antibody-secreting plasma cells/plasmablasts and atypical B cell populations are commonly associated with autoimmune disease ^63^.

#### Persistence

For many patients, Nefs persist across time. Therefore, we prioritized sequences that appeared in both timepoints.

#### Uniqueness

In examining complementarity-determining regions (CDRs), we prioritized CDR3 sequences that appeared in our C3G Nef+ patient only and not in controls or public single-cell BCR datasets. For example, we excluded sequences with CDR3s similar to those found in COVID datasets ^64^.

#### Increased Cellular Frequency

Nefs target C3 convertase. With continual antigenic exposure *in vivo*, we expected Nef-producing B cell proliferation and therefore favored sequences found within larger clonotypes (i.e., larger clonotype clusters containing a greater number of B cells). *Correlations to Clinical Data*: Nef levels change by clinical testing across time. Thus, we prioritized clonotypes displaying cell numbers that correlated with the fluctuations observed in the C3G patient’s clinical Nef scores. i.e., we prioritized clonotypes with higher cell counts in timepoint one, in which polyclonal Nef scores were highest.

### C3Nef detection

Convertase stabilization by Nefs was detected using hemolytic-based assays (C3 convertase stabilizing assay, and C3 convertase stabilizing assay with properdin to detect C3Nefs and C5Nefs, respectively) as previously described ^66^. Briefly, to detect C3Nefs, C3 convertase was built on the surface of C3b-decorated sheep erythrocytes using purified FB and factor D. The amount of FB was titrated to yield approximately 1 convertase per erythrocyte at 30⁰C after 5 minutes, at which time patient-purified IgG was added. Twenty minutes were allowed for natural decay of the convertase; rat serum in ethylenediamine tetraacetic acid was then added to supply terminal complement components. The addition of rat serum leads to cell lysis in the presence of Nef-stabilized C3 convertase. C3 convertase stabilizing assay with properdin included properdin to facilitate C5 convertase formation on the erythrocyte, with a longer incubation period to accommodate the longer C5 convertase half-life. Nef stabilization capacity was measured as a percentage of hemolysis relative to a negative control. Clinical cutoff for a positive assay was >20% hemolysis.

### Amine-coupled C3 convertase specificity assay

C3b was amine coupled on Flow cell 2 of a CM5 sensor chip to 1206 RUs (target RU= 1200) and docked into a Biacore X100 SPR machine. Monoclonal antibodies and polyclonal patient samples were tested in HBS-EP supplemented with 10mM MgCl_2_. Analyte conditions were as follows: Buffer +/- IgG, FD (50nM) +/- IgG, FB (500nM) +/- IgG, and FB (500nM) + FD (50nM) +/- IgG for a total of eight analyte conditions. Monoclonal IgG-positive samples were tested at 100nM, and polyclonal IgG-positive samples were tested at 3uM. For each cycle, the analyte was injected for 120 seconds, followed by 120 seconds of dissociation, and 90s of regeneration (1M NaCl, 10mM Na-acetate, pH 4.0). Binding was recorded at 20 seconds post-injection, and ΔRU was calculated using the following formula ^67^:

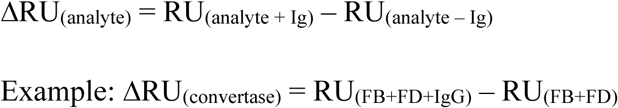

Each antibody sample was run in triplicate and is shown with an average cycle.

### Amine-coupled avidity kinetics assay

C3b was amine coupled to Flow cell 2 of a CM5 sensor chip to 1222 RUs (target RU= 1200) and docked into a Biacore X100 machine with temperature adjusted to 37° Celsius. FB (500nM), FD (50nM), and monoclonal (200nM) or polyclonal (3uM) IgG were injected in HBS-EP supplemented with 10mM MgCl_2_ for 543 seconds at 10 uL/min, then permitted 6000 seconds of dissociation before regenerating with 1M NaCl, 10mM Na-acetate, pH 4.0 (or pH 3.6 as needed) for 120 seconds. Data were referenced to reagent convertase cycles (Ig-negative control cycles) at point A to verify reagent convertase half-life of 90 seconds before normalizing the sensorgram at 600 seconds post-injection (report point “B”) to 100 normalized RUs. The estimated half-life was calculated as:

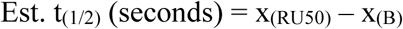

such that
x_(RU50)_ = average time x with domain RUx with the range [49.9, 50.1]

Each antibody sample was run in triplicate and shown with an average cycle.

### Protein G capture specificity assay

A Sensor Chip Protein G (Cytiva Product # 29179316) was loaded into a Biacore X100 SPR machine. Monoclonal Ig (150nM) samples in HBS-EP supplemented with 10mM MgCl_2_ were captured on Flow cell 2 for 180 seconds at 5uL/min. Flow cell 1 was left as a reference.

Complement alternative pathway components were injected over the Ig captured chip in sequential cycles with analyte conditions as follows:

1. Bb (500nM)
2. FB (500nM)
3. C3b (500nM)
4. FP (500nM)
5. Proconvertase [C3b (200nM) + FB (500nM)]
6. FP-Proconvertase [C3b (200nM) + FB (500nM) + FP (150nM)]
7. Convertase [C3b (200nM) + FB (500nM) + FD (200nM)]
8. FP-Convertase [C3b (200nM) + FB (500nM) + FD (200nM) + FP (150nM)]
9. FP-C3b [C3b (200nM) + FP (150nM)]

Due to the labile nature of the proconvertase and convertase complexes, precise timing was required in analyte preparation. All convertase and proconvertase analyte conditions were prepared with the following considerations: 1) FB was added ∼500 seconds before injection, 2) C3b was added 325 seconds before injection, 3) the sample was mixed gently by pipetting then incubated at room temperature until injection. Analytes were injected for 60 seconds followed by 60 seconds of dissociation at 10 uL/min. Analyte was removed by regeneration with 1M NaCl, 10nM Na-acetate, pH 4.0 for 30 seconds. The final regeneration (to remove captured antibody) after all analyte conditions were evaluated was accomplished by injecting 10mM Glycine-HCl pH 1.5 for 30 seconds. Binding was recorded 20 seconds post-injection and is presented in reference subtracted sensorgrams. Each monoclonal antibody was tested in duplicate.

### RCA effect on C3Nef-stabilized convertases (C3bBb)

C3b (Complement Technology) was biotinylated at the TED domain using the EZ-Link^TM^ Maleimide-PEG2-biotin kit (ThermoFisherScientific) according to the manufacturer’s instructions, at a molar ratio of 1:20 of C3b to biotin. Subsequent experiments were conducted in a Biacore X100 at 20 °C, using HBS-P+ containing 1 mM MgCl_2_ as the running buffer at a flow rate of 10 µl/min. The SA chip was conditioned according to the manufacturer’s instructions.

Biotinylated C3b was immobilized by injection until a final amount of 450-550 RU was reached. During each cycle, convertases formation was induced by injecting 500 nM FB and 50 nM FD (Complement Technology) for 180 s. Next, 100 nM of C3Nefs or buffer was injected for 2 min followed by a 2 min dissociation phase. 1 µM of a regulator [FH(1-6), sCR1(1-3) or sDAF(1-4)] was injected for 3 min. The chip was regenerated by injecting 100 mM acetate with 1 M NaCl at pH 3.6 twice for 30 s. To facilitate the comparison of data from multiple cycles in a tabular format, convertase/Nef levels were normalized (set to 100%) against the observed response levels prior to regulator injection. Convertase decay was measured by determining and expressing the remaining response levels at 1125 s of each cycle as a percentage of the 1125 s set point. All injections were performed in duplicate.

### RCA effects on C5Nef stabilized convertases (C3bBbFP)

C3b (Complement Technology) was tagged at the TED domain using an ALFA-tag peptide with N-terminal maleimide (5’-maleimide-GGSRLEEELRRRLTE-3’, synthesized by GenScript).

C3b was incubated with the ALFA-tag peptide at a 1:25 molar ratio for 2 h on ice and separated from remaining peptide by using size-exclusion chromatography (Cytiva Superdex™ 200 Increase 10/300 GL). Subsequent experiments were conducted in a Biacore X100 (Cytiva) at 20 °C, using HBS-P+ containing 1 mM MgCl_2_ as the running buffer at a flow rate of 10 µl/min. The SA chip was conditioned according to the manufacturer’s instructions. Both flow cells were equally saturated with 10 µg/ml SdAb anti-ALFA (NanoTag Biotechnologies; Cat: N1505) by injection to each flow cell separately. During each cycle, C3b-ALFA was captured to reach a response of approx. 1000 RU (in the first cycle) on one flow cell by injecting it for 180s. Then 200 nM properdin (Comptech) was injected on both flow cells for 2 min, followed by convertases formation with 500 nM FB and 50 nM FD (Complement Technology) for 3 min, 100 nM C5Nef injection for 2 min and RCA [FH(1-6), sCR1(1-3) or sDAF] injection for 3 min. The chip was regenerated by injecting 4 M Gdn*HCl with 2 M MgCl_2_, pH 1.9 for 30 s. To facilitate the comparison of data from multiple cycles, the C3b-ALFA levels were normalized (set to 100%) and convertase decay was measured by determining the remaining signal at the report point 2500 s of each cycle. All injections were performed in duplicate.

### Protein A C3 activation/inhibition assay

A Sensor Chip Protein A (Cytiva Product # 29127557) was loaded into a Biacore X100 SPR machine. Sample conditions were tested iteratively using the following framework:

1. Monoclonal Ig (200nM) in 10mM MgCl_2_ HBS-EP running buffer was captured on Flow cell 2 for 180 seconds at 5 uL/min. Flow cell 1 was left blank as a reference.
2. Samples were tested via three sequential analyte injections at 10uL/min: Injection 1: (60 seconds): buffer (control) or convertase (test sample) Injection 2: (120 seconds) + 60 seconds dissociation: buffer (control) or C3 (test sample) Injection 3: (90 seconds) + 60 seconds dissociation: FB with FD
3. Sensor chips were regenerated with a 30 second injection of 10mM Glycine-HCl pH 1.5.

The background binding of C3 to the monoclonal antibody ligand was established first. In these cycles, Injection 1= Buffer and Injection 2= C3 (1uM). To evaluate for the impact of drug, C3 was injected alone or supplemented with 16uM CP40 or with 30uM iptacopan.

The Nef-captured AP convertase was analyzed for the ability to activate C3. In these cycles, Injection 1= Convertase and was prepared with FD (200nM), FB (500 nM) and (for C5Nefs) monomeric FP (500nM). At 325 seconds before sample injection, C3b (200nM) was added and incubated at room temperature. To evaluate the impact of drug, convertase was injected alone or supplemented with 16uM CP40 or with 30uM iptacopan. A reference curve for the association and decay of each convertase condition was established by Injection 2= Buffer. To evaluate the captured Convertase-IgG complex for the ability to activate C3, the convertase injection was followed by the 1uM C3. Iptacopan and CP40 convertase/C3 analytes were tested as a pair (i.e. CP40-Convertase analyte and CP40-C3 analyte were sequentially injected in one cycle). Activated and deposited nascent C3b on the chip surface was revealed with Injection 3= FB (500nM) with FD (100nM).

Each background C3 binding sensorgram (Inj 1= buffer; Inj 2= C3) was subtracted from the corresponding convertase test cycle (Inj 1= Convertase; Inj 2= C3), and this residual subtracted sensorgram was normalized to 100RU at 20 seconds post-injection. The corresponding convertase reference cycle (Inj 1= Convertase; Inj 2= Buffer) was also normalized to 100RU. Convertase activity was determined by the formation of convertase during the FB+FD injection and the mechanism of inhibition modeled by comparing the C3 injection between conditions (reagent vs CP40 vs iptacopan).

### Properdin and SCIN dose-response assays

A Sensor Chip Protein G (Cytiva Product # 29179316) was loaded into a Biacore X100 SPR machine. Monoclonal Ig (200nM) in HBS-EP with 10mM MgCl_2_ running buffer was captured on Flow cell 2 for 180 seconds at 5uL/min. Flow cell 1 was left blank as a reference.

The SCIN titration utilized the monomeric chimera SCIN protein “SCIN-ChC3b2” ^34^ and was prepared with a 10x dilution factor from 5000nM SCIN to 0.5nM SCIN and a 0nM SCIN control in running buffer. Each SCIN dose was added to FD (200nM) and FB (500nM) and (for C5Nefs) monomeric FP (200nM). C3b (200nM) was added to each sample and incubated for 325 seconds at room temperature then injected on the captured monoclonal antibody.

The FP titration was prepared with a 5x dilution factor from 3750nM FP to 1.2nM FP and a 0nM FP control in running buffer. Each FP dose was added to FD (200nM) and FB (500nM). C3b (20nM) was added to each sample and incubated for 325 seconds then injected on the captured monoclonal antibody.

Analytes were injected for 60 seconds followed by 60 seconds of dissociation at 10 uL/min. Analyte was removed by regeneration with 1M NaCl, 10mM Na-acetate, pH 4.0 for 30 seconds. The final regeneration of the Protein G chip was accomplished by injecting 10mM Glycine-HCl pH 1.5 for 30 seconds. Binding was measured 20 seconds post-injection and is presented as dose-response curves and in reference subtracted sensorgrams.

### Alternative pathway activity on guinea pig red blood cells

Fresh guinea pig erythrocytes (GP-E) in alservers solution (#30100 - Colorado Serum Company, Colorado, United States) were washed 3 times in veronal buffer solution (VBS) (3.13mM barbital C-IV) supplemented with 10mM ethylene glycol-bis(2-aminoethylether)-N,N,N′,N′-tetraacetic acid (EGTA) (AP-VBS) to prevent the activation of the classical pathway. Alternative pathway lytic activity of a Normal Human Serum (NHS) (Innovative research, Inc., #39490) was titrated by serial dilutions starting from 20% in 100µL reactions containing 1×10^8 GP-E/mL. Mixtures were incubated for 30 min at 37°C in a water bath and reactions were stopped with 100µL of ice-cold VBS containing 20mM ethylenediaminetetraacetic acid (EDTA) (VBS-EDTA). Mixtures were spun at 2000rpm for 5min and 120µL supernatant were transferred to a flat-bottom microtiter plate to measure absorbance at 415nm in a Tecan Infinite M plex (#30190085 - Tecan Group Ltd., Männedorf, Switzerland). The minimum amount of NHS with 100% lytic capacity was determined and fixed for the rest of the assays. Reactions were replicated with the fixed amount of NHS, and C3Nefs and C5Nefs impact on lytic activity was titrated by serial dilutions starting from 150µg/mL.

### Fluid phase activity

A modified immunofixation electrophoresis (IFE) was performed to assess fluid-phase C3 convertase activity by mixing increasing concentrations of monoclonal IgG (Nefs) with normal human serum. Following a 25-minute incubation period at 37 ⁰C and immunoprecipitation by an anti-C3 antibodies, C3 convertase activity was measured by quantifying the generation of C3 activation products by agarose gel electrophoresis using a SPIFE Touch per manufacturer’s instructions (Helena Laboratories, Inc).

### Mass photometry

MP experiments were performed on a Refeyn Two^MP^ mass photometer (Refeyn Ltd, Oxford, UK). Microscope coverslips (24 mm x 50 mm, Thorlabs Inc.) were cleaned by serial rinsing with Milli-Q water and HPLC-grade isopropanol (Sigma Aldrich) followed by drying with a filtered air stream. Silicon gaskets (Grace Bio-Labs) to hold the sample drops were cleaned in the same procedure immediately prior to measurement. All MP measurements except those using Ni^2+^ were performed at room temperature in a buffer (25 mM HEPES pH7.5, 150 mM NaCl and 5 mM MgCl_2_). The instrument was calibrated using a protein standard mixture: β-amylase (Sigma-Aldrich, 56, 112 and 224 kDa) and thyroglobulin (Sigma-Aldrich, 670 kDa). Before each measurement, 15 µL of buffer was placed in the well to find focus. The focus position was searched and locked using the default droplet-dilution autofocus function after which 5 µL sample was added and pipetted up and down to briefly mix before movie acquisition was promptly started. Movies were acquired for 60 s (3000 frames) using AcquireMP (Refeyn Ltd) using standard settings. All movies were processed, analyzed using DiscoverMP (Refeyn Ltd).

### Statistics and data visualization

Statistics and data visualization were completed using GraphPad Prism version10.6.1, R version 4.4.1 ^68^, RStudio version 2024.04.2+764^69^. Statistical significance was evaluated using a Student’s t-test or an ordinary one-way analysis of variance, as indicated.

### Preparation of purified proteins

Human C3b (catalog # A114), FB (catalog # A135) and FD (catalog # A136) were purchased from Complement Technology. C3Nef.9 was obtained from TWIST Bioscience, Inc. SCIN proteins were expressed in E. coli BL21 (DE3) and purified from inclusion bodies by refolding on Ni-NTA column and followed by Superdex 30 size exclusion chromatography.

### Cryo-EM sample preparation

The complex of C3 convertase (C3bBb) with C3Nef.9 was formed by mixing 1 uM C3b and 1 uM FB in a buffer (25 mM HEPES pH7.5, 150 mM NaCl and 5 mM MgCl_2_) and incubating at room temperature for 10 mins, adding 0.5 uM FD for 1 min and then adding 1 uM C3Nef.9 for another 10 mins. A volume of 3 uL of the complex mixture was applied immediately to a Quantifoil copper grid (R2/1 300 mesh), which was freshly glow-discharged at 15 mA for 60 s using PELCO easiGlow (Ted Pella). The grids were blotted at 4 °C and 95% humidity using a Vitrobot Mark IV (Thermo Fisher) for 3.5 or 4 s with −5 blot force after 10 s waiting time. The grids were then plunge-frozen into liquid ethane and stored in liquid nitrogen.

### Cryo-EM data acquisition

Two cryo-EM datasets were collected for the complex of C3Nef.9:C3bBb. Dataset 1 (no tilt) was collected on the 300 kV Titan Krios microscope (Thermo Fisher) equipped with a CFEG, Falcon 4i detector and Selectris energy filter with a slit width of 10 eV. The micrographs were acquired in EER format at a magnification of 165,000x (pixel size of 0.74 Å) with a defocus range of −2.4 to −0.9 um. Dataset 2 (a tilt angle of 30°) was collected on the 300 kV Titan Krios microscope (Thermo Fisher) equipped with a XFEG, Gatan K3 detector and Bioquantum energy filter with a slit width of 20 eV. The micrographs were acquired in a super-resolution mode (pixel size of 0.413 Å) at 105,000x corresponding to a physical pixel size of 0.826 Å with a defocus range of - 2.2 to −1.0 um. Each movie stack contains 50 frames with a total dose of 50 e^-^ Å ^-2^. 11,128 movies (dataset 1) and 6,301 movies (dataset 2) were collected.

### Cryo-EM image processing

Movies were motion corrected and CTF estimated using patch motion correction and patch CTF estimation in CryoSPARC V4.6.2. The movies collected in super-resolution mode were binned 2x during motion correction. The exposures with CTF resolution fit above 8 Å, thick ice or target on carbon support were removed before further processing. The particles were picked using a blob picker. After particle inspection and 3-4 rounds of 2D classification, the classes with high-resolution features were selected as references for template picking. Another 3-4 rounds of 2D classification were performed to remove “junk” particles. Two datasets were processed separately then the picked particles from the non-tilt dataset were extracted with a 672-pixel box down sampled to 602 pixel (0.826 Å/pixel). The combined particles were subjected to another round of 2D classification and used to generate ab initio models and 3 class hetero refinement.

The class containing 157,414 particles, which had the feature of C3 beta chain MG ring selected for CTF refinement and Non-uniform refinement, yielding a 3.69 Å reconstruction of the complex of C3Nef.9:C3bBb. To improve resolution of the core region of the complex, the two classes containing 462,324 particles were pooled together for CTF refinement and local refinement with a static mask, achieving a final resolution of the complex at 2.99 Å.

### Model building and refinement

Models of C3b and Bb were obtained from the crystal structure of C3bBb-SCIN complex (PDB: 2WIN). Models of C3Nef.9 were predicted by AlphaFold3. The individual structures were rigid body fitted into the density map in UCSF Chimera. The model was refined manually in Coot and refined further by ISOLDE in ChimeraX. The final model was real space refined in Phenix using phenix.real_space_refine. Figures were generated with UCSF Chimera and ChimeraX. All software was compiled in SBGrid.

## Acknowledgements

We thank Dr. Roberta Pelanda and Dr. Raul Torres from the Department of Immunology & Microbiology at the University of Colorado Anschutz School of Medicine for their kind input regarding B cell gating strategies and single-cell BCR sequencing. We also thank Prof. Dr. Nils Johnsson and the Institute of Molecular Genetics and Cell Biology of the University Ulm for allowing us use of their Biacore X100. We extend our thanks to Dr. Zhe (James) Chen and Dr. Yang Li from University of Texas Southwestern Medical Center for cryoEM data collection. CryoEM studies at the Structural Biology Laboratory and the Cryo-Electron Microscopy Facility at UT Southwestern Medical Center were partially supported by grant RP220582 from the Cancer Prevention & Research Institute of Texas (CPRIT) for cryo-EM studies.

## Funding

This work was funded in part by the following grants:

National Institutes of Health grant R01 DK110023

Deutsche Forschungsgemeinschaft (SCHM3018/4-1, project number 495891565 to CQS

## Author contributions

Conceptualization: SJW, YZ, CMN, RJHS

Data curation: SJW, CTC, ZX, HM, JS, DW, NJS

Formal analysis: SJW, CTC, ZX, HM, JS, DW, NJS

Funding acquisition: CQS, YZ, CMN, RJHS

Investigation: SJW, CTC, ZX, HM, JS, RG, CD, KK, SR, NCM, AFMN, ARC, SSJ, SNC, MDH, RF, TL, JH, LOF

Methodology: SJW, CTC, ZX, HM, JS, DW, NJS, CQS, YZ, RJHS

Project administration: SJW, MH, RF, YZ, CMN, RJHS

Resources: NJS, CQS, YZ, CMN, RJHS

Supervision: RJHS

Validation: SJW, CTC, ZX, HM, JS, NJS

Visualization: SJW, CTC, ZX, HM, JS, NJS

Writing – original draft: SJW, RJHS

Writing – review & editing: SJW, CTC, ZX, HM, JS, NJS, CQS, YZ, CMN, RJHS

## Declaration of interests

CQS is a named inventor of patent applications that describe the use of complement inhibitors for therapeutic applications. CQS has received research honoraria for speaking at symposia organized by Alexion Pharmaceuticals, Sobi and Sanofi, and serving on an advisory committee for Sobi. RJHS Directs the Molecular Otolaryngology and Renal Research Laboratories and serves on the Advisory Board for Novartis. CMN is the Associate Director of Molecular Otolaryngology and Renal Research Laboratories; serves on advisory boards for Novartis, Apellis, Vertex, and Alexion; participates as a site investigator for Novartis, Apellis, Biocryst, and Retrophin; is a member of the data safety monitoring board for Kira; serves as Chair of a data safety monitoring board for FIT4KID; and receives author royalties for UpToDate. RJHS, CMN, YZ, CTC, and SJW declare that a patent application related to some of the findings presented in this manuscript is currently being prepared. All other authors declare they have no competing interest.

## Data and materials availability

Requests for resources or reagents can be directed to the corresponding authors. All cryo-EM datasets are publicly available in RCSB PDB and EMDB with PDB accession numbers 10OM and 10OL and EMDB accession numbers 75336, 75335, respectively. All sequencing data generated or analyzed during this study are included in this manuscript and are available at the Gene Expression Omnibus^70,71^ series accession number GSE329746. This paper does not report original code.

## List of Supplementary Materials

**Materials and Methods**

**Fig. S1.** Complement profile of index C3G patient.

**Fig. S2.** Obtaining and screening C3Nef candidates.

**Fig. S3.** Convertase stabilizing properties of C3Nefs and C5Nefs.

**Fig. S4.** Workflow and diagram of Surface Plasmon Resonance (SPR) experiments to determine C3Nef reactivity to C3bBb convertase.

**Fig. S5.** C3Nefs bind the C3bBb convertase.

**Fig. S6.** Methodology for determining C3Nef estimated half-life values.

**Fig. S7.** Estimated half-life calculation: Step 1.

**Fig. S8.** Estimated half-life binding: Steps 2.

**Fig. S9.** Estimated half-life binding: Steps 3-5.

**Fig. S10.** Workflow and diagram of surface plasmon resonance (SPR) experiments used to identify C3Nef-mediated inhibition of RCA-accelerated convertase decay.

**Fig. S11.** C3Nefs inhibit the decay acceleration activity of regulators of complement activity.

**Fig. S12.** Workflow and diagram of Protein G-coupled surface plasmon resonance (SPR) experiments used to characterize C3Nef binding responses to C3 convertase and convertase components.

**Fig. S13.** C3Nef binding characterization by SPR.

**Fig. S14.** C5Nef binding characterization by SPR.

**Fig. S15.** C3Nefs are distinct from C5Nefs.

**Fig. S16.** Workflow and diagram of surface plasmon resonance (SPR) experiments used to characterize C5Nef inhibition of RCA accelerated convertase decay.

**Fig. S17.** C5Nefs inhibit the decay acceleration activity of regulators of complement activity.

**Fig. S18.** C3Nef binding to C3bBb is hindered by convertase-stabilizing modifications.

**Fig. S19.** Cryo-EM reconstruction of the C3Nef.9-stabilized C3bBb complex.

**Fig. S20.** Structural analysis the C3Nef.9-stabilized C3bBb complex.

**Fig. S21.** Variable light chain of C3Nef drives the interaction with factor B.

**Fig. S22.** C3bBb contact residues are conserved across all 12 C3Nefs.

**Fig. S23.** C3Nef and properdin binding sites overlap.

**Fig. S24.** C345c and Bb orientations in Bb-up compared to known models (Bb-down).

**Fig. S25.** Bb-up configuration reduces contact between Bb and C345c domain.

**Fig. S26.** Structural models suggest C3Nef-mediated inhibition of C3bBb.

**Fig. S27.** Workflow and diagram of surface plasmon resonance (SPR) experiments used to characterize binding to, or cleavage of, free C3 by C3Nef-stabilized C3bBb convertase.

**Fig. S28.** C3Nef-bound C3bBb convertase binds but does not cleave free C3.

**Fig. S29.** Workflow and diagram of surface plasmon resonance (SPR) experiments used to characterize cleavage of C3 by C5Nef-bound C3bBbP(mono) convertase.

**Fig. S30.** C5Nef-bound C3bBbP(mono) convertases cleave C3.

**Fig. S31.** Guinea pig red blood cell lysis is inhibited by C3Nefs but not C5Nefs.

**Fig. S32.** Effects of concentration and time on C3Nef activity.

**Fig. S33.** C3Nef estimated half-life and Nef effects on properdin levels.

**Fig. S34.** C3G patient datasets.

**Fig. S35.** Filtering on Nef dominance separates DDD from C3GN samples.

**Fig. S36.** Biomarker signatures differ between DDD and C3GN patients.

**Fig. S37.** Effect of properdin levels on Nef-dominance in DDD vs. C3GN patients.

**Fig. S38.** Properdin clashes with Bb-up and sterically hinders rotation around the MIDAS site.

**Fig. S39.** Bb-up rotational model of C3bBb decay.

**Fig. S40.** Comparison of C3Nefs to C5Nefs.

**Table S1.** Reagents used for FACS enrichment of B lineage PBMCs.

**Table S2.** C3Nef and C5Nef V(D)J sequence information; heavy chain.

**Table S3.** C3Nef and C5Nef V(D)J sequence information; light chain.

**Table S4.** CryoEM data collection and processing.

**Table S5.** CryoEM model refinement and validation statistics.

**Table S6.** Analysis of the interfaces between C3bBb convertase and C3Nef.

**Supplemental References (55-65)**

**Movie S1.** Hypothetical Bb-rotation model of properdin-mediated C3bBb stabilization and C3Nef binding.

**Movie S2.** Hypothetical Bb-rotation model of RCA-mediated C3bBb decay.

## Supplementary Materials

**The PDF file includes:**

Figs. S1 to S40

Tables S1 to S6

References (55-65)

**Other Supplementary Materials for this manuscript include the following:**

Movies S1 to S2

**Fig. S1.**
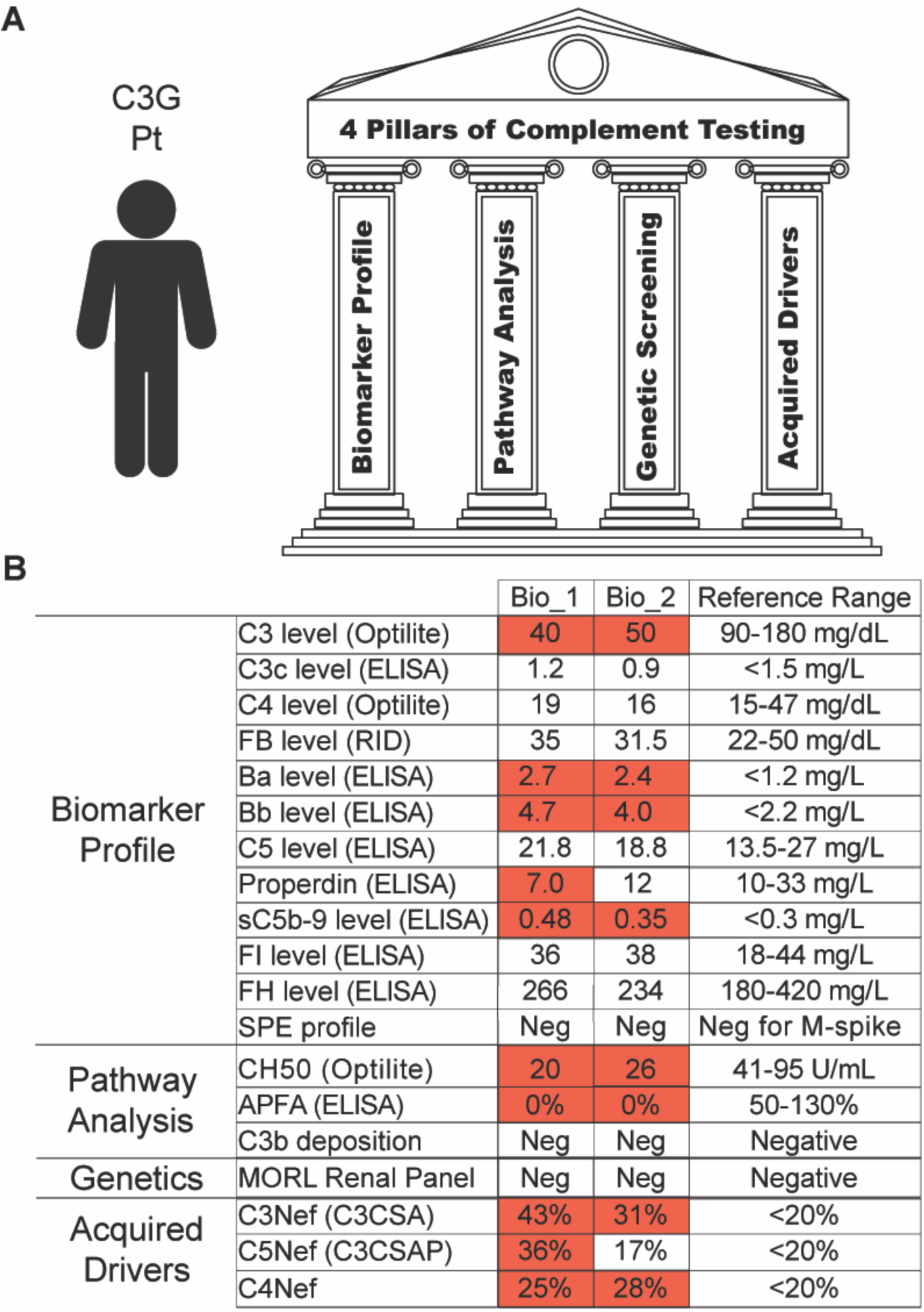
Complement profile of index C3G patient. (**A**) The four pillars of complement testing: complete complement biomarker profile, functional pathway analysis, genetic driver screening, and acquired driver screening. (**B**) Results of 4-pillar complement testing from two blood samples acquired 7 months apart (Bio_1 and Bio_2) from C3G index patient. Normal reference ranges are listed on the right. Abnormal values are highlighted in red. FB, factor B; RID, radial immunodiffusion; FI, factor I; FH, factor H; SPE, serum protein electrophoresis; APFA, alternative pathway functional assay; C3CSA, C3 convertase-stabilizing assay; C3CSAP, C3 convertase-stabilizing assay with properdin.

**Fig. S2.**
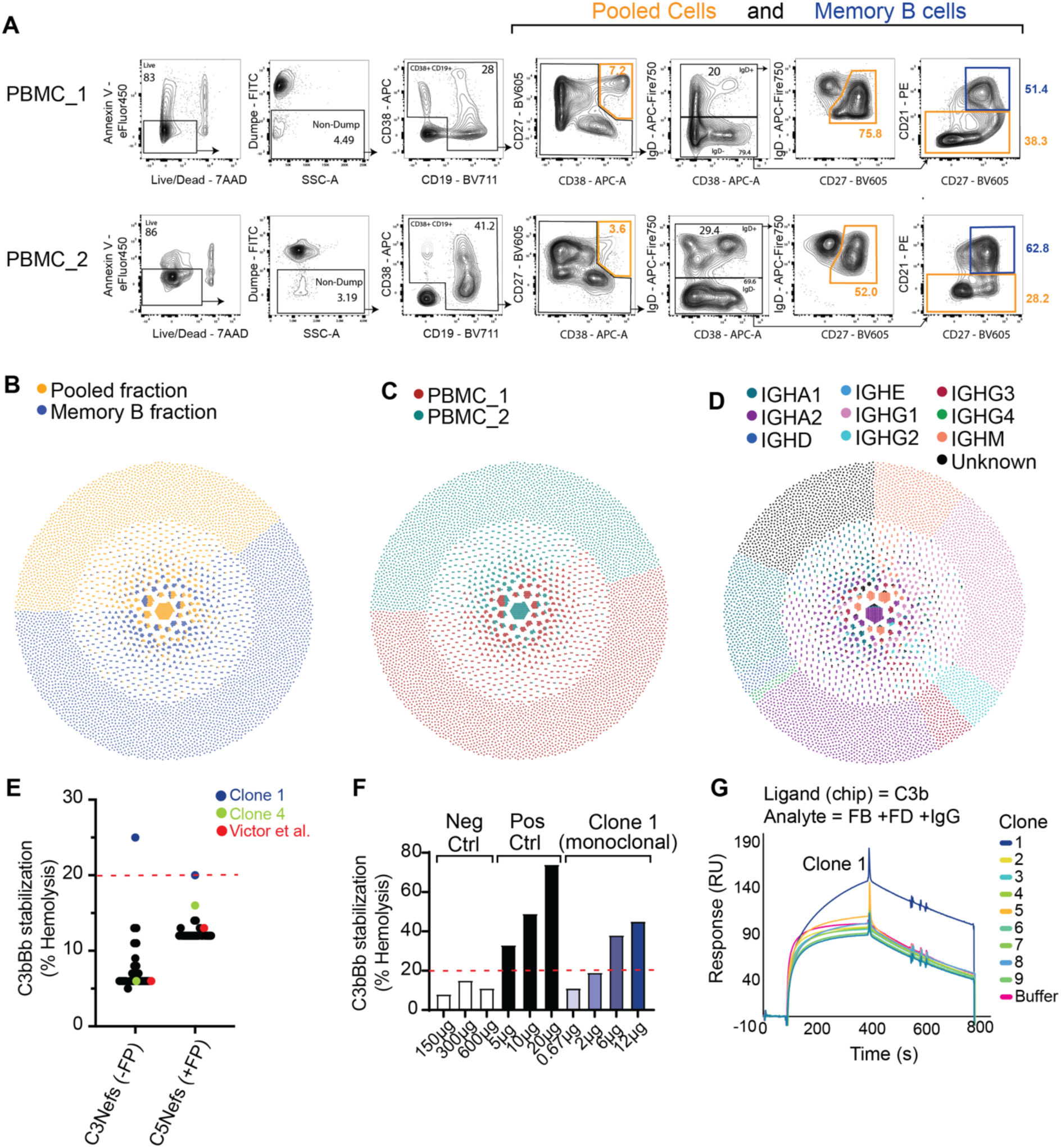
Obtaining and screening C3Nef candidates. **(A)** FACS plots showing PBMC gating strategy used to sort Memory B and Pooled (atypical memory, plasma cells, plasmablasts, and marginal zone-like B cells) samples from 2 different timepoints (PBMC_1 and PBMC_2) collected 9 months apart. Cells were sorted immediately prior to single-cell sequencing. Surface markers are listed on FACS plot axes. Numbers on FACS plots show percentage of parent population. (**B-D**) Honeycomb plots showing unique clonotypes for all PBMCs collected. Each dot is a single cell with ≥1 productive variable heavy and/or one paired variable light chain rearrangement. Cells are clustered into unique clonotypes using enclone software. Each honeycomb plot is the same dataset but is displayed using a different color scheme: (**B**) Cells obtained from Pooled fraction vs. Memory B fraction. (**C**) Cells obtained from timepoint 1 (PBMC_1) vs. timepoint 2 (PBMC_2). (**D**) Cells colored by B cell isotype. (**E**) Results from the initial hemolytic C3 convertase stabilization screen of patient-derived monoclonals in the absence (C3Nef assay) or presence (C5Nef assay) of properdin. All monoclonal IgGs are shown as black dots with the exception of Clone 1 (blue), Clone 4 (green), and the previously published Victor et al. clone (red) ^32^. 3µg of monoclonal IgG was used per test. Dashed red line indicates clinical cutoff for positive result (20%). (**F**) Same hemolytic assay as (**E**) but showing concentration-dependent increases in stabilizing properties for candidate Clone 1. Negative and positive polyclonal IgG samples shown in white and black, respectively. (**G**) Representative surface plasmon resonance sensorgram overlay showing properdin-independent reactivity of Clone 1 to a C3bBb convertase built on a SPR chip. Only sensorgram traces for the first 9 of the 28 tested clones are shown. RU, resonance unit.

**Fig. S3.**
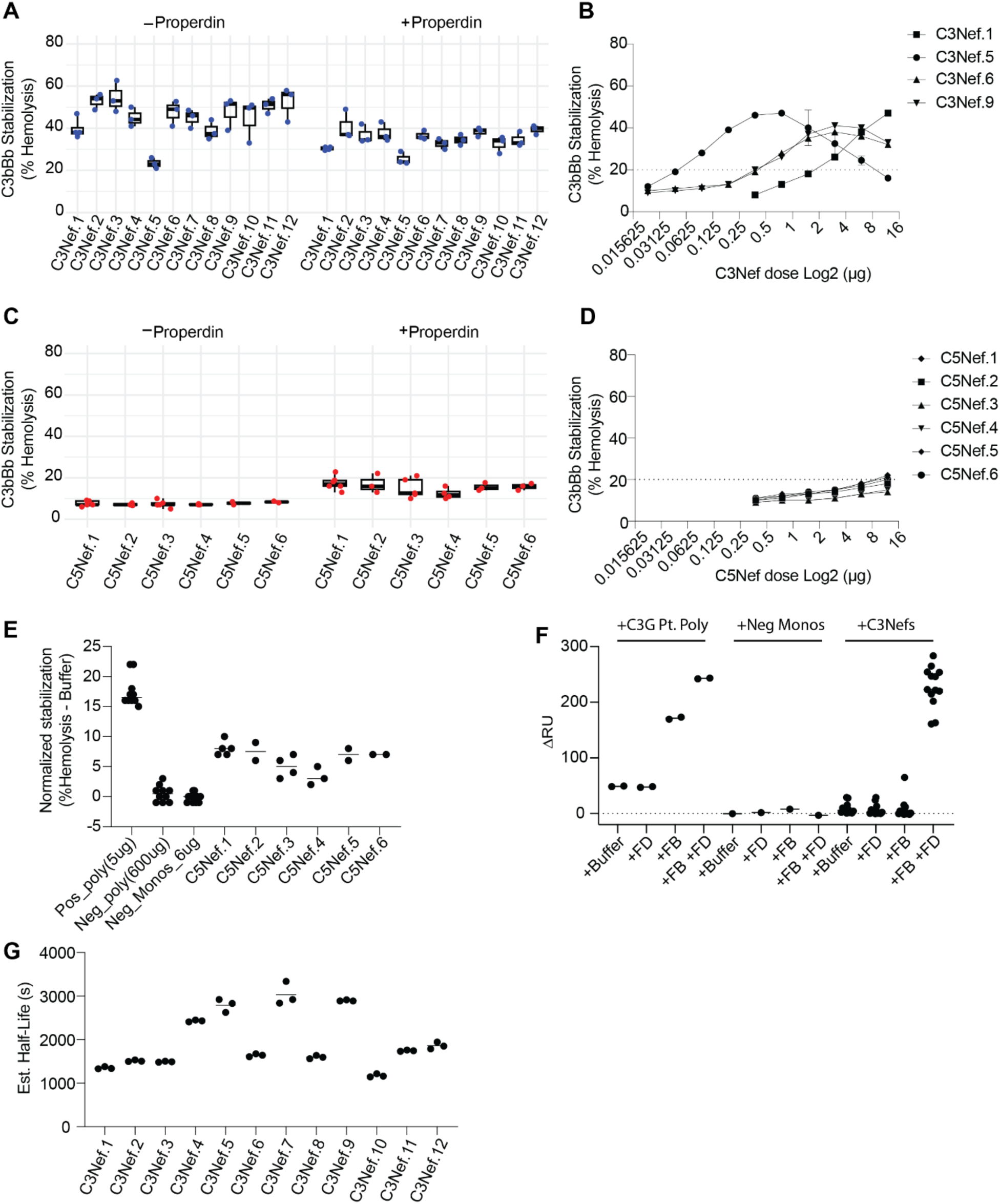
Convertase stabilizing properties of C3Nefs and C5Nefs. (**A**) Ability of individual properdin-independent C3Nefs (**A-B**) and properdin-dependent C5Nefs (**C-E**) to stabilize C3bBb(±P) convertase as measured by percent hemolysis in C3 convertase stabilization assays. (**A**) C3bBb stabilization by properdin-independent C3Nefs. n=3 technical replicates and 6µg for all clones. Clinical threshold for positive result, 20% hemolysis. (**B**) Same assay as (**A**) but showing concentration curve for select C3Nef subclones. (**C**) C3bBbP stabilization by properdin dependent C5Nefs. n=3 technical replicates and 12µg for all clones. (**D**) Same hemolytic assay as (**C**) but showing concentration effects for all 6 C5Nefs. (**E**) Same hemolytic convertase stabilization assay as (**A-D**), but with % Hemolysis normalized to buffer background and showing additional controls. Pos_poly and Neg_poly controls are clinically validated polyclonal normal human IgG-enriched samples. Neg_Monos are non-responsive monoclonals generated *ex vivo* from the same C3G index patient as the C5Nefs and expressed in the same IgG backbone as the C5Nefs. (**F**) Amine-coupled surface plasmon resonance (SPR) specificity assay showing reactivity of index patient’s polyclonal IgGs, negative monoclonals, and 12 monoclonal C3Nefs to either C3b (+Buffer / +FD), C3bB (+FB), or C3bBb (+FB +FD). Each dot shows the mean of n=2 technical replicates. (**G**) Estimated half-life values for all 12 C3Nefs as determined by SPR. Samples were run as 3 technical replicates. Ligand = C3b. Analyte = FB +FD +IgG. Calculations described in methods. RU, resonance units.

**Fig. S4.**
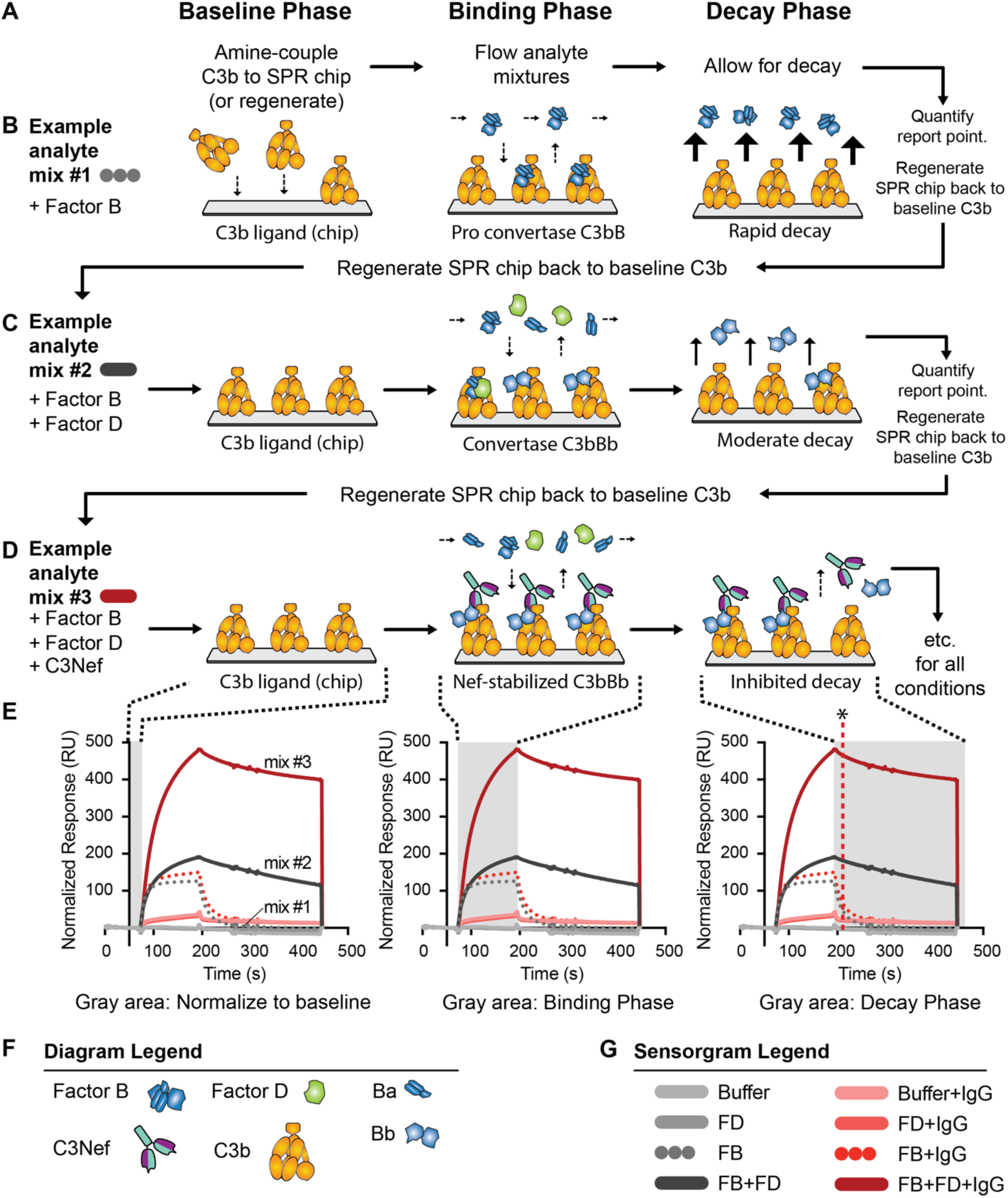
Workflow and diagram of Surface Plasmon Resonance (SPR) experiments to determine C3Nef reactivity to C3bBb convertase. (**A**) Basic workflow showing 3 iterative phases for each analyte cycle in a typical C3bBb reactivity assay by SPR. (**B-D**) Diagram showing 3 example cycles for a typical SPR reactivity experiment. (Not all conditions tested are shown in diagram. Details in methods). Individual analytes and/or analyte mixtures consisting of buffer, factor B, factor D, and/or C3Nef were prepared and floated over a C3b-amine-coupled SPR chip. Factor B can bind to C3b to form C3bB (pro-convertase). Factor D can then bind C3bB and cleave Factor B, forming the active convertase, C3bBb. Under normal *in vivo* conditions the C3bBb convertase is highly unstable and decays in ∼90 seconds. The addition of Nef into the analyte mixture binds to and stabilizes any C3bBb convertase present. (**E**) Representative SPR sensorgram overlays showing unique binding responses for different analyte mixtures. Shaded gray windows correspond to the cycle phase depicted diagrammatically above each sensorgram overlay (**B-D**). Vertical dashed red line with black asterisk (far right) marks the RU report point. (**F**) Diagram legend. (**G**) SPR sensorgram legend. FB, factor B; FD, factor D, RU, resonance units.

**Fig. S5.**
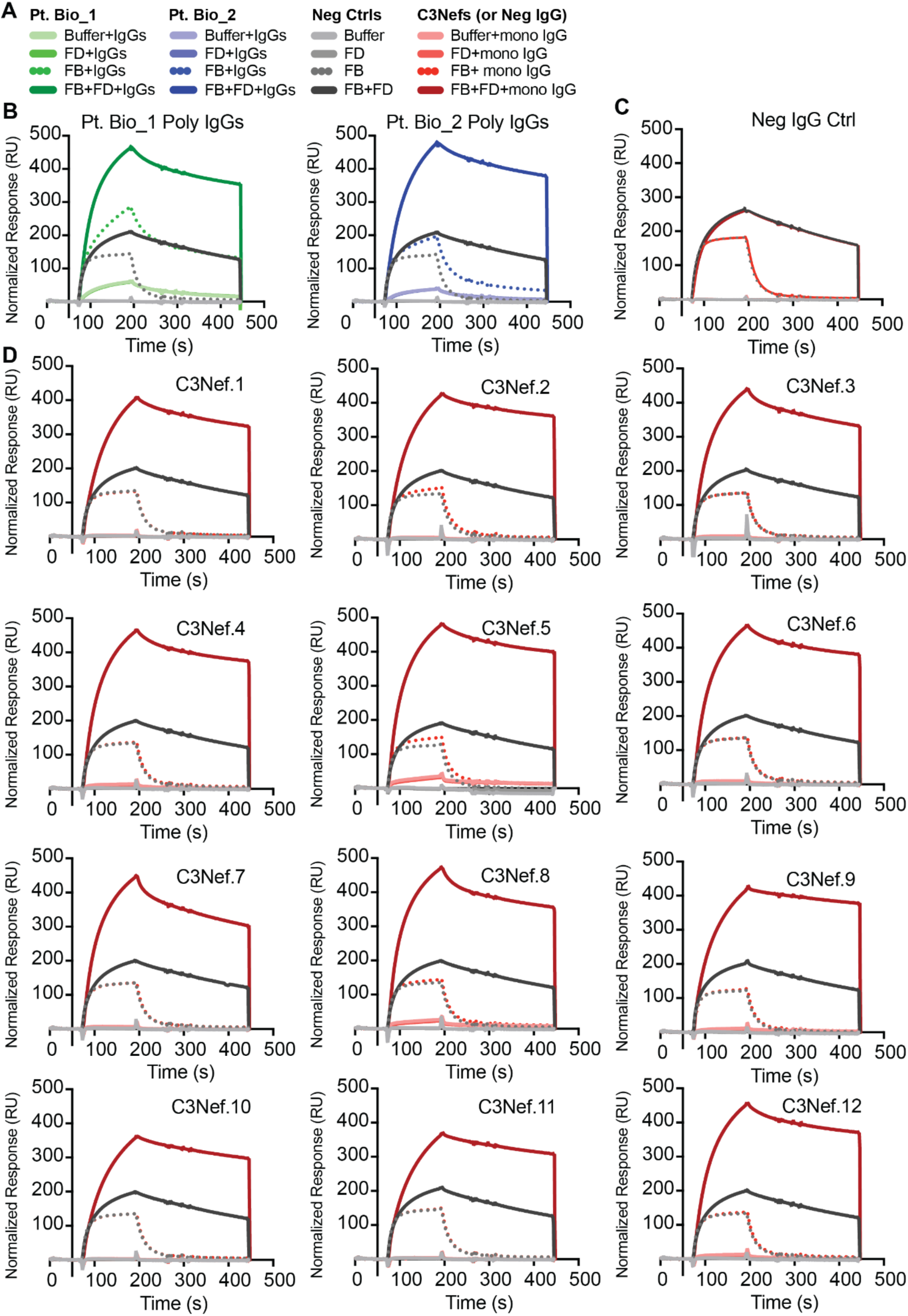
C3Nefs bind the C3bBb convertase. (**A**) Sensorgram legend showing all samples and analyte mixtures testing reactivity with C3b amine-coupled to SPR chip. See fig. S4 and Methods for details. Neg control analyte mixtures were repeated with each individual mono- and polyclonal run and are shown in grayscale overlay for each run. (**B**) Individual sensorgram overlays for polyclonal patient IgG samples (Bio_1 and Bio_2). Polyclonal IgGs were obtained at the same timepoints as the Nef-expressing PBMCs. Note the reactivity of the polyclonal IgGs to the pro-convertase (C3bB; FB+IgGs) and to C3b coupled to the SPR chip (C3b; Buffer+IgGs). (**C**) Sensorgram overlay for Neg Ctrl IgG monoclonal from C3G index patient showing no reactivity to any analyte mixtures. (**D**) Sensorgram traces for all 12 individual C3Nefs. Note that robust C3Nef binding only occurs following the formation C3bBb convertase (C3bBb; FB+FD+ mono IgG), with little-to-no binding of the C3Nef to C3bB proconvertase or C3b. RU, resonance units.

**Fig. S6.**
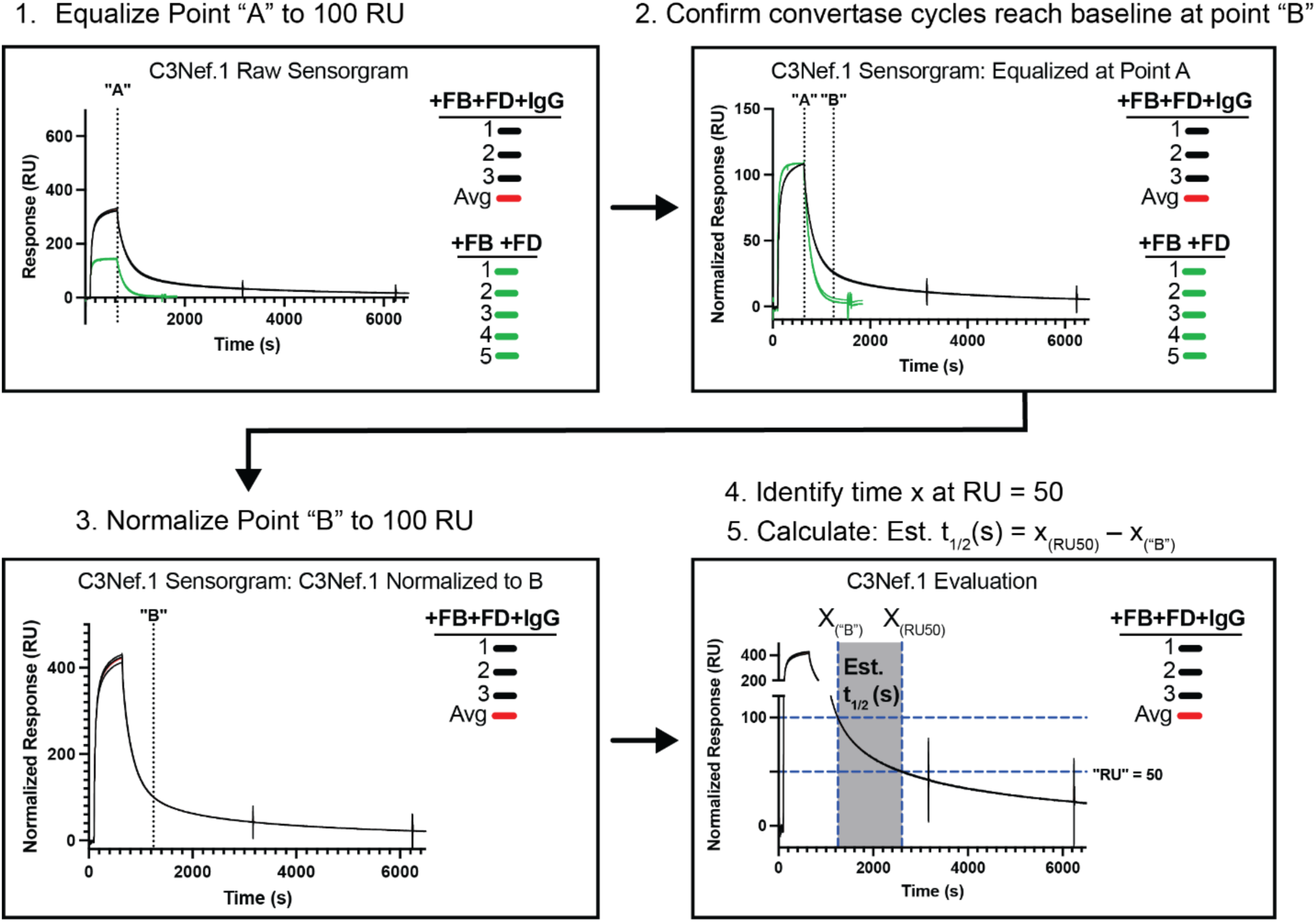
Methodology for determining C3Nef estimated half-life values. Flow chart outlining the workflow and calculation of estimated half-life binding times determined by SPR at 37°C. See methods for details. RU, resonance units.

**Fig. S7.**
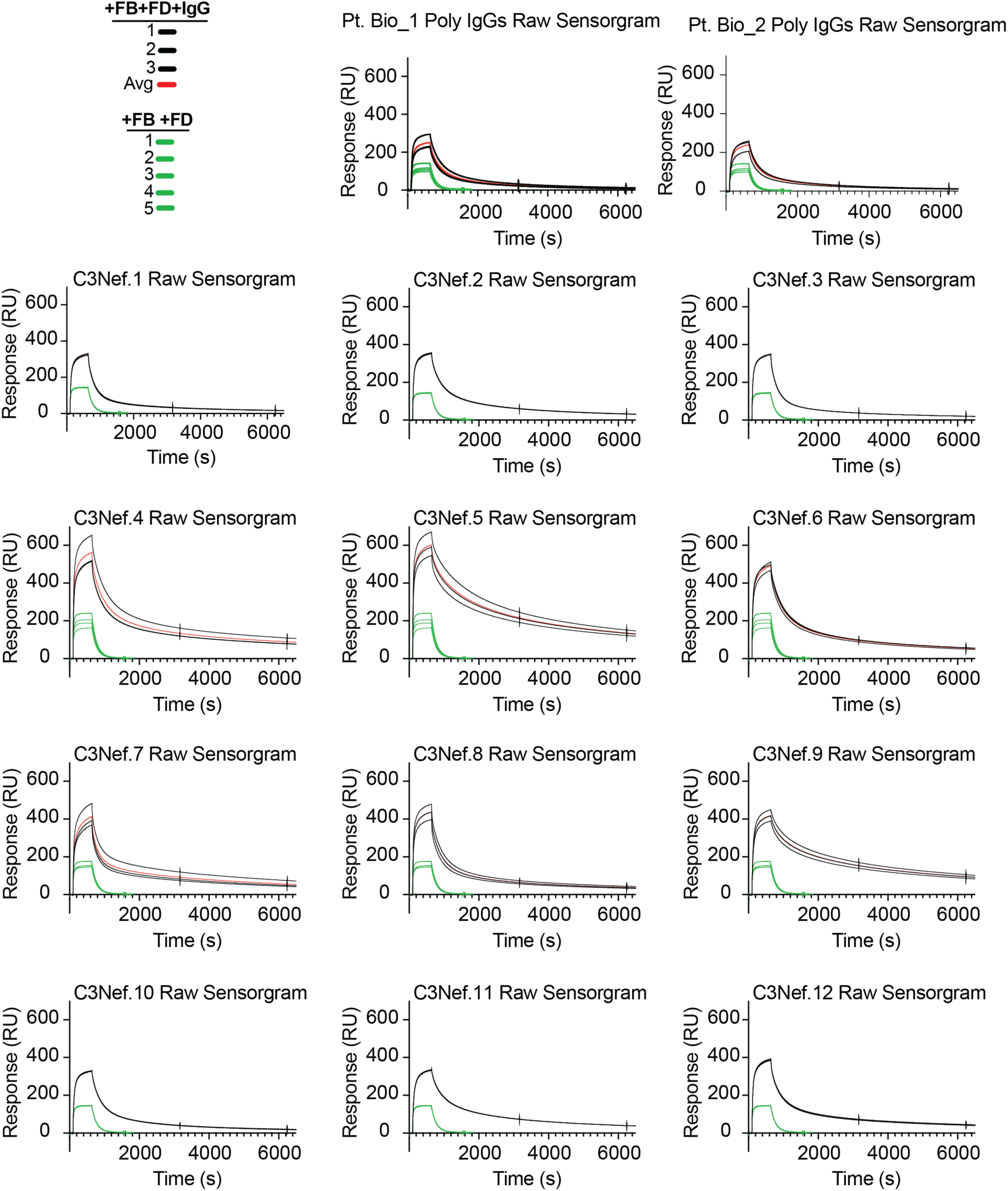
Estimated half-life calculation: Step 1. SPR overlays of raw sensorgram data for patient polyclonal IgGs and all 12 C3Nefs used for “Point A equalization” of estimated half-life calculations. RU, resonance units; FB, factor B; FD, factor D.

**Fig. S8.**
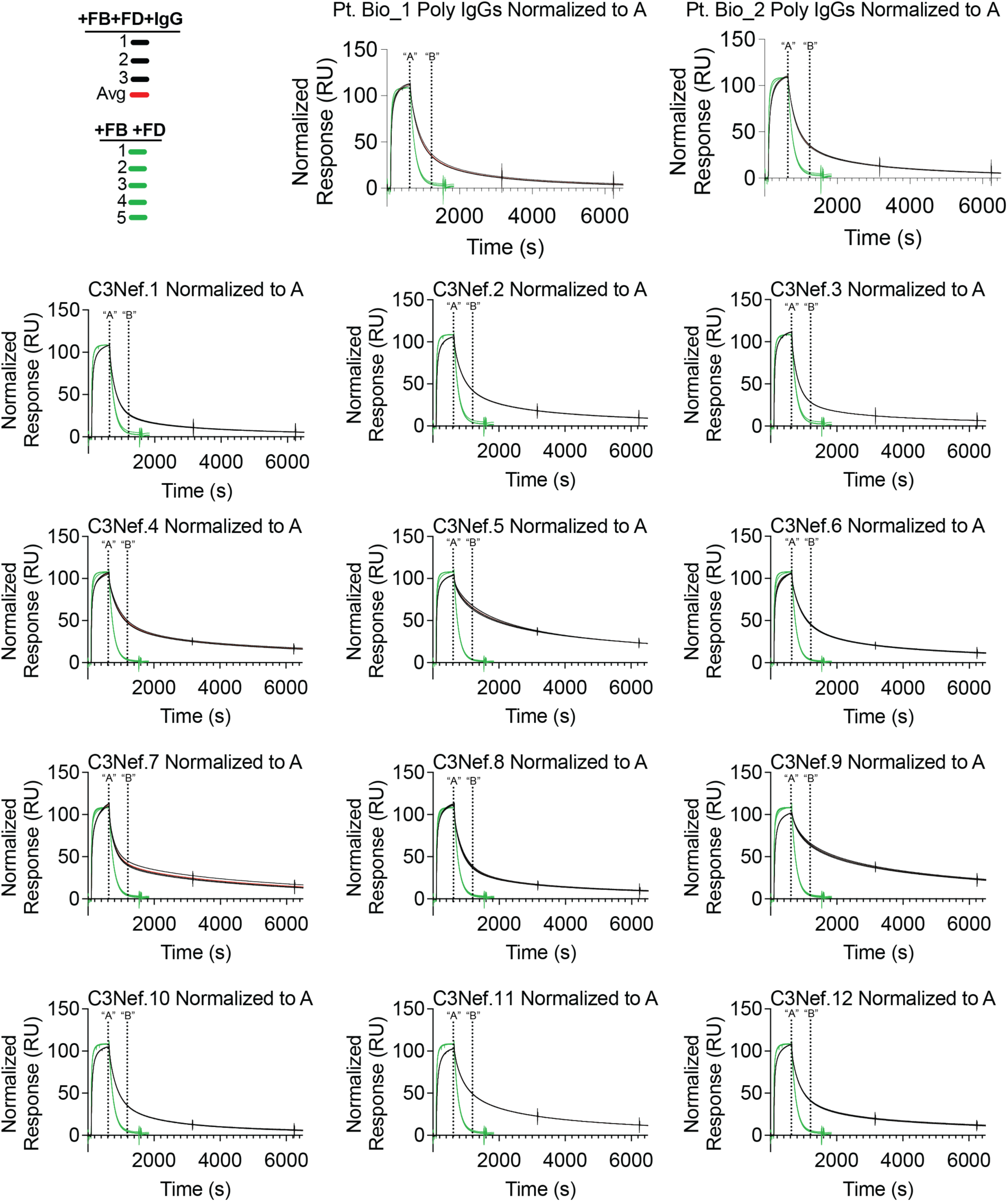
Estimated half-life binding: Steps 2. Raw C3Nef sensorgram data from fig. S7 normalized to point A as shown outlined fig. S6. Importantly, all convertase-only sensorgram traces reach baseline RU = 0 by point B. RU, resonance units; FB, factor B; FD, factor D.

**Fig. S9.**
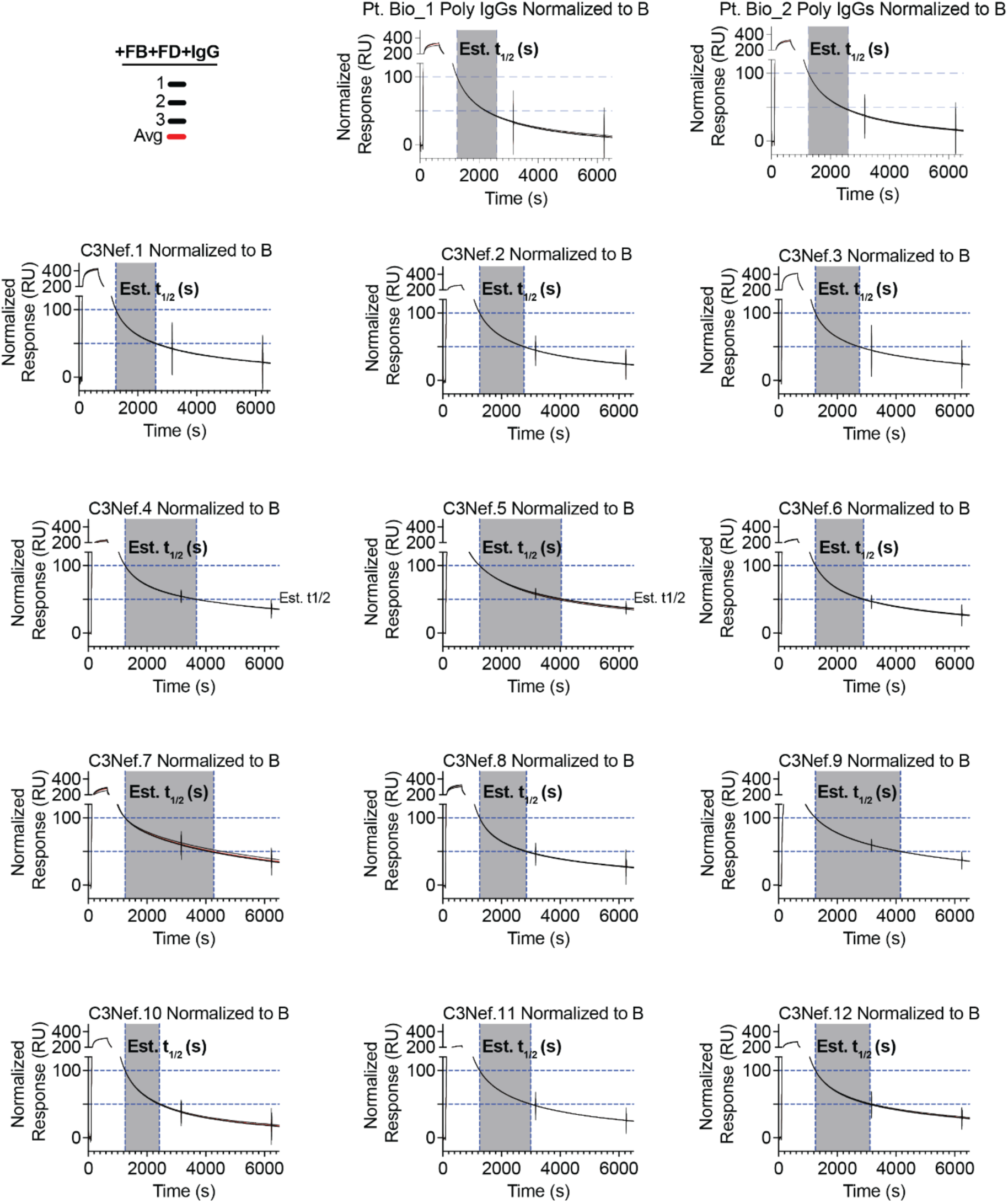
Estimated half-life binding: Steps 3-5. C3Nef sensorgrams normalized to point B. Gray shading indicates the estimated half-life window as shown outlined fig. S6. RU, resonance units; EST, estimated half-life; FB, factor B: FD, factor D.

**Fig. S10.**
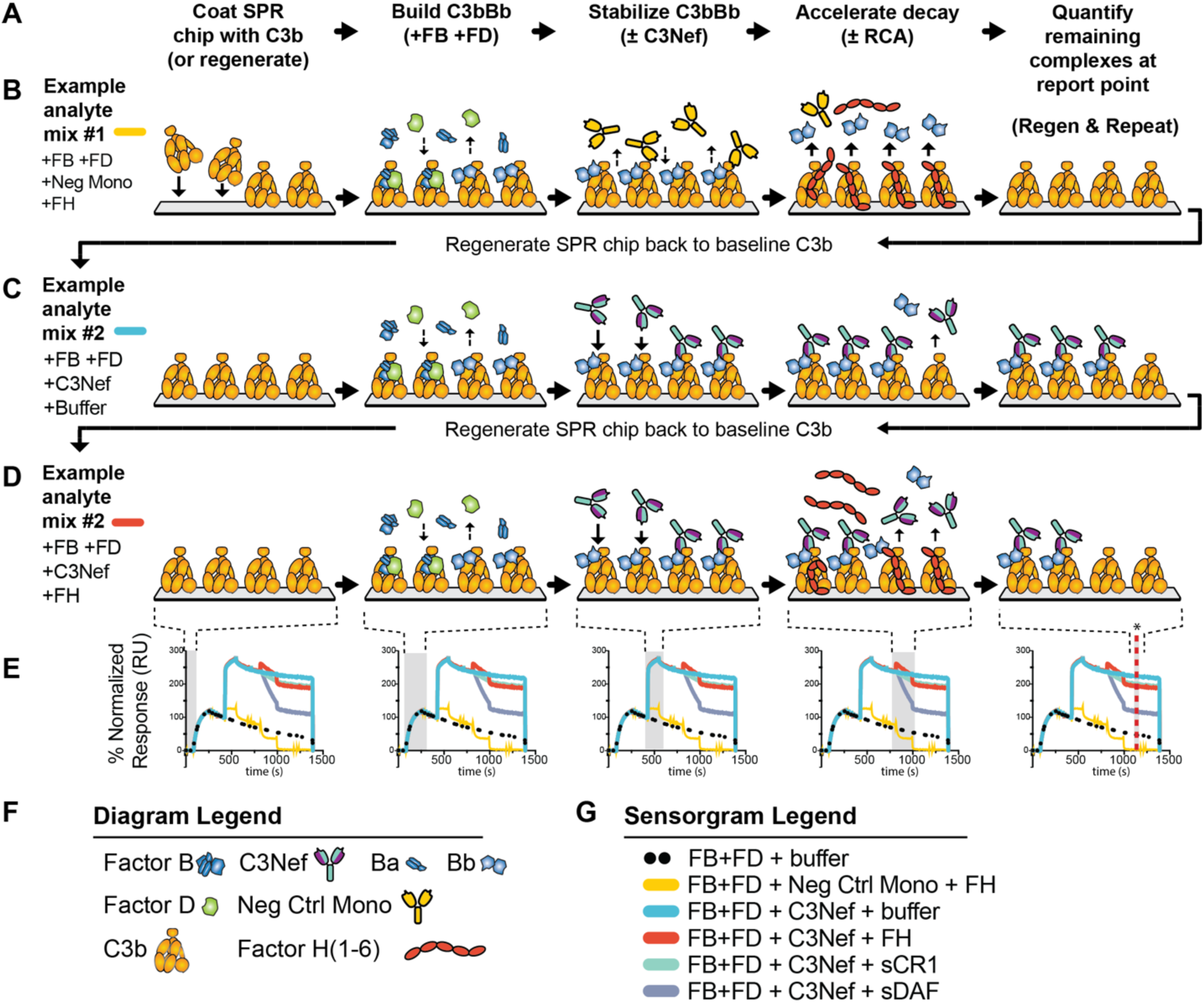
Workflow and diagram of surface plasmon resonance (SPR) experiments used to identify C3Nef-mediated inhibition of RCA-accelerated convertase decay. (**A**). Basic SPR workflow showing 5 iterative phases for each analyte cycle used to test effects of C3Nef on RCA-accelerated decay assay. (**B-D**) Diagram depicting 3 representative analyte mixture cycles tested for RCA-mediated decay (not all conditions tested are shown in diagram). (**E**) Representative SPR sensorgram overlays for all RCA conditions tested. Shaded gray areas correspond to the phases depicted diagrammatically above the sensorgram in (**B-D**). Vertical dashed red line with black asterisk (far right panel) denotes RU report point. (**F**) Diagram legend. (**G**) SPR sensorgram legend for all RCA cycles (analyte mixtures) tested. Mono; monoclonal antibody. FH; factor H short consensus repeat domains 1-6. sCR1; soluble complement receptor protein 1 domains 1-3. sDAF; soluble decay accelerating factor domains 1-4.

**Fig. S11.**
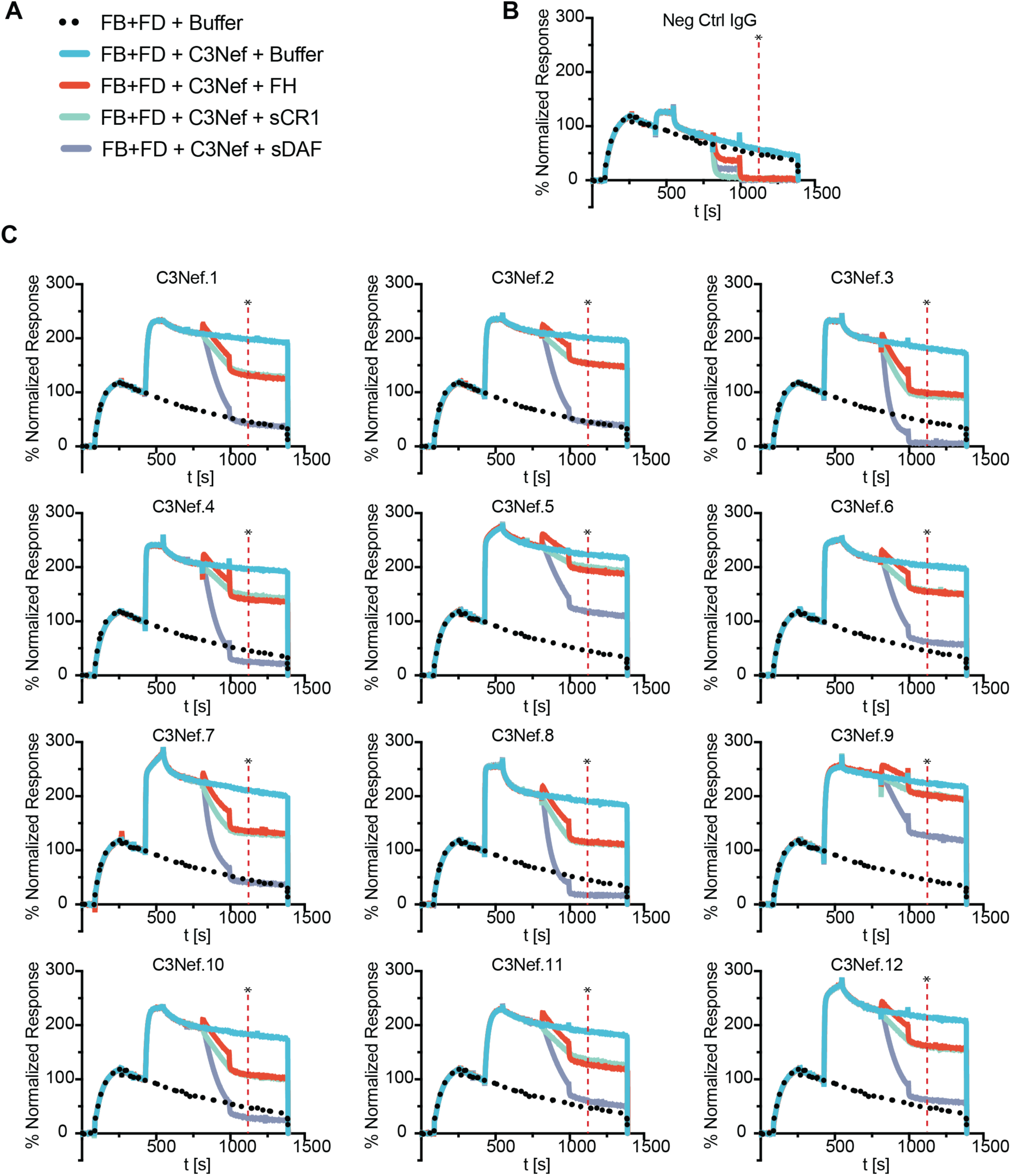
C3Nefs inhibit the decay acceleration activity of regulators of complement activity. (**A**) Sensorgram legend showing analyte conditions tested for all twelve C3Nefs. (**B**) SPR sensorgram overlay showing neg monoclonal, which was an IgG from the C3G patient that displayed weak convertase-reactive and was used as a negative control. Note the return of the sensorgram to baseline C3b levels following RCA treatment. (**C**) SPR sensorgram overlays for each individual C3Nef. Each trace shows a 4-cycle overlay of Nef-stabilized C3bBb that is exposed to 3 different regulators of complement activity: FH, factor H domains 1-6; sCR1, soluble complement receptor 1 domain 1-3; sDAF, soluble decay accelerating factor CCP domains 1-4; or buffer-only control. Amount of remaining convertase is quantitated and shown in Fig. 1F.

**Fig. S12.**
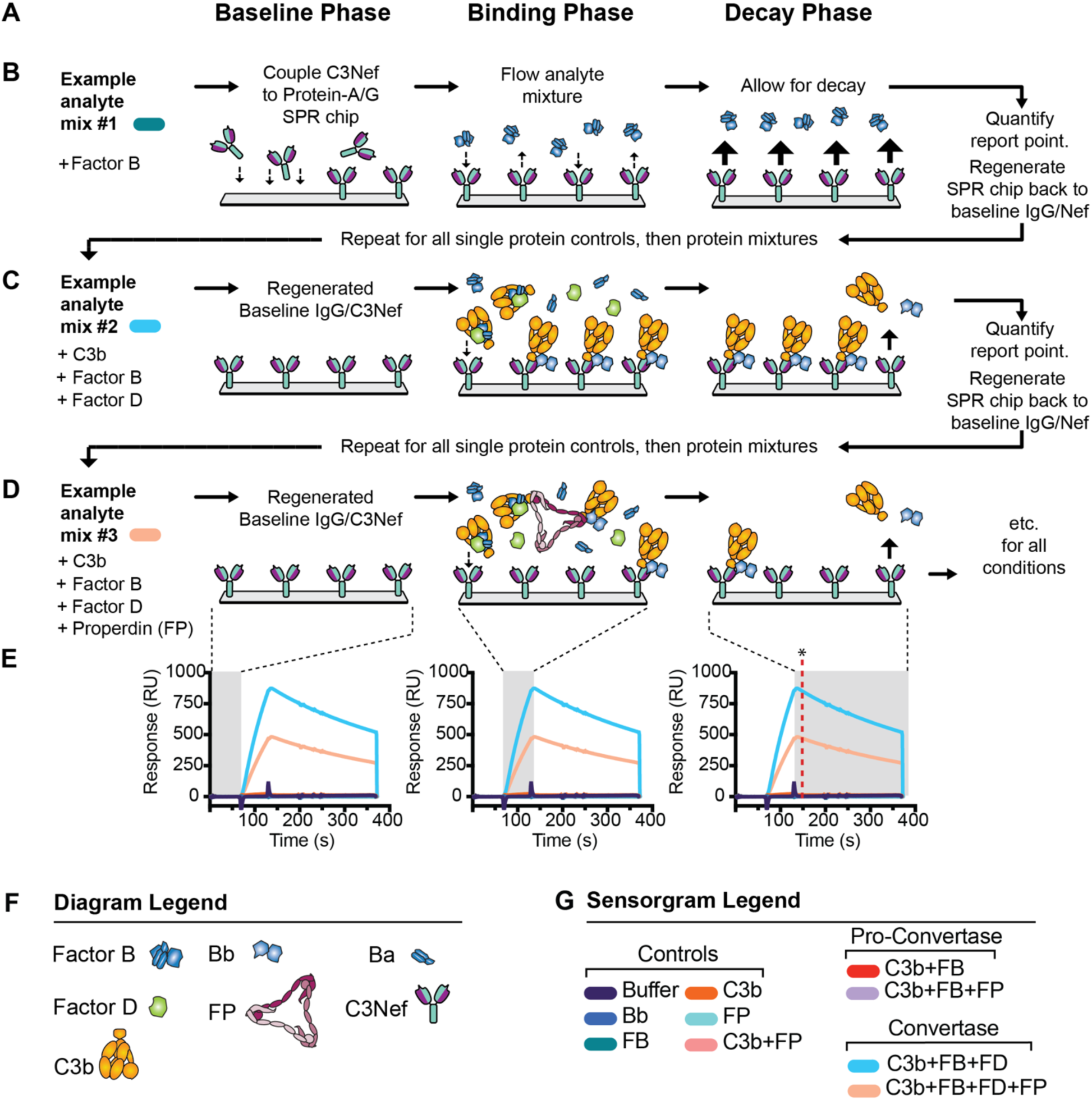
Workflow and diagram of Protein G-coupled surface plasmon resonance (SPR) experiments used to characterize C3Nef binding responses to C3 convertase and convertase components. (**A**) Workflow listing iterative phases of protein-G C3Nef-coupled SPR assay. (**B-D**) Diagram showing three representative analyte cycles for typical protein-G C3Nef-coupled SPR assay (not all conditions tested are shown in diagram). See Methods for details. (**E**) Representative SPR sensorgram overlays showing all cycles tested for C3Nefs. Shaded gray windows correspond to the phase depicted diagrammatically above each sensorgram. Vertical dashed red line with black asterisk (far right) denotes RU report point. (**F**) Diagram legend. (**G**) Sensorgram legend listing all conditions tested for each C3Nef-coated SPR chip. FP, factor P / properdin; FB, factor B; FD, factor D.

**Fig. S13.**
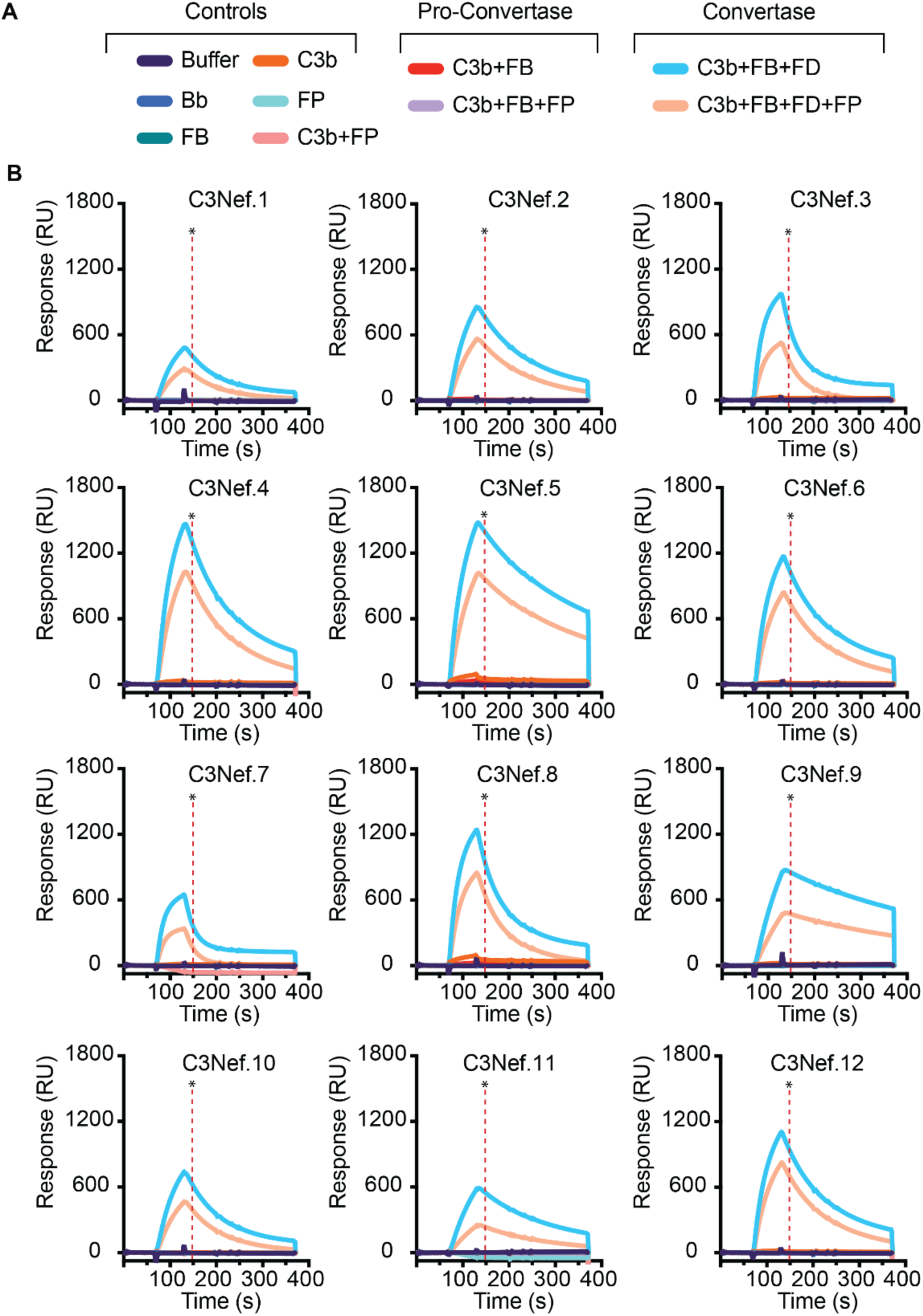
C3Nef binding characterization by SPR. (**A**) Sensorgram legend showing all analyte conditions tested for C3Nef binding as outlined in fig. S12. SPR chip was coated with C3Nef via protein G-coupling. (**B**) SPR sensorgram overlays for all twelve C3Nefs and all analyte conditions. Sensorgram traces show the mean of two separate SPR experiments for each Nef and condition tested. All cycles listed in (**A**) are ran in duplicate for each individual C3Nef. Vertical dashed red line with asterisk denotes RU report point. FB, factor B; FD, factor D; FP, properdin/factor P; RU, resonance units.

**Fig. S14.**
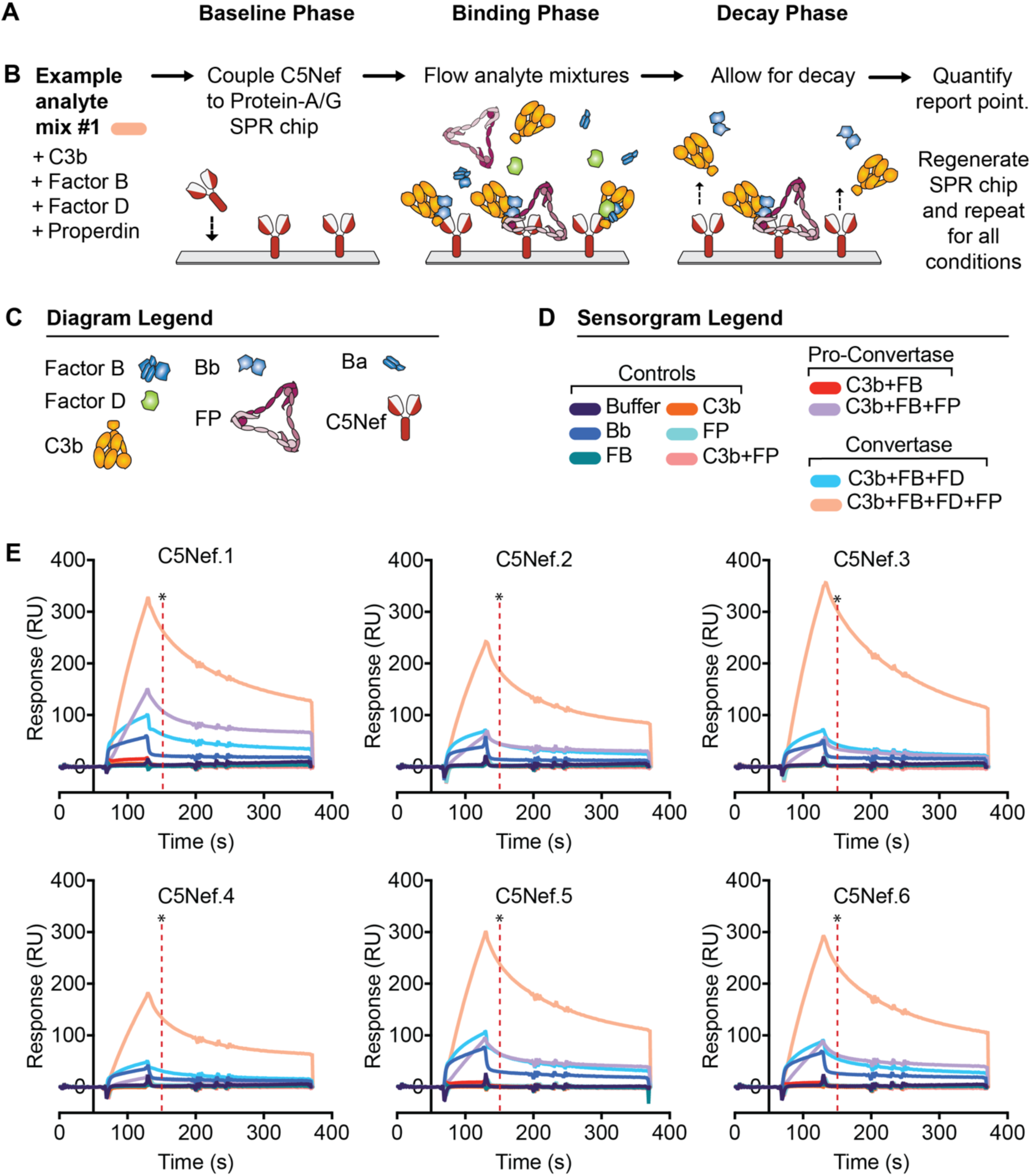
C5Nef binding characterization by SPR. (**A**) Workflow outlining iterative phases of protein-G C5Nef-coupled SPR assay. (**B**) Diagram showing one representative analyte cycle for protein-G C5Nef-coupled SPR assay (not all conditions tested are shown in diagram). See fig. S12 and Methods for details. (**C**) Diagram legend. (**D**) Sensorgram legend showing all conditions tested on C5Nef-coated SPR chip. (**E**) SPR sensorgram overlays for all 6 C5Nefs tested. Each trace represents the mean of two technical replicates. All conditions listed in (**D**) were run in duplicate for each C5Nef. Vertical dashed red line with black asterisk denotes RU report point. FP, factor P / properdin; FB, factor B; FD, factor D; RU, resonance units.

**Fig S15.**
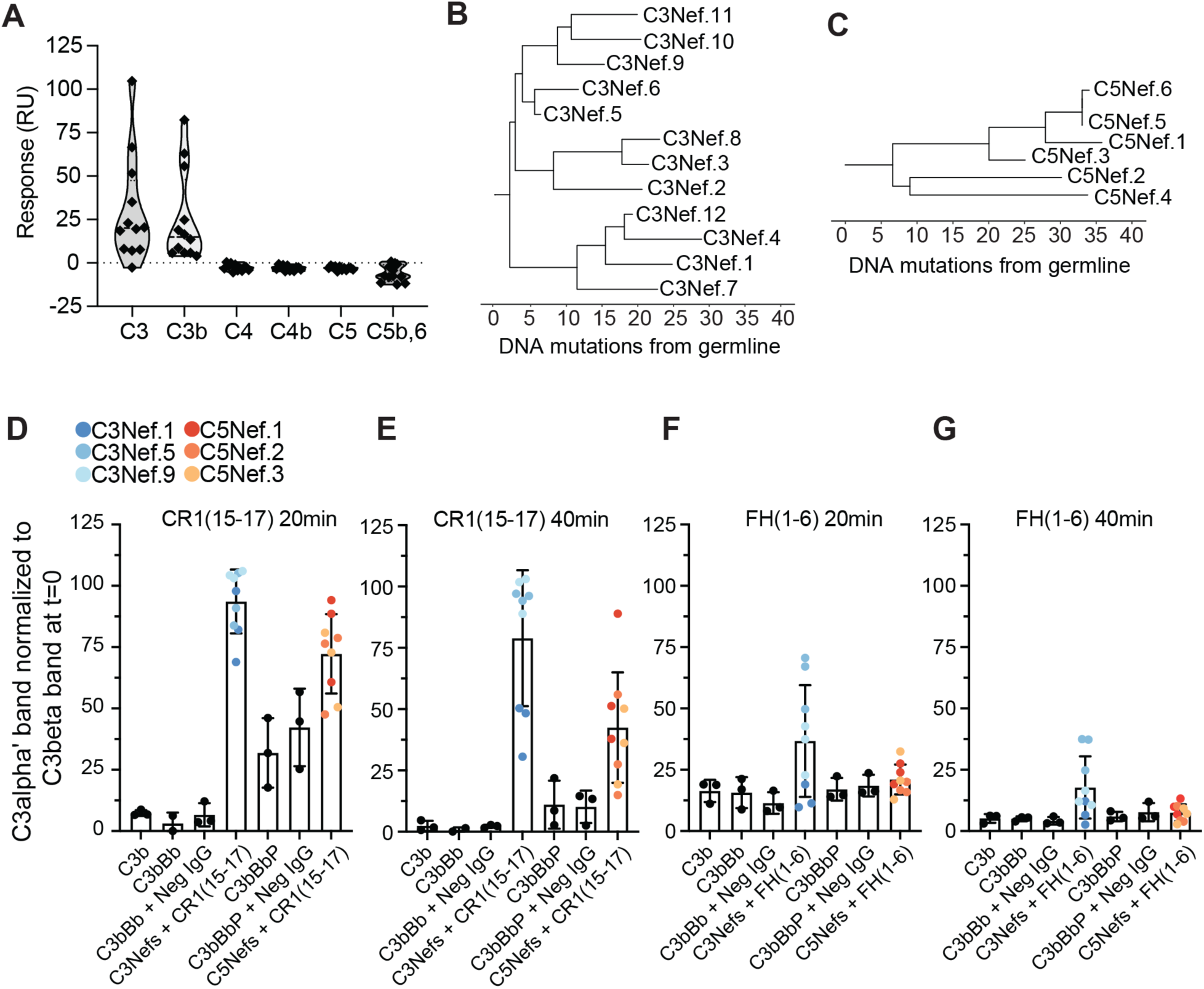
C3Nefs are distinct from C5Nefs. (**A)** C3Nefs do not bind C4 or C5 proteins. SPR response of protein G-coupled C3Nefs to various C345c-containing protein analytes. **(B-C**) Phylogenic trees generated using enclone software showing the Levenshtein distances between C3Nef (**B**) and C5Nef (**C**) subclones. Predicted number of mutations from germline shown on x axis. (**D-G**) Bar graphs showing the ability of select C3Nefs (blue) or C5Nefs (red) to block the cofactor activity of sCR1(15-17)+FI (**D-E**) or FH(1-6)+FI (**F-G**) in a fluid-phase cofactor activity assay. n=3 for each Nef tested. Anti-RhD used as Neg IgG. sCR1, soluble complement receptor protein 1 domains 15-17; FH, factor H short consensus repeat domains 1-6; FI, factor I.

**Fig. S16.**
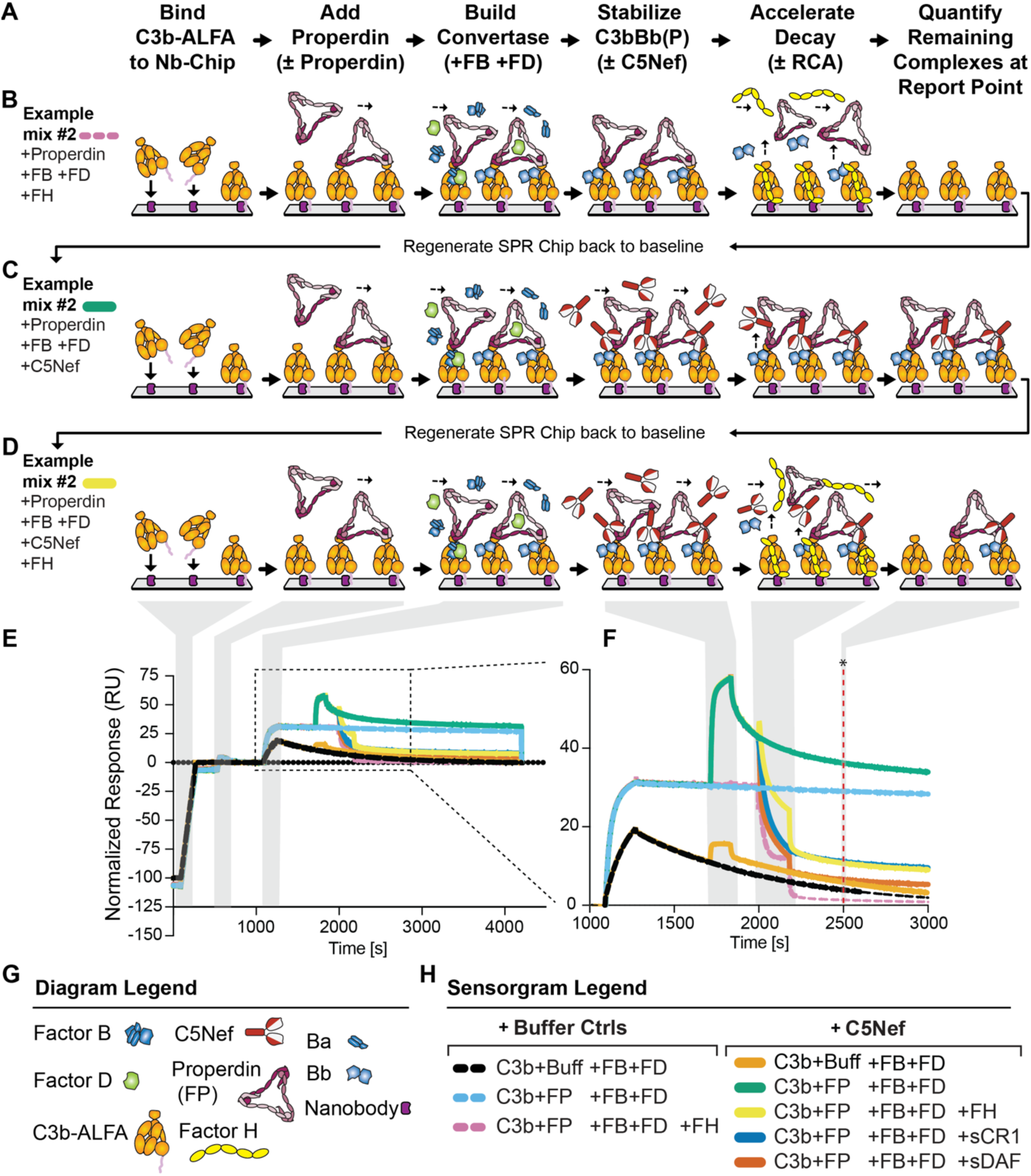
Workflow and diagram of surface plasmon resonance (SPR) experiments used to characterize C5Nef inhibition of RCA accelerated convertase decay. **(A)** Workflow outlining iterative phases to examine effects of C5Nefs on regulators of complement activity (RCAs) convertase decay acceleration. **(B – D)** Diagrams depicting three representative cycle conditions (not all cycle conditions tested are shown). See Methods for details. **(E-F)** Representative SPR sensorgram overlays showing all cycle conditions tested. Full sensorgram trace shown in (**E**), with zoomed-in insert shown in (**F**). Shaded gray areas correspond to the phase depicted diagrammatically above each sensorgram. Vertical dashed red line with black asterisk (far right) denotes RU report point. **(G)** Diagram legend. **(H)** Sensorgram legend showing all conditions tested. FP, factor P/properdin; FB, factor B; FD, factor D; Buff, buffer control; RCA, regulators of complement activity; FH, factor H domains 1-6; sCR1, soluble complement receptor 1 domains 1-3; sDAF, soluble decay accelerating factor domains 1-4. SPR; surface plasmon resonance.

**Fig. S17.**
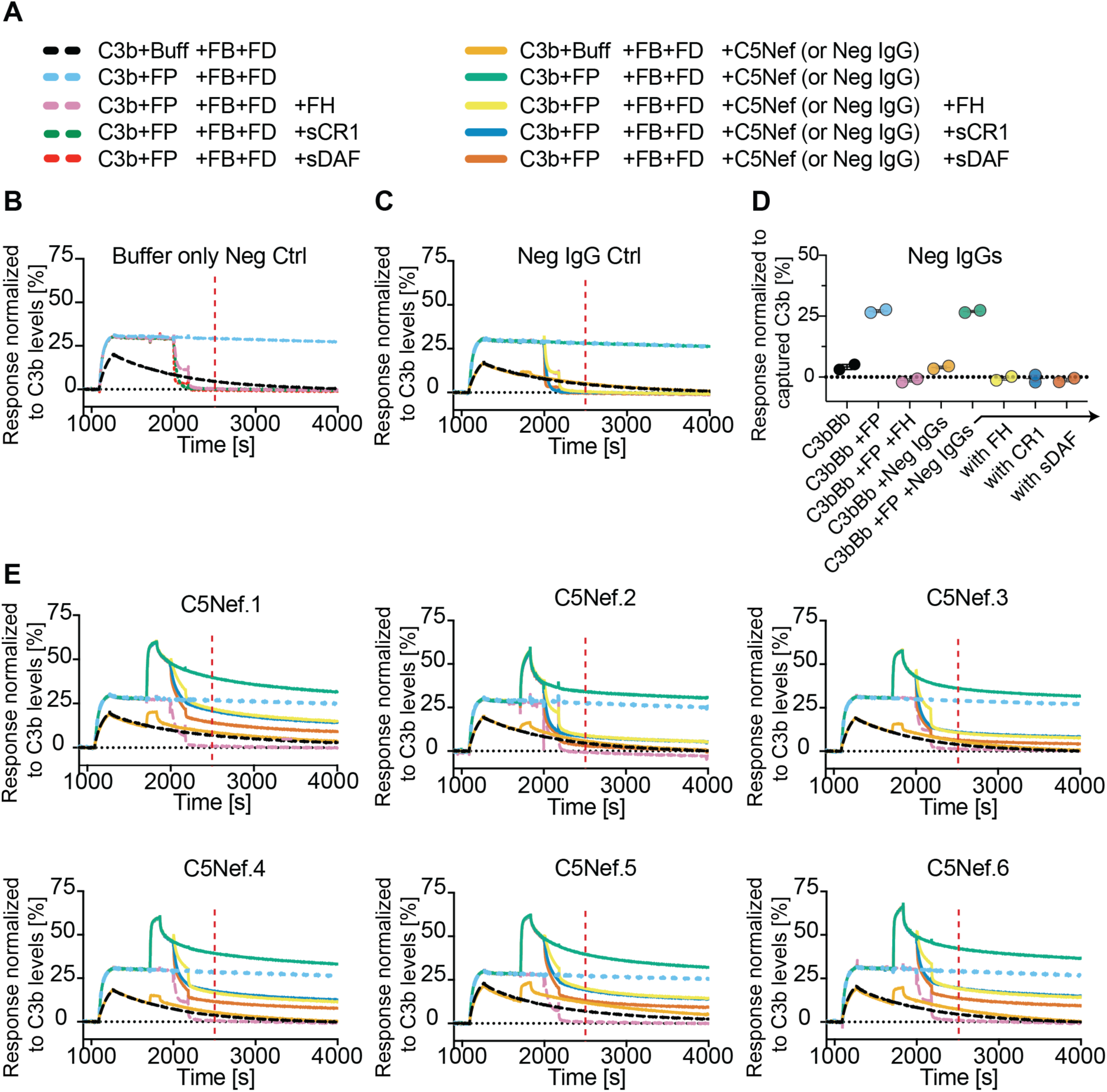
C5Nefs inhibit the decay acceleration activity of regulators of complement activity. (**A**) Sensorgram legend showing all cycles run in duplicate for buffer only, monoclonal Neg IgG, or all C5Nefs to determine effects of C5Nefs on regulators of complement activity decay acceleration. (**B**) Buffer-only negative control showing the effects of RCAs on Properdin-stabilized C3bBb convertase only. Vertical dashed red line indicates RU report point for values shown in (**D**). (**C**) Neg IgG control (non-Nef monoclonal IgG from C3G patient) showing no inhibition of RCA-accelerated decay of C3bBb. (**D**) Jitter plot analysis of SPR response for Neg monoclonal IgGs for all cycles tested at RU report point 2500[s] (normalized). n = 2 technical replicates. (**E**) SPR sensorgram overlays for all 6 C5Nefs. Each trace on the sensorgram is the mean of two technical replicates. Analyte mixtures tested are listed in legend (**A**). FB, factor B; FD, factor D; Buff, buffer control; FH; factor H short consensus repeat domains 1-6. sCR1; soluble complement receptor protein 1 domains 1-3; sDAF; soluble decay accelerating factor domains 1-4.

**Fig. S18.**
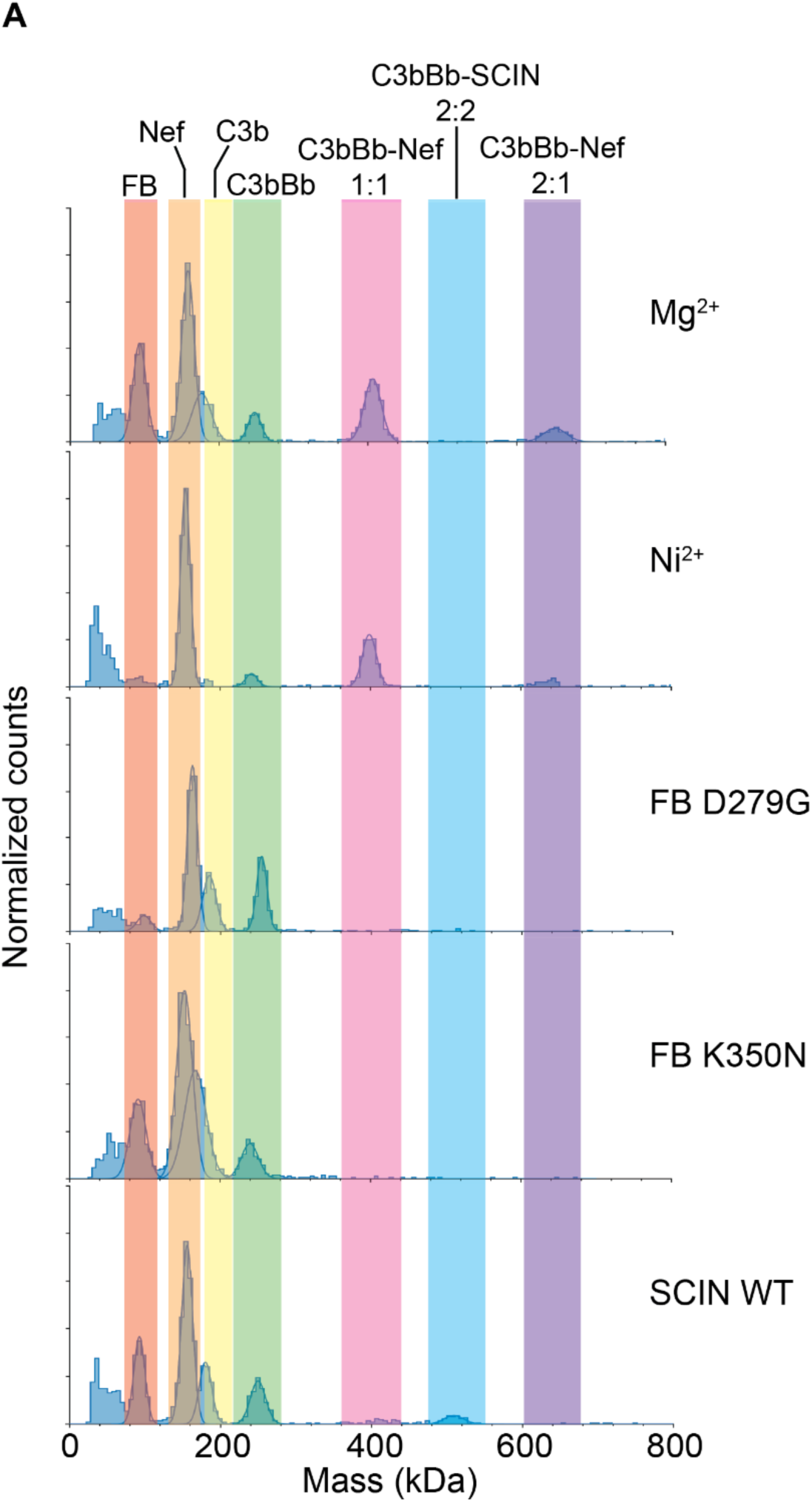
C3Nef binding to C3bBb is hindered by convertase-stabilizing modifications. (**A**) Representative histogram overlays showing mass photometry normalized counts (y axis) across measured molecular masses (x axis). Normalized peaks show the ability of C3Nef to form various convertase complexes under normal physiologic conditions (Mg^2+^ top panel) or reported convertase-stabilizing conditions (all other conditions, lower panels). From top to bottom: Normal WT C3bBb generated using a mixture of C3b +factor B, and +factor D in a buffer supplemented with Magnesium (Mg^2+^), as compared with C3bBb stabilized by Nickel (Ni^2+^), C3bBb stabilized by factor B mutations D279G or K350N, or C3bBb stabilized by WT Staphylococcal Complement Inhibitor (SCIN-WT).

**Fig. S19.**
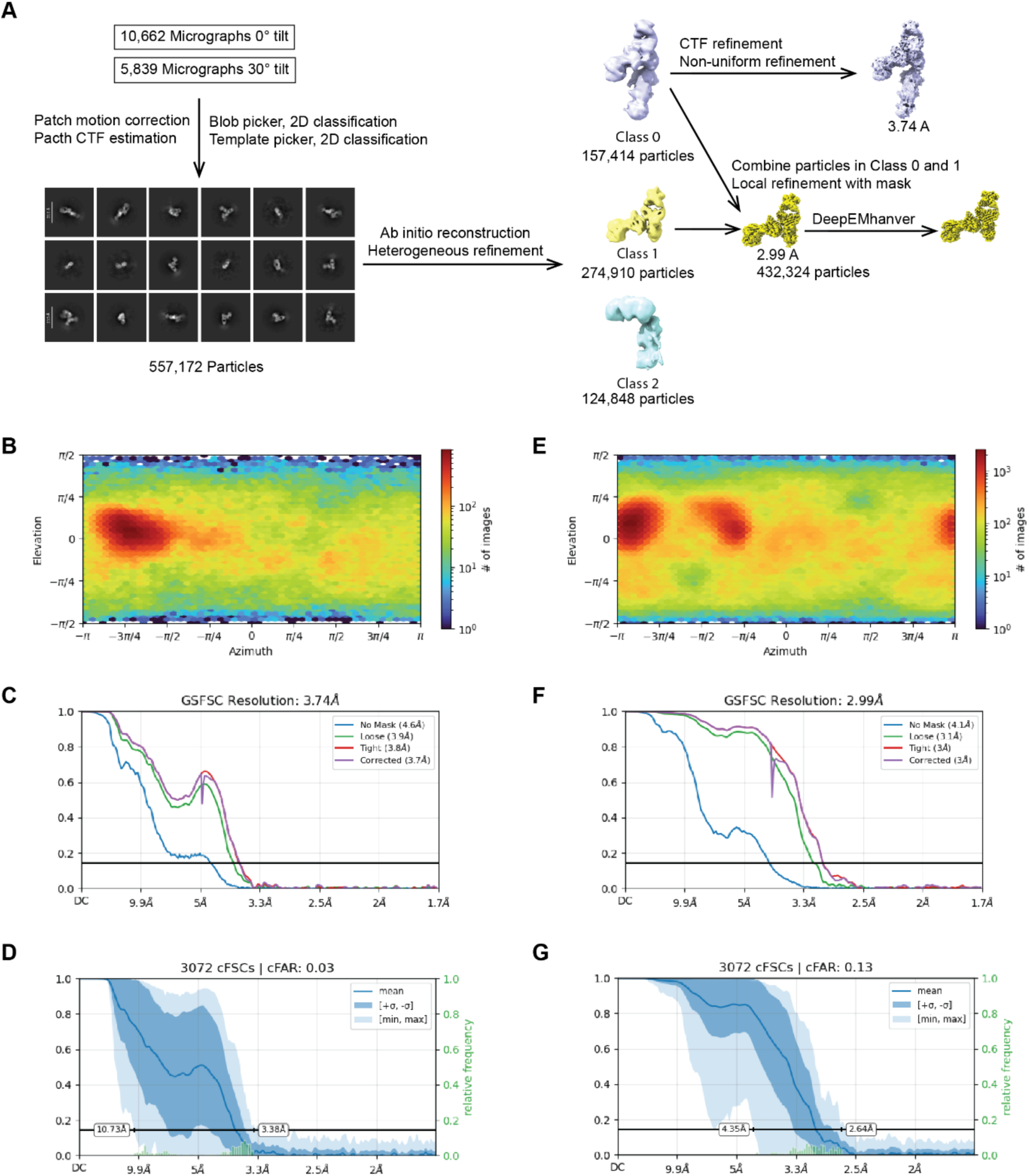
Cryo-EM reconstruction of the C3Nef.9-stabilized C3bBb complex. (**A**) Workflow of cryo-EM data processing for the C3Nef-stabilized C3bBb complex. Two datasets were collected in 0° and 30° tilt angle, respectively, and processed independently before ab initio reconstruction. Two maps were constructed from the dataset (3.74 Å and 2.99 Å overall resolution). 3.74 Å map has an extra density of MG ring. (**B-D**) Angular particle distribution map, gold standard Fourier shell correlation (GSFSC) and conical FSC (cFSC) curves of 3.74 Å map. (**E-G**) Angular particle distribution map, gold standard Fourier shell correlation (GSFSC) and conical FSC (cFSC) curves of 2.99 Å map.

**Fig. S20.**
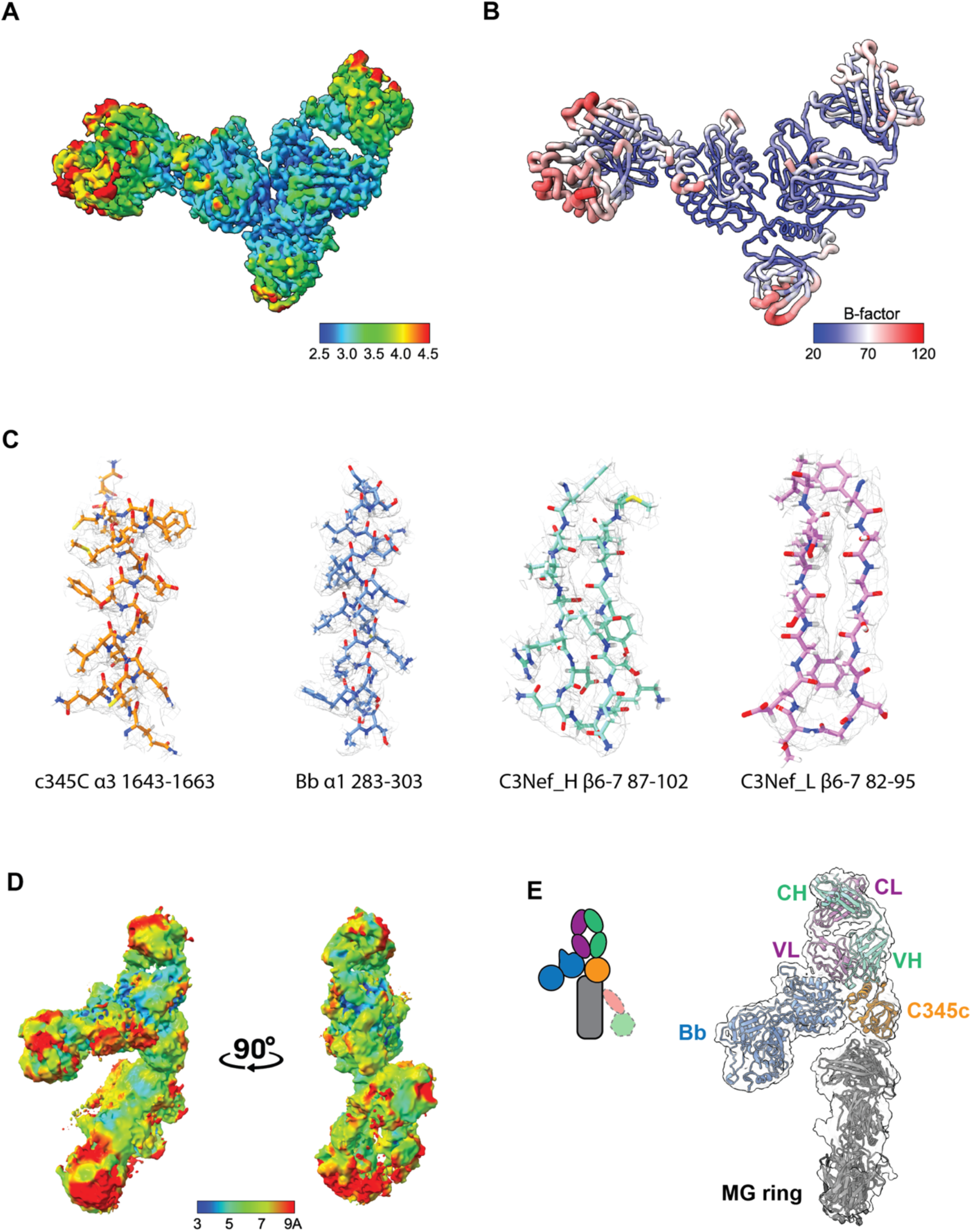
Structural analysis the C3Nef.9-stabilized C3bBb complex. (**A**) Local resolution map of the C3Nef-stabilized C3bBb complex in 2.99 Å overall resolution. (**B**) Putty tube representation of B-factor for the C3Nef-stabilized C3bBb complex in 2.99 Å overall resolution. (**C**) Representative density covering of C3 C345c (orange), Bb (blue), C3Nef heavy chain (aquamarine) and light chain (purple). (**D**) Local resolution map of the C3Nef-stabilized C3bBb complex in 3.74 Å overall resolution. (**E**) Diagram (left) and map fitted with model for lower resolution Nef-bound C3bBb showing partial MG ring resolution. CH, constant heavy chain; CL, constant light chain; VH, variable heavy chain; VL, variable light chain; MG, macroglobulin domain. Heavy chain shown in aquamarine, light chain in purple, C345c domain of C3b in orange, Bb in blue, MG ring in grey.

**Fig. S21.**
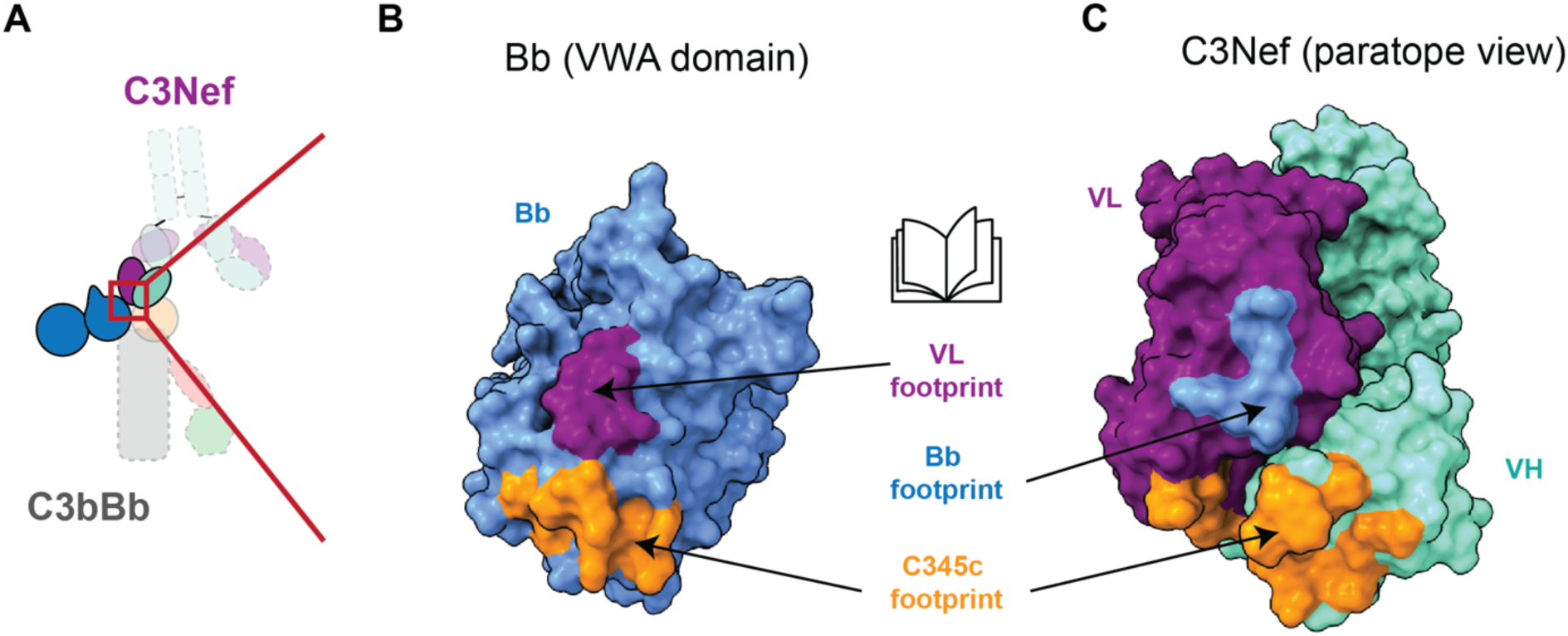
Variable light chain of C3Nef drives the interaction with factor B. **(A)** Diagrammatic representation of interactions between C3Nef VH and VL chains with C3b and Bb, respectively. (**B-C**) Open book model highlighting the interaction surfaces between the Von Williebrand type A (VWA) domain of Bb (**B**) and the CDRL2 region of the C3Nef paratope (**C**). Bb and Bb footprint in blue. VL and VL footprint in purple. VH domain in aquamarine. C345c footprints in orange. Note C345c domain makes 3-way contact with Bb, VH and VL domains. VH, variable heavy chain domain; VL, variable light chain domain.

**Fig. S22.**
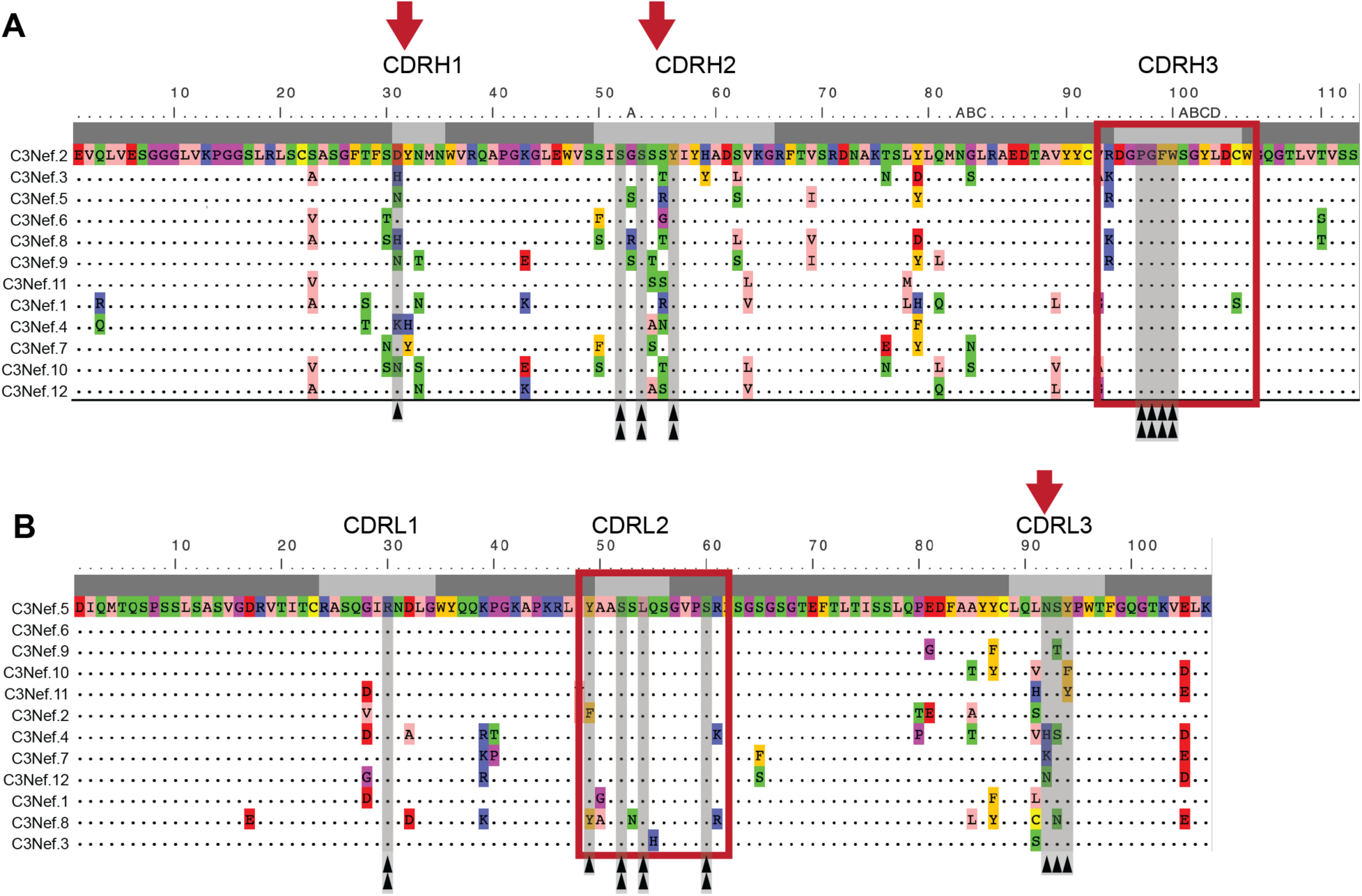
C3bBb contact residues are conserved across all 12 C3Nefs. (**A-B**) VH chain (**A**) and VL chain (**B**) sequence alignment for all 12 C3Nefs. Each dot represents a shared identity to the amino acid residue that precedes it vertically (in some cases, this is not always the top trace). Shaded horizontal grayscale bars (top) depict Kabat numbering for V(D)J with framework regions shown in dark gray and CDR regions shown in light gray. Narrow vertical gray bars with black arrows at the bottom indicate contact residues between C3Nef and the C345c or the Bb domain. Red boxes highlight CDR regions with high levels of sequence homogeneity. Red arrows highlight CDR regions with increased sequence heterogeneity. Double black arrows indicate C3bBb-contacting residues that are completely conserved across all 12 Nefs.

**Fig. S23.**
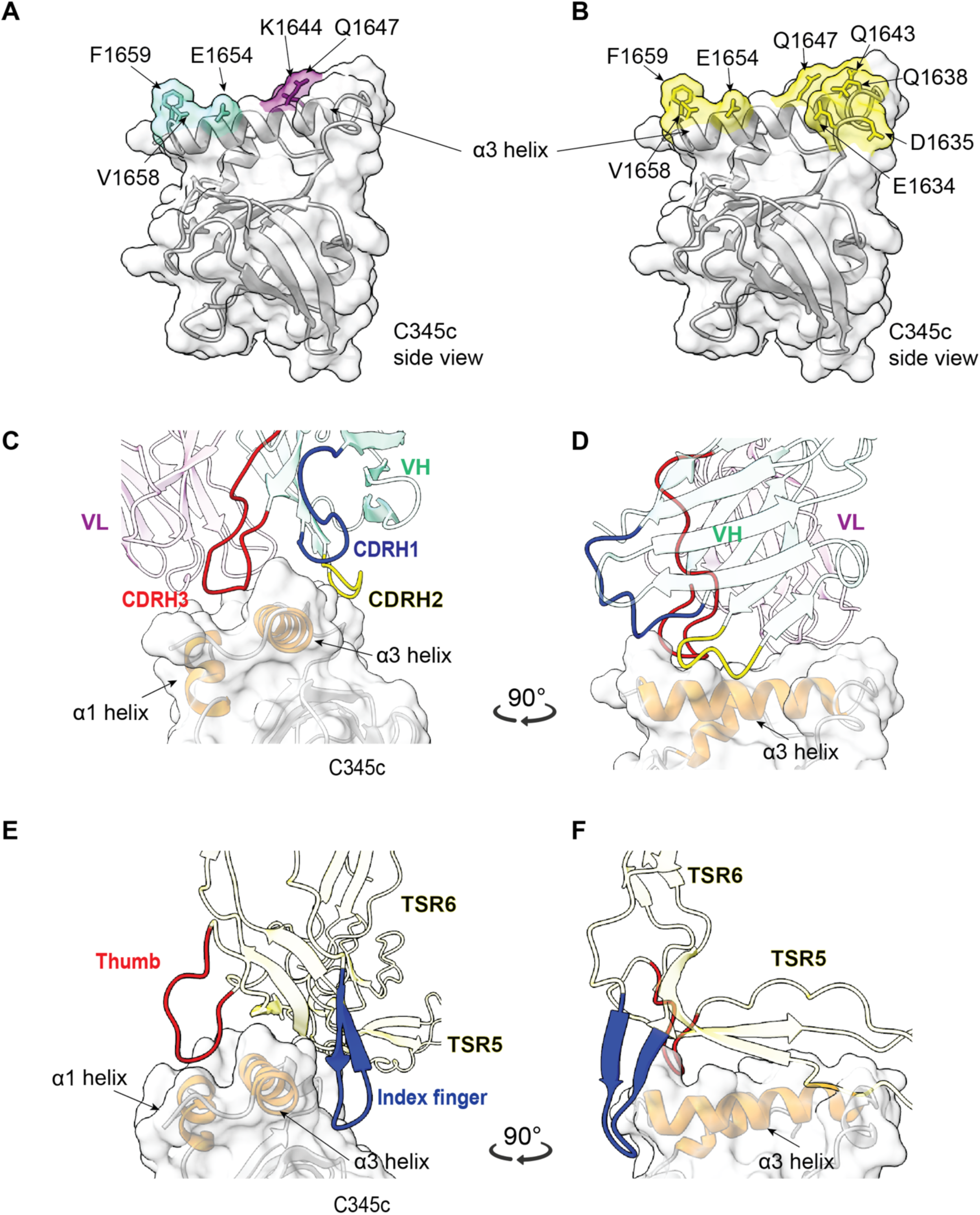
C3Nef and properdin binding sites overlap. (**A**) Side view showing the C3Nef footprint on the α3 helix of C3b’s C345c domain. VH in aquamarine, VL in purple. C345c domain in gray. Key contact residues are noted. (**B**) Same C345c domain as in (**A**) but showing the footprint of properdin (PDB:6RUR) in yellow. Important contact residues are noted. (**C-D**) Two angles showing the C3Nef CDR heavy regions 1-3 (blue, yellow, red, respectively) interacting with the α1 and α3 helix of C345c. (**E-F**) Same two angles of C345c highlighting the similarity in binding and location for the “thumb” (red) and “index finger” (blue) of properdin TSR5 and TSR6, respectively, compared with C3Nef above. C345c α1 and α3 helix highlighted in orange. VH, variable heavy domain; VL variable light domain; CDR, complementarity determining region; TSR, thrombospondin type 1 repeat domain.

**Fig. S24.**
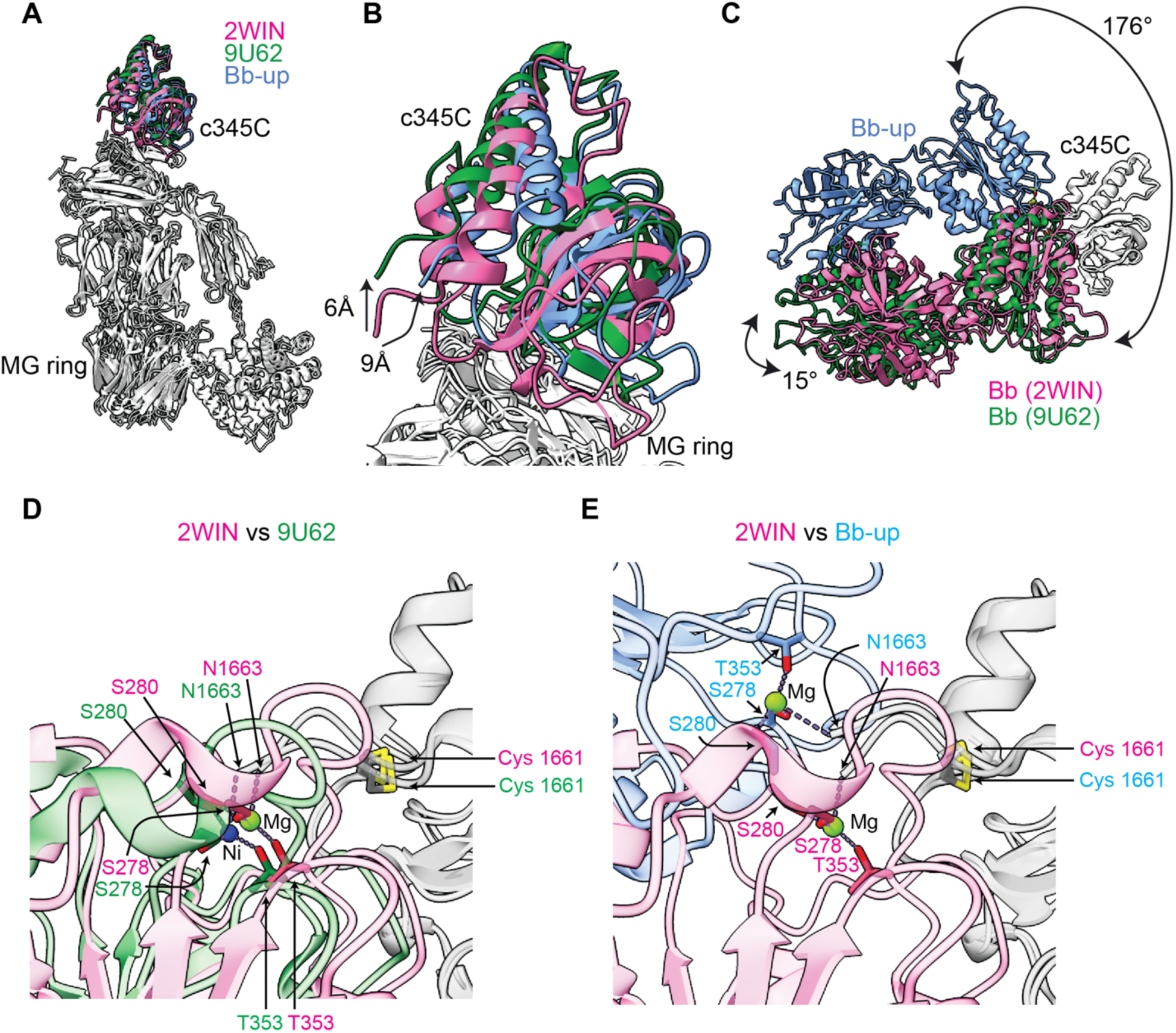
C345c and Bb orientations in Bb-up compared to known models (Bb-down). (**A**) C345c domain orientation relative to the MG ring upon convertase binding to substrate C3 or Nef. Superposition of C3bBb:C3Nef (Bb-up in blue), C3bBb(FP):C3 (green, PDB: 9U62), and SCIN-stabilized C3bBb (pink, PDB: 2WIN), aligned on the MG ring. (**B**) Zoomed-in view of panel A highlighting the different positions of C345c domain across structures with a focus on the α3 helix and C-terminus. (**C**) Bb re-orientation relative to the C345C domain of C3b upon convertase binding to substrate C3 (9U62) or Nef (Bb-up). Superposition of C3bBb-Nef (blue, Bb-up) and C3bBb-FP-C3 (green, PDB: 9U62) onto the C345C domain of SCIN-stabilized C3bBb (pink, PDB: 2WIN). (**D** and **E**) zoom-in view highlighting the MIDAS site in Bb domain re-orientations. Amino acids involved in Mg^2+^ coordination are listed.

**Fig. S25.**
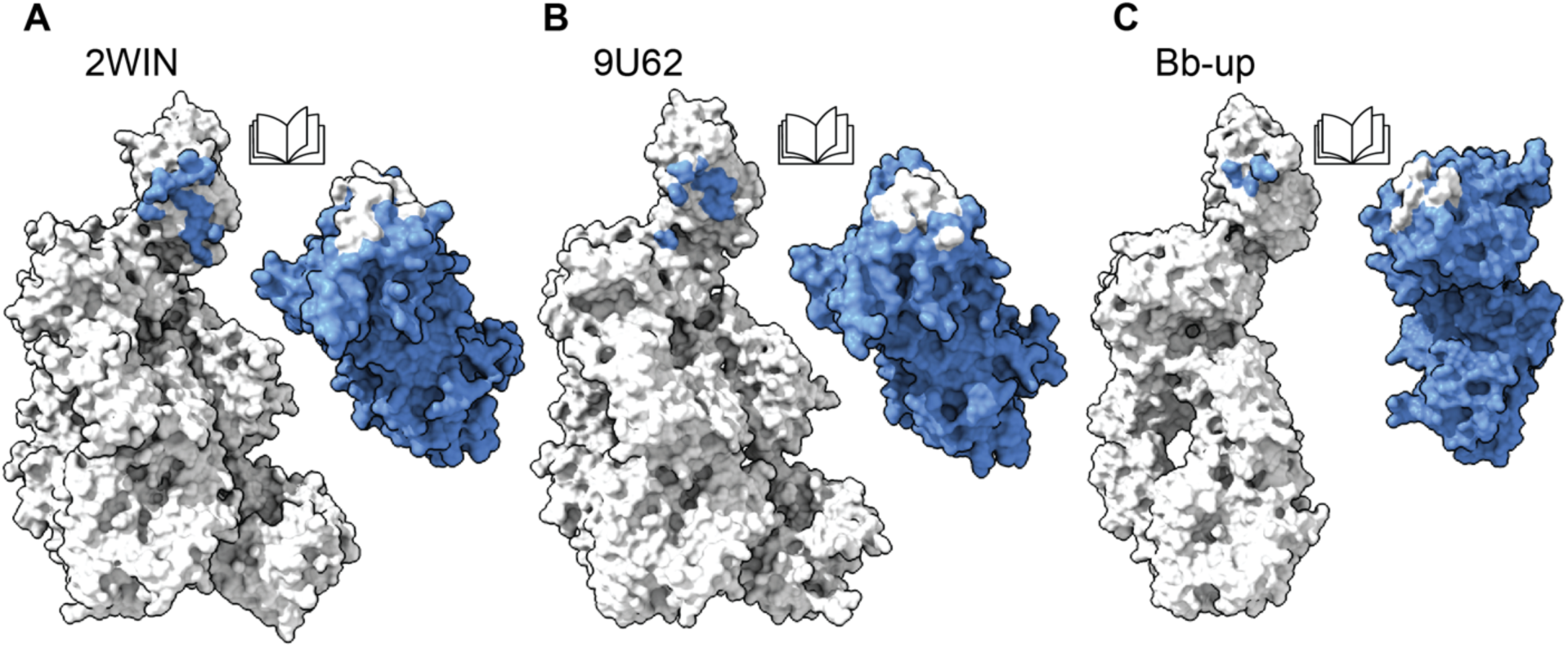
Bb-up configuration reduces contact between Bb and C345c domain. (**A**) Open book view of the contact footprints between C3b and Bb domains for SCIN-stabilized C3bBb (PDB:2WIN; Bb-down), (**B**) C3bBb-FP-C3 (PDB; 9U62; Bb-down) and (**C**) C3bBb-Nef (Bb-up).

**Fig. S26.**
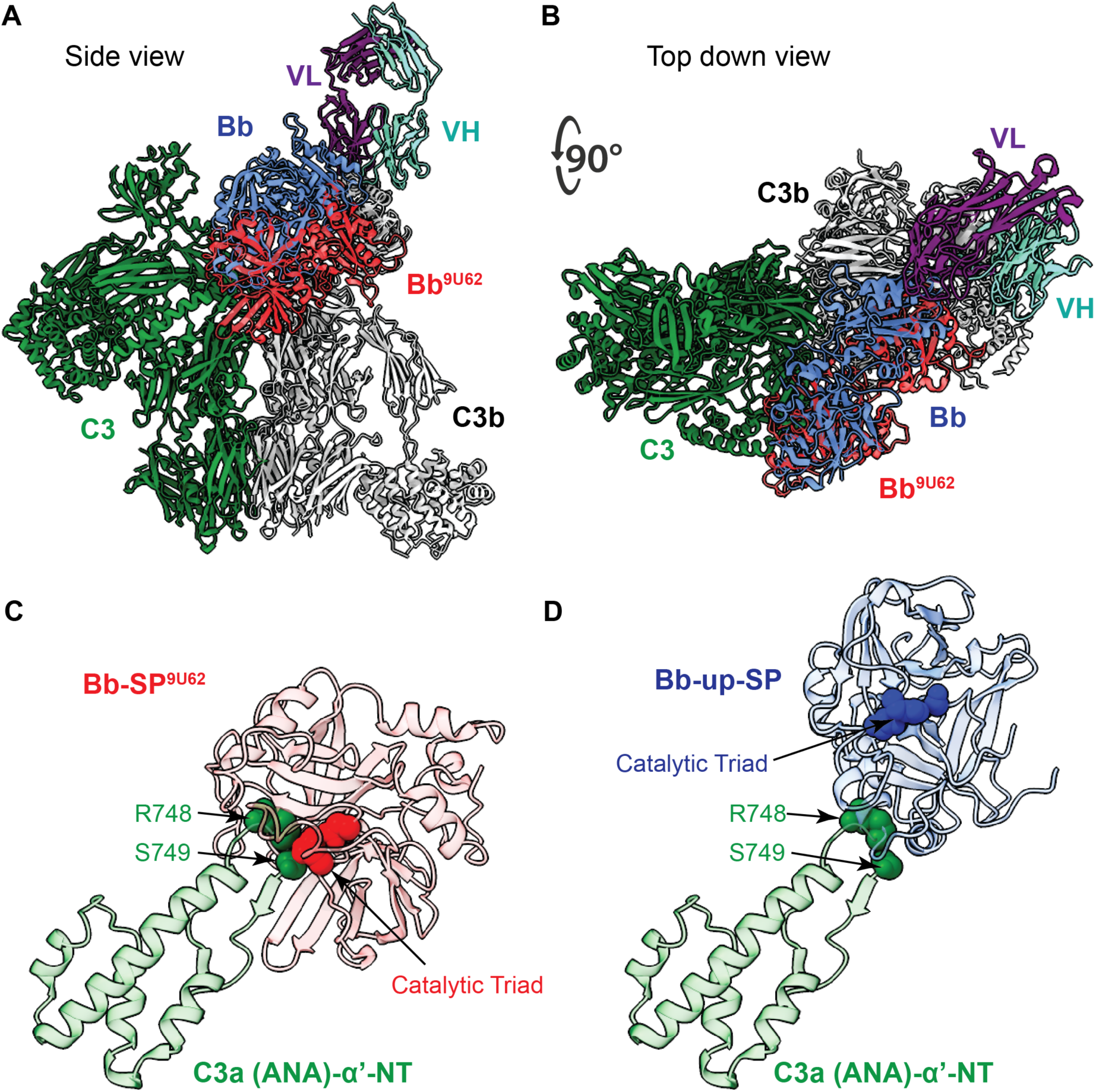
Structural models suggest C3Nef-mediated inhibition of C3bBb. (**A**) C3bBb-Nef (Bb in blue, VH in aquamarine, VL in purple, C3b in grey) superposed on the C3b chain of C3bBb-C3-FP (C3 in green, Bb in red, C3b in grey and FP not shown in the figure). (**B**) Rotation of 90 degrees showing the top view. (**C**) Zoom-in view of the interaction between the catalytical triad in Bb-SP and the C3 scissile loop in 9U62. (**D**) Zoom-in view of the distance between the catalytical triad in Bb-SP and the C3 scissile loop in Bb-up.

**Fig. S27.**
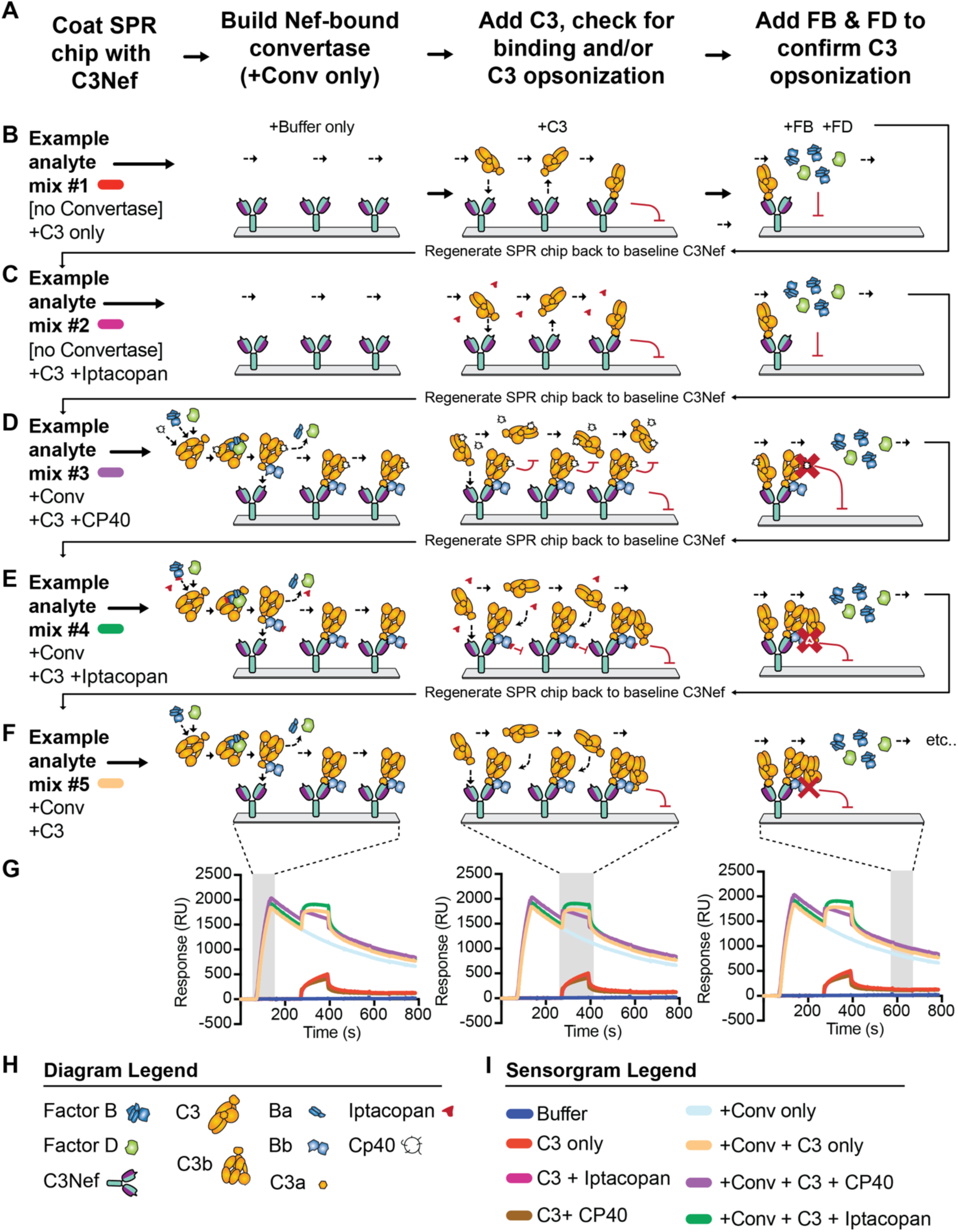
Workflow and diagram of surface plasmon resonance (SPR) experiments used to characterize binding to, or cleavage of, free C3 by C3Nef-stabilized C3bBb convertase. **(A)** Experimental workflow performed in iterative cycles as described in Methods. (**B-F**) Diagram depicting 5 different cycle conditions tested in assay (not all cycle conditions tested are shown in diagram). (**G**) Representative sensorgram overlays. Shaded gray windows correspond to the phase depicted diagrammatically above each sensorgram. (**H**) Diagram legend. (**I**) Sensorgram legend listing all conditions tested on C3Nef-coated SPR chip. FP, factor P properdin; FB, factor B; FD, factor D. Conv; convertase; RU, resonance units. “+Conv” denotes an analyte mixture of +C3b, +FB, and +FD.

**Fig. S28.**
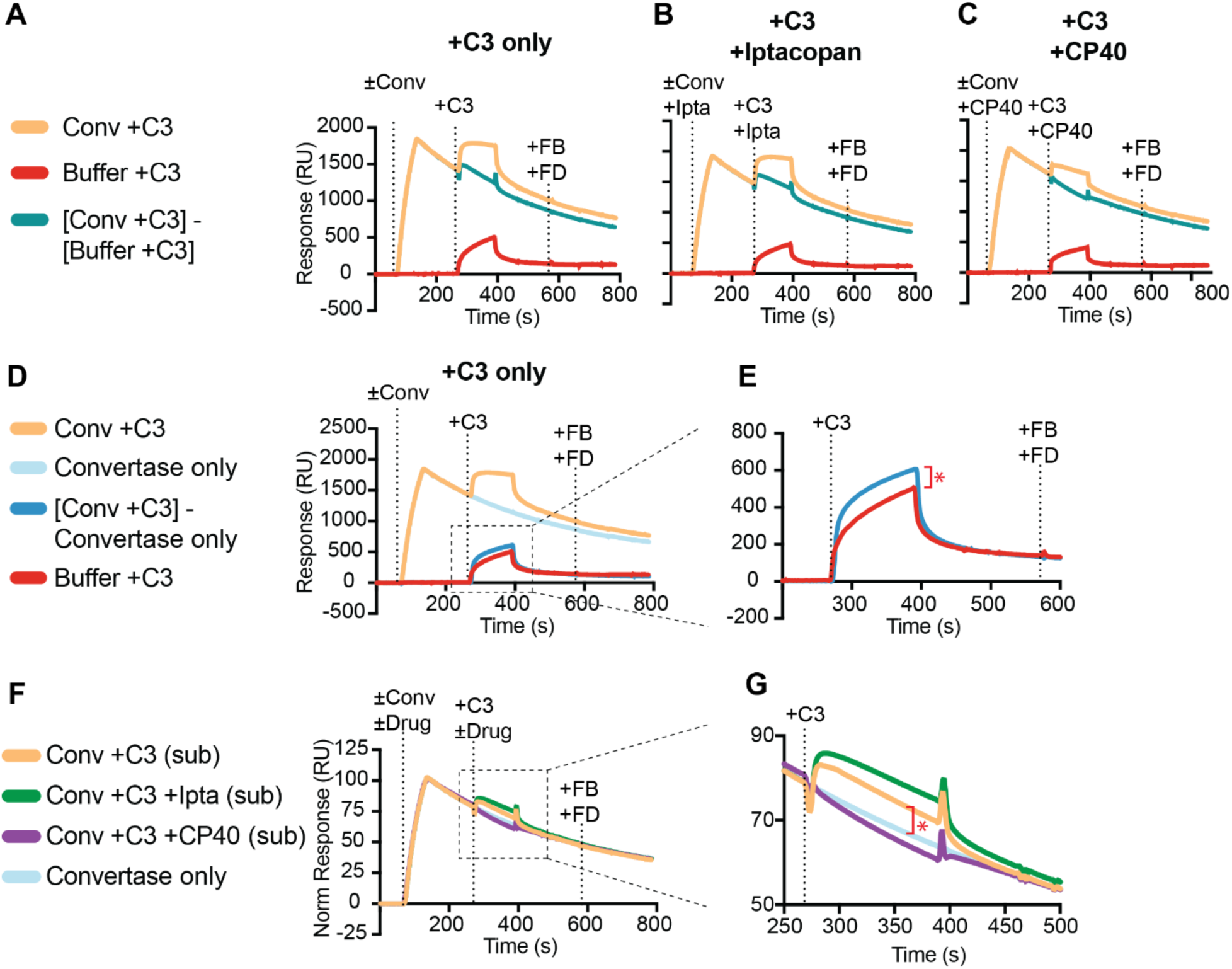
C3Nef-bound C3bBb convertase binds but does not cleave free C3. (**A**) SPR sensorgram overlays showing the amount of C3 binding to C3bBb convertase that is stabilized by C3Nef (teal trace). Control cycles used to calculate C3 binding to C3bBb: Red trace captures background low-level binding of C3 to the variable domains of C3Nef; tan trace captures the combined binding of both C3 to C3bBb convertase stabilized by C3Nef in addition to C3 binding to C3Nef variable domains alone. (**B and C**) Same as (**A**), but with the addition of complement inhibitors iptacopan (Ipta) and CP40, which were added to the analyte mixtures prior to convertase-forming injection and free C3 injection steps, respectively. (**D**), same as in (**A**), but showing the convertase-only control (light blue) and the [Conv+C3] - [Conv-only] trace (dark blue). (**E**) Zoomed-in region showing the increase in binding when comparing +C3 only (red) to convertase-subtracted C3bBb+C3 (dark blue). Red asterisk denotes the calculated fraction of C3 that binds specifically to C3Nef-stabilized C3bBb. (**F**) C3-only-subtracted traces showing C3 binding to C3Nef-stabilized C3bBb for C3 (tan) and also C3 + iptacopan (green), but not for C3 + CP40 (purple). (**G**) Zoomed-in region; red asterisk denotes the calculated fraction of C3 that binds specifically to C3Nef-stabilized C3bBb. FB, factor B; FD, factor D; Ipta, iptacopan. “Conv” is an analyte mixture of +C3b, +FB, and +FD. See Methods for details.

**Fig. S29.**
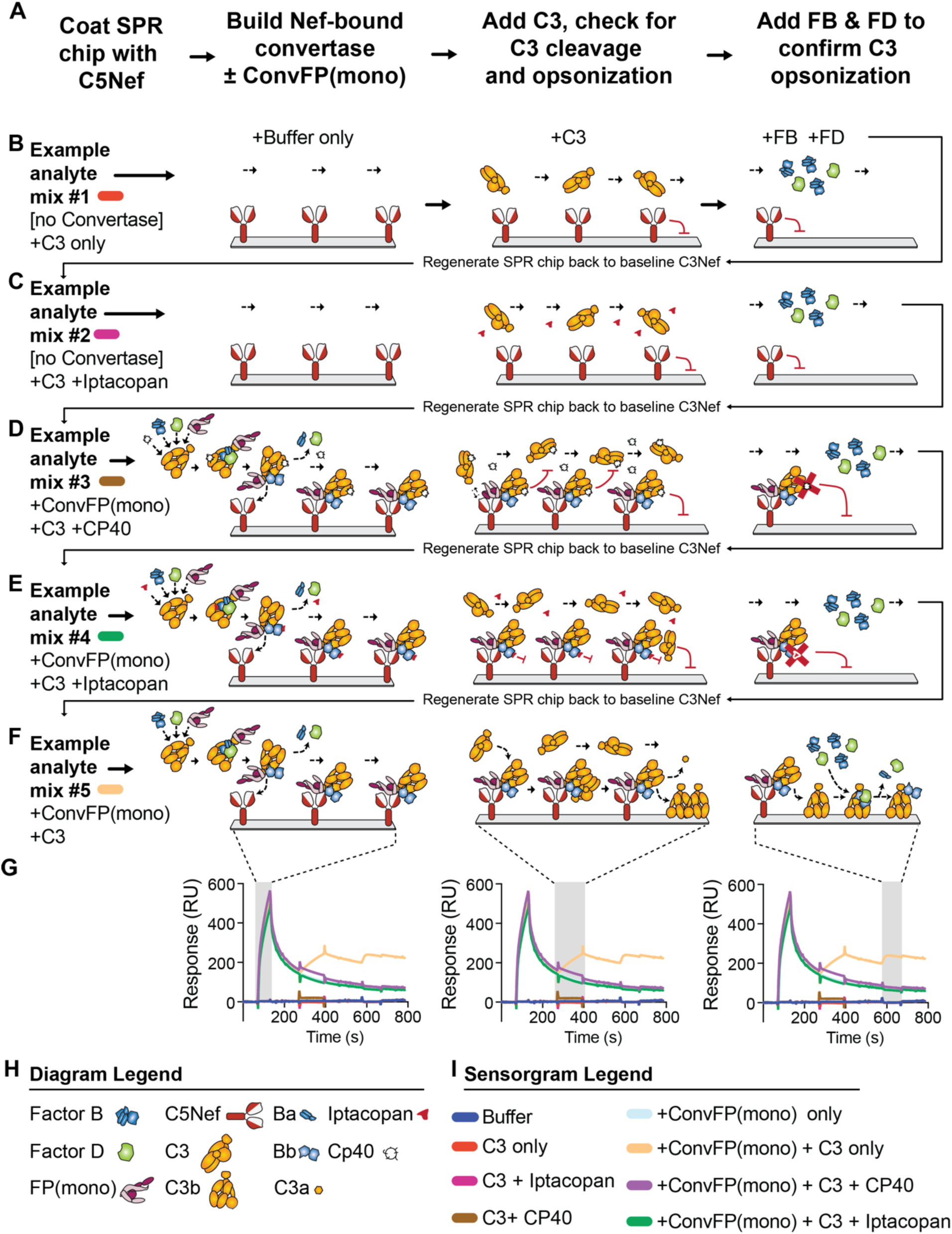
Workflow and diagram of surface plasmon resonance (SPR) experiments used to characterize cleavage of C3 by C5Nef-bound C3bBbP(mono) convertase. (**A**) Experimental workflow performed in iterative cycles. (**B-F**) Diagram depicting 5 different cycle conditions tested in assay (not all cycle conditions tested are shown in diagram). (**G**) Representative sensorgram overlays. Shaded gray windows correspond to the phase depicted diagrammatically above each sensorgram. (**H**) Diagram legend. FP, factor P / properdin. (**I**) Sensorgram legend listing all conditions tested on C3Nef-coated SPR chip. FP(mono); monomeric factor P / properdin; FB, factor B; FD, factor D. Conv; C3bBb convertase. “+Conv” is a mixture that includes C3b, FB, FD, and FP(mono).

**Fig. S30.**
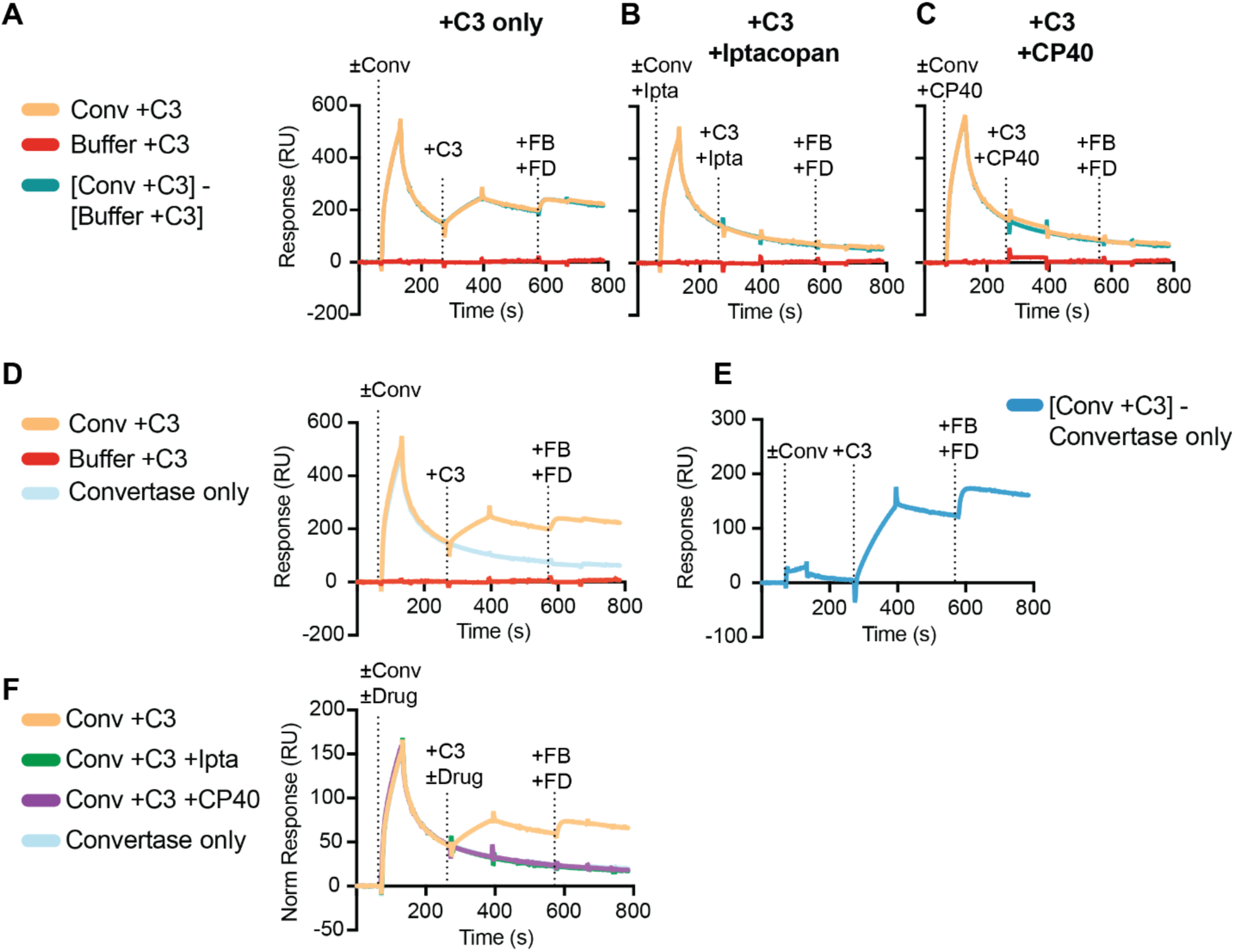
C5Nef-bound C3bBbP(mono) convertases cleave C3. (**A**) SPR sensorgram overlay showing C3 cleavage by C5Nef-stabilized C3bBbP(mono) convertases (tan and teal traces) and subsequent opsonization of C3b onto SPR chip (verified by +FB +FD addition). Control trace (red) showing non-reactivity of C3 to C5Nef on SPR chip. (**B and C**). Same as (**A**), but with the addition of complement inhibitors iptacopan (Ipta) and CP40 added to the first two injection steps. Both iptacopan and CP40 effectively block C3 cleavage and C3b opsonization. (**D**), same as in (**A**), but showing the convertase-only control (light blue). (**E**) Convertase-subtracted trace (dark blue) showing simultaneous C3 cleavage and C3b opsonization (+C3) that supports subsequent C3bBb formation following the injection of +FB and +FD. (**F**) Normalized traces inhibitory effects of iptacopan (green) and CP40 (purple) on C3 cleavage and C3b deposition. “+Conv” is a mixture of +C3b, +FB, +FD, and +FP(mono).

**Fig. S31.**
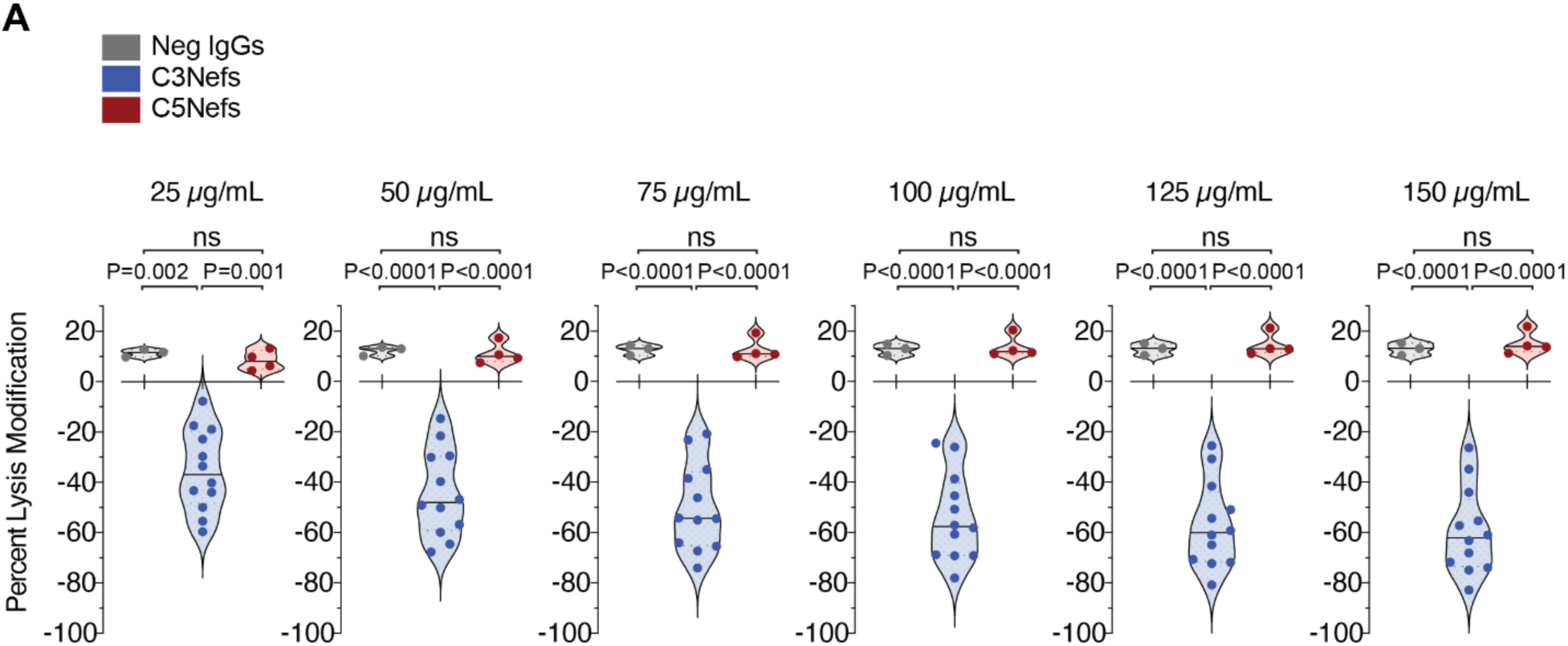
Guinea pig red blood cell lysis is inhibited by C3Nefs but not C5Nefs. (**A**) Guinea Pig RBCs mixed with equal amounts of NHS and increasing concentrations of C3Nef (blue) or C5Nef (red) or Neg IgG (gray). Changes in baseline lysis shown on Y axis. P values determined by multiple comparison ordinary one-way ANOVA.

**Fig. S32.**
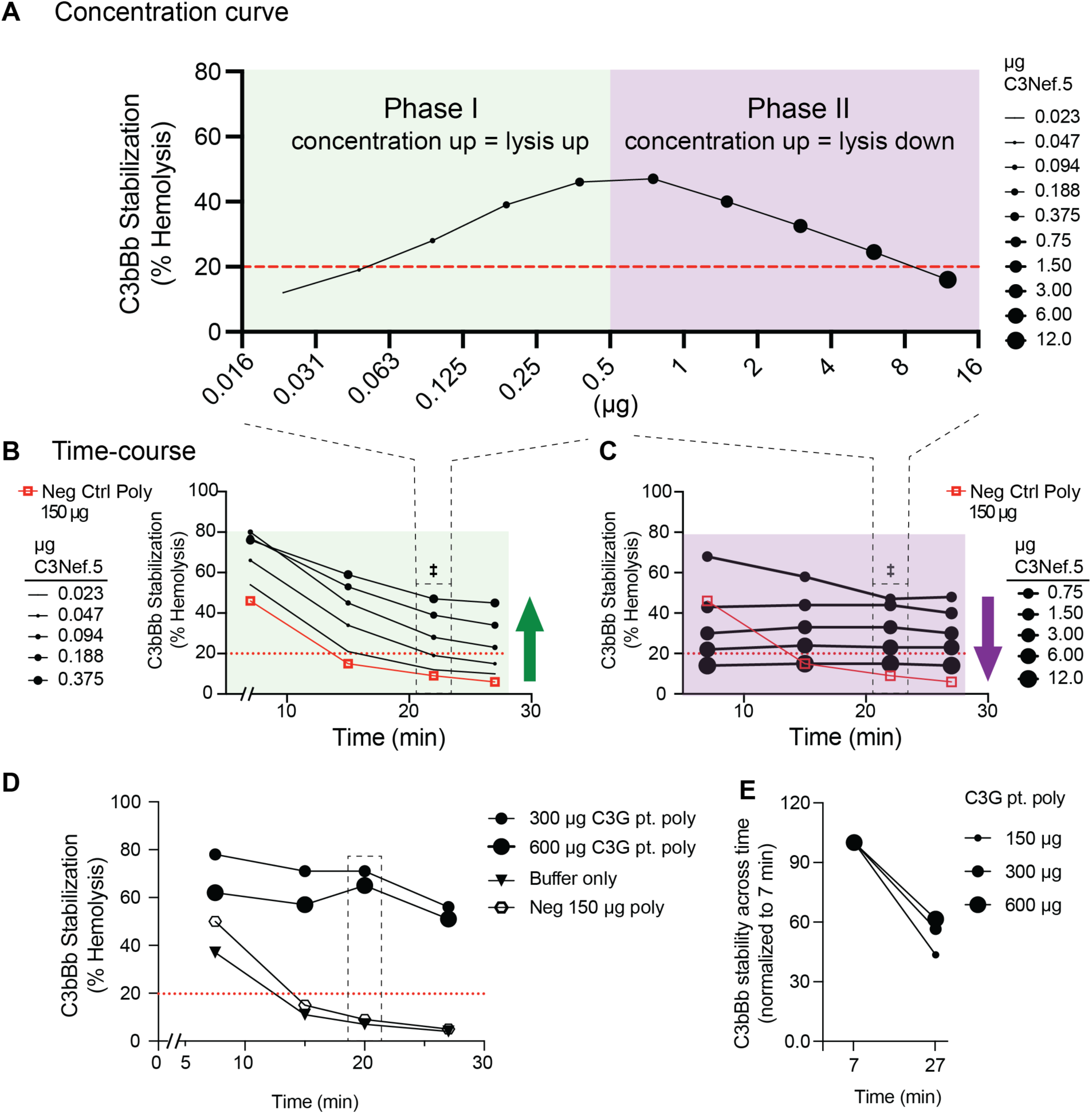
Effects of concentration and time on C3Nef activity. (**A**) Hemolytic C3bBb stabilization assay data showing concentration effects for monoclonal C3Nef.5 across a 500-fold range. Increased C3Nef.5 concentration results in increased hemolysis up to 0.5µg (“Phase I”), after which further increases in C3Nef.5 concentration result in decreased hemolysis (“Phase II”). (**B-C**) Time course overlay showing every C3Nef.5 concentration and timepoint for Phase I (**B**) and Phase II (**C**) tested. (**D**) Same assay as (**B-C**), but for C3G index patient polyclonal IgGs showing greater convertase stabilization and reduced hemolysis with increased polyclonal IgG concentrations. Dashed box denotes % Hemolysis at the 20 min timepoint reported out clinically. (**E**) Increases in polyclonal IgG from C3G index patient results in increased C3bBb stability across time.

**Fig. S33.**
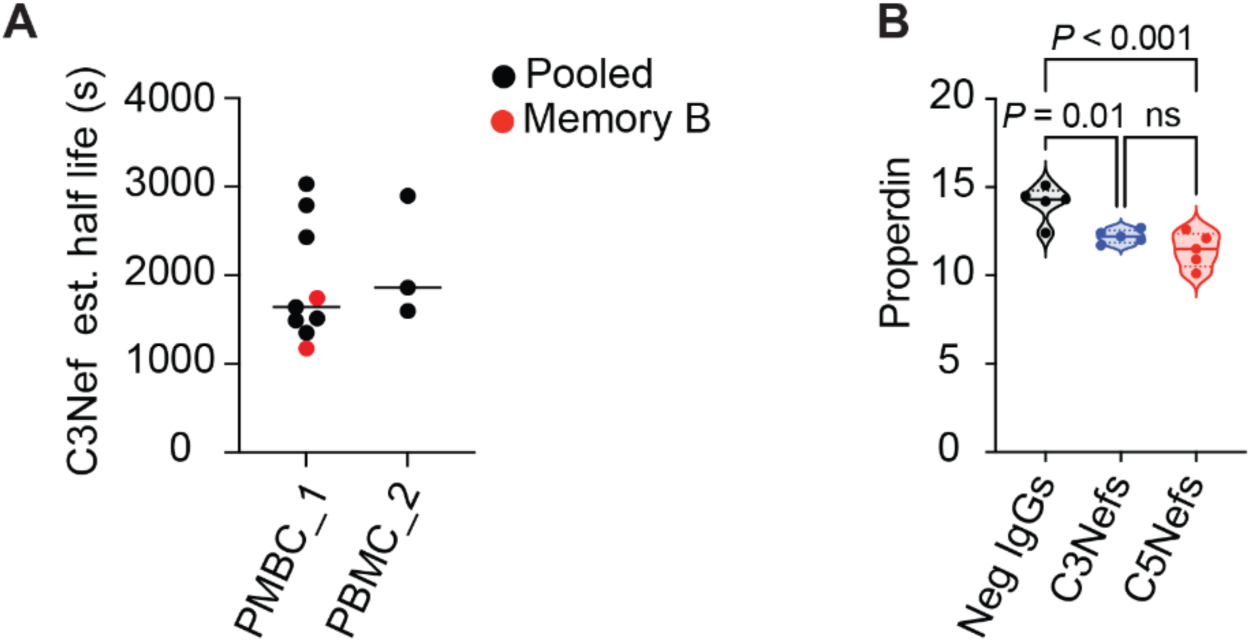
C3Nef estimated half-life and Nef effects on properdin levels. (**A**) Estimated half-life values for C3Nefs grouped by collection timepoint. (**B**) Changes in properdin levels in normal human serum spiked with Neg IgG, C3Nefs, or C5Nefs. Ordinary one-way ANOVA.

**Fig. S34.**
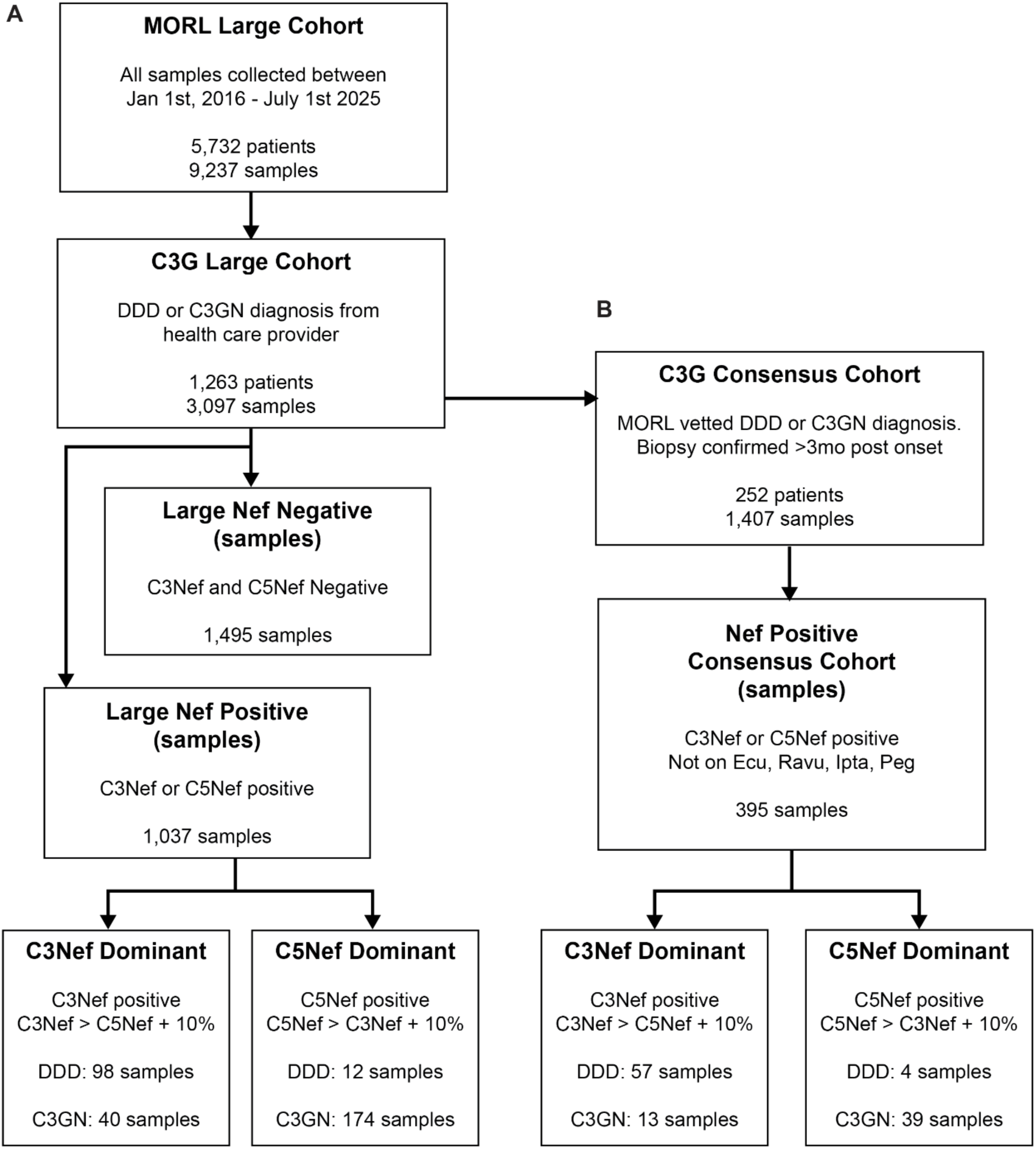
C3G patient datasets. (**A**) C3G Large Cohort data filtering flowchart. (**B**) C3G Consensus Cohort data filtering flowchart. MORL; University of Iowa Molecular Otolaryngology and Renal Research Laboratories, DDD; dense deposit disease, C3GN; C3 glomerulonephritis, Ecu; Eculizumab, Ravu; Ravulizumab, Ipta; iptacopan, Peg; Pegcetacoplan.

**Fig. S35.**
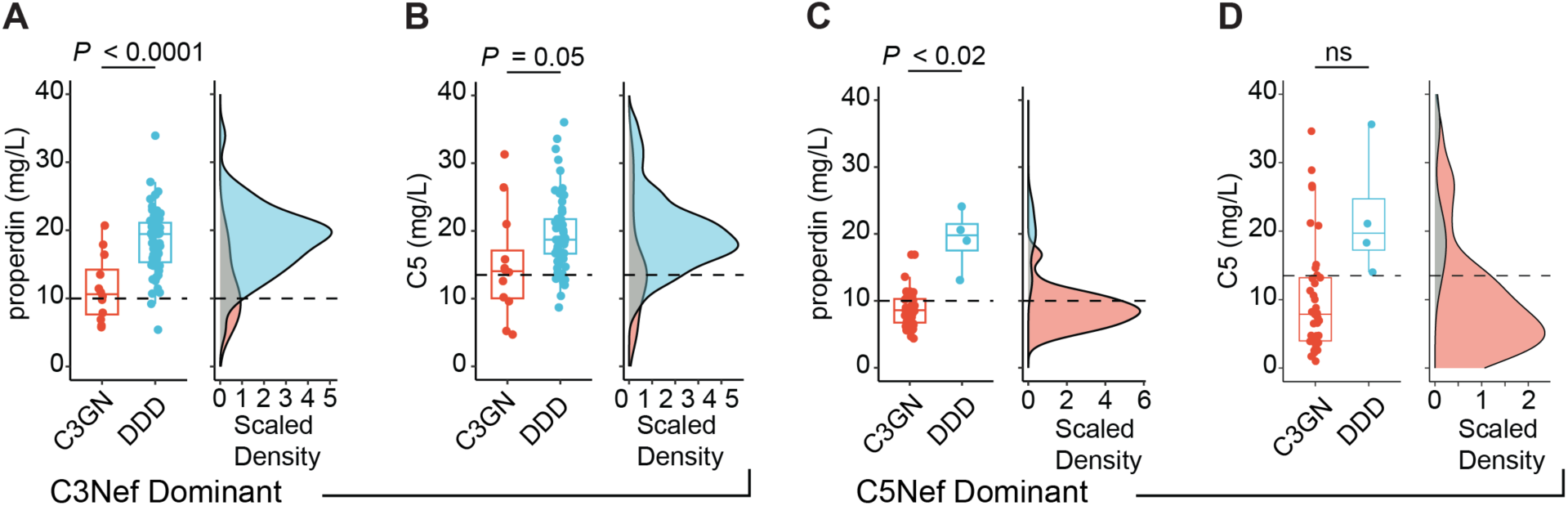
Filtering on Nef dominance separates DDD from C3GN samples. (**A-B**) Properdin and C5 levels for C3Nef-dominant C3G Consensus Cohort patient samples. (**C-D**) Properdin and C5 levels for C5Nef-dominant C3G Consensus Cohort patient samples. DDD; dense deposit disease, C3GN; C3 glomerulonephritis. P values listed for Student’s t-test.

**Fig. S36.**
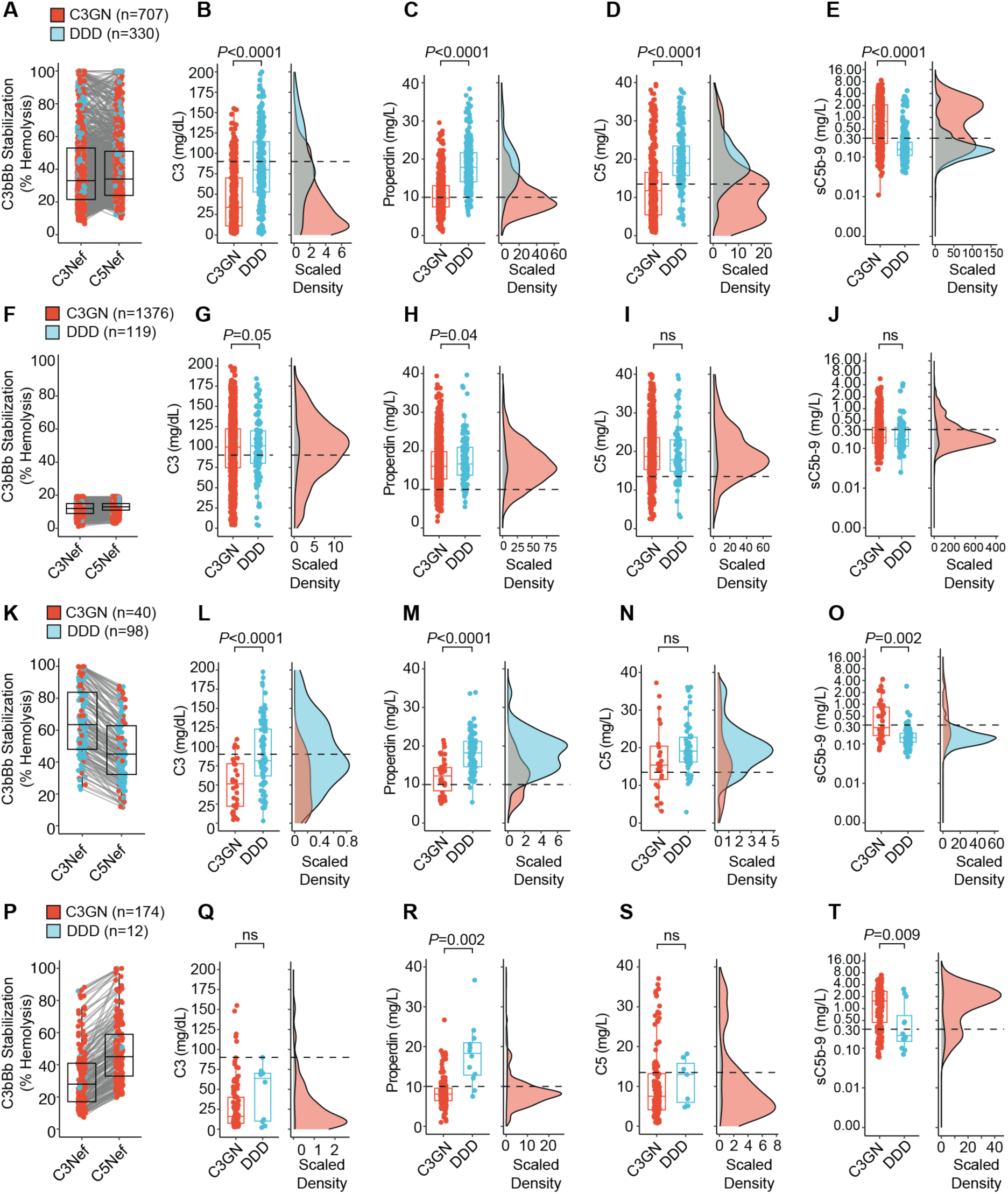
Biomarker signatures differ between DDD and C3GN patients. (**A-D**) C3G Large Cohort patient samples filtered on C3Nef or C5Nef positivity (**A**) and subsequently separated into their provider-assigned DDD or C3GN diagnoses, then examined for levels of C3 (**B**), properdin (**C**), C5 (**D**), and sC5b9 (**E**). For each biomarker a boxplot and the corresponding scaled density histogram are displayed. P values denote Student’s t test. (**F-J**) Same as **A-D** but filtered on Nef-negative samples. (**K-O**) Same as **A-D** but filtered on C3Nef-dominant samples. (C3Nef-positive, and C3Nef > C5Nef + 10%) (**P-T**) Same as **A-D**, but for C5Nef-dominant samples. (C5Nef-positive, and C5Nef > C5Nef + 10%).

**Fig. S37.**
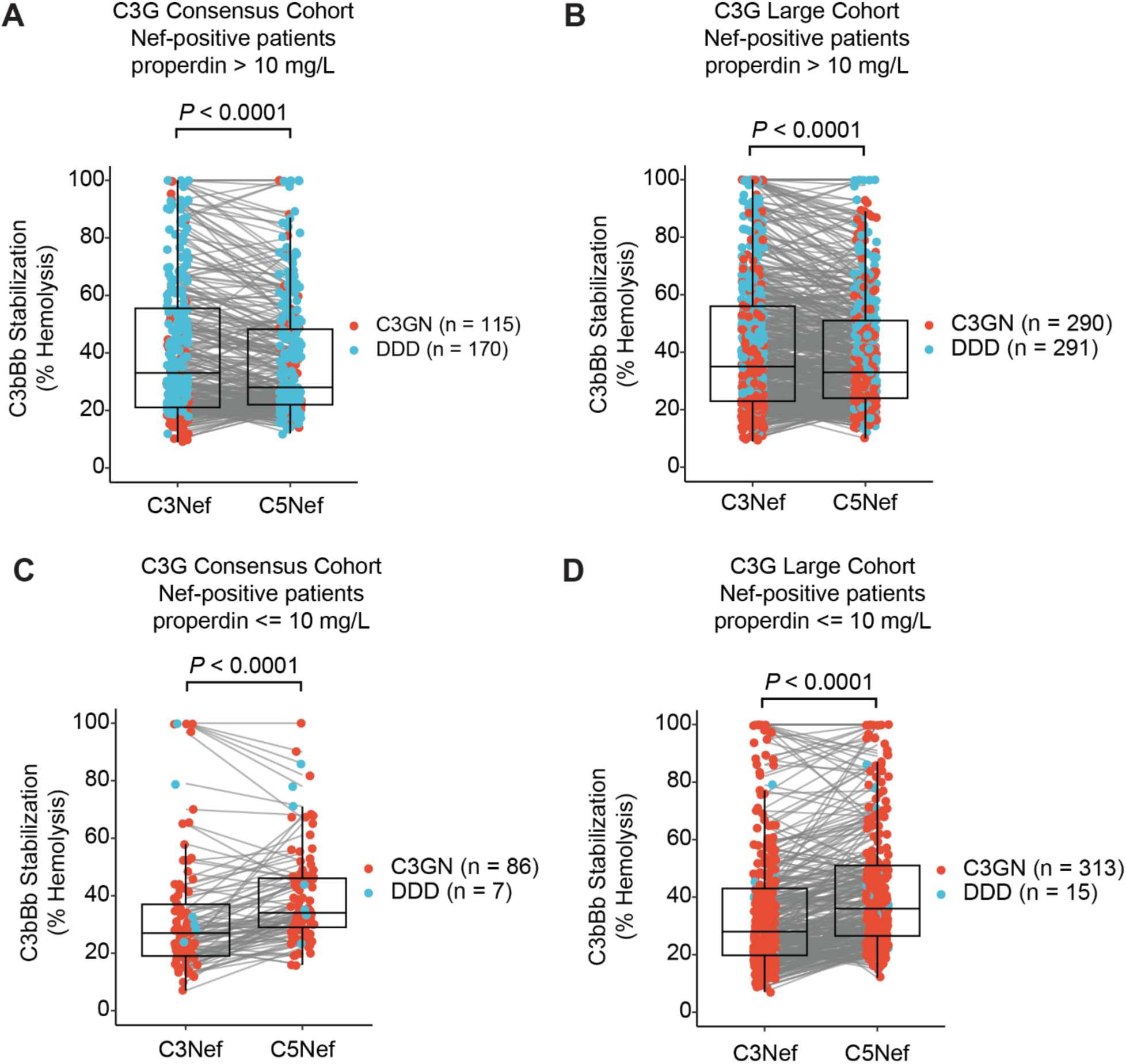
Effect of properdin levels on Nef-dominance in DDD vs. C3GN patients. (**A-B**) Paired C3Nef and C5Nef levels for C3G Cohort patient samples filtered for normal properdin levels (>10mg/L) by the Consensus Cohort (A), or Large Cohort (B) groups. After filtering, samples are colored by disease subtype. (**C-D**) Same as **A-B** but using samples with clinically low levels of properdin (<10mg/L). Of note, the vast majority of DDD samples in both cohorts are enriched in the properdin normal group. P values denote Student’s t test.

**Fig. S38.**
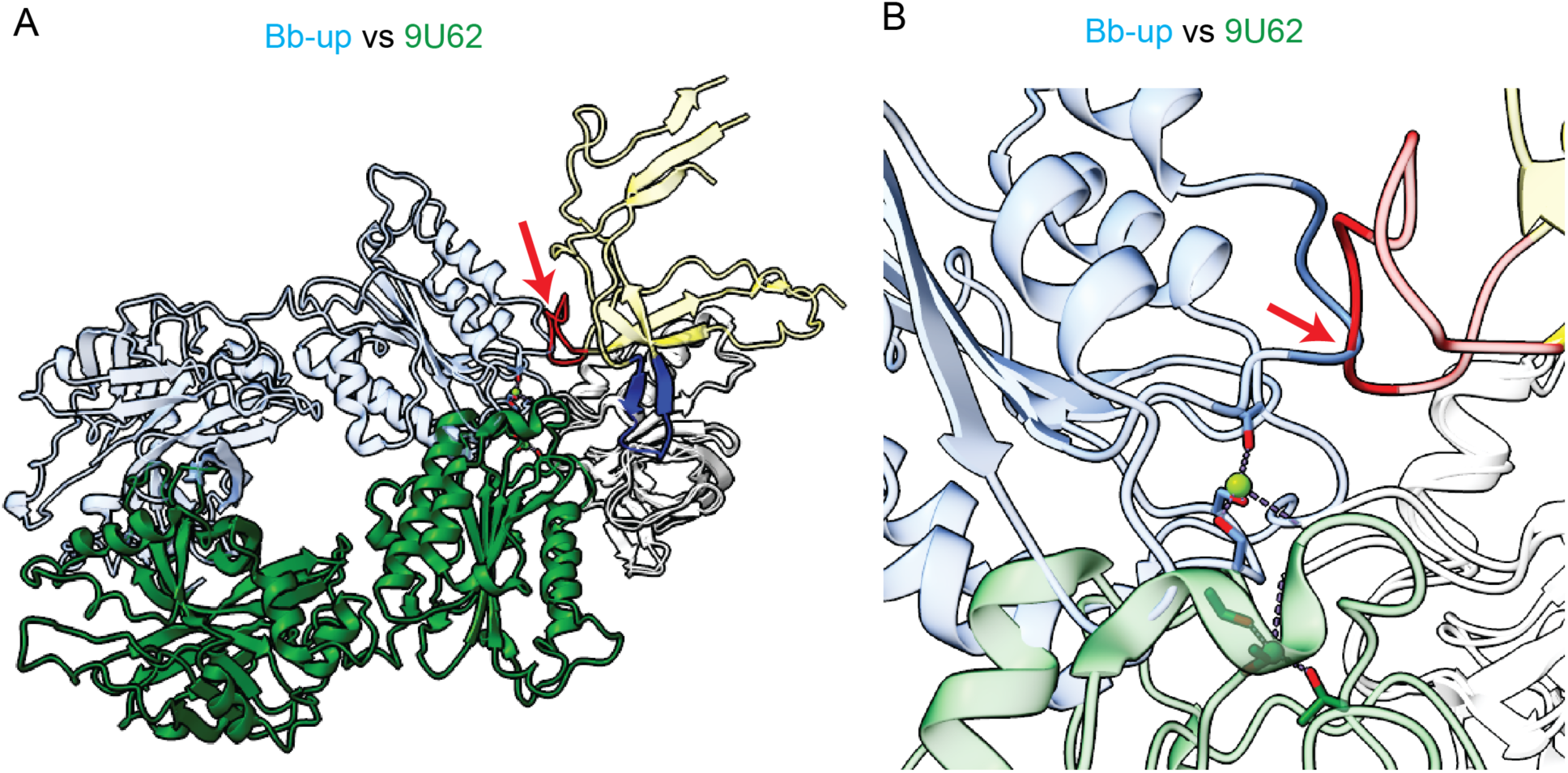
Properdin clashes with Bb-up and sterically hinders rotation around the MIDAS site. (**A**) Bb from Bb-up (blue) overlayed onto Bb from 9U62 (green, Bb-down) aligned on C345c domain. Properdin from 9U62 model shown in yellow. Finger and thumb of properdin shown in blue and red, respectively. (B) Zoomed in model near the MIDAS region showing steric clash of Bb from Bb-up (dark blue) and the thumb (red) of properdin from 9U62.

**Fig. S39.**
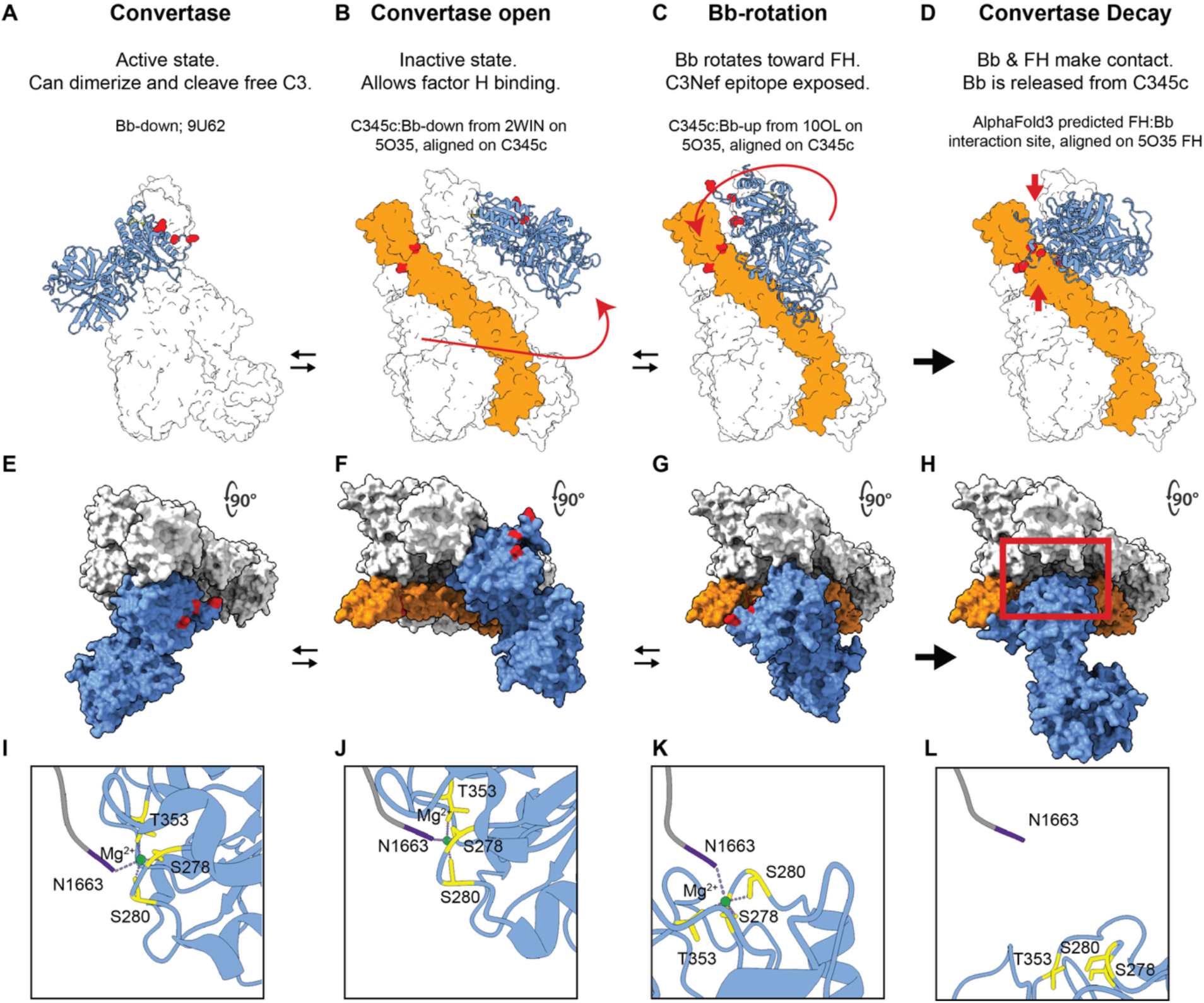
Bb-up rotational model of C3bBb decay. (**A-D**) Surface area models depicting a series of hypothetical progression events leading to Bb displacement from C3bBb by factor H. The main events for each stage are shown on top. The various structures and alignments used to generate each model are also listed. C3b in white. Bb in blue. Factor H in orange. Red dots indicate known factor H variants [FH(p.R53C), FH(p.Q81P)] and factor B variants [FB(p.K323E), FB(p.K323Q), FB(p.S367R), FB(p.D371G)] that are associated with impaired decay, but which lack known mechanisms. Red arrows indicate hypothetical Bb domain movement and proposed rotation. Black arrows depict progression direction. (**E-H**) Top-down view of **A-D** highlighting the predicted lateral and rotational movement of Bb relative to the C3b:FH complex. (**H**) Top-down view of AlphaFold3 model of a predicted Bb:FH(1-6) interaction that is overlayed onto the 5O35 model aligned on factor H. The red box highlights the predicted physical separation of Bb from C345c. (**I-K**) Zoomed in ribbon models showing the integrity of the MIDAS region for each structure shown above. In the final step Bb is physically separated from C345c resulting in disruption of the MIDAS coordinating residues.

**Fig. S40.**
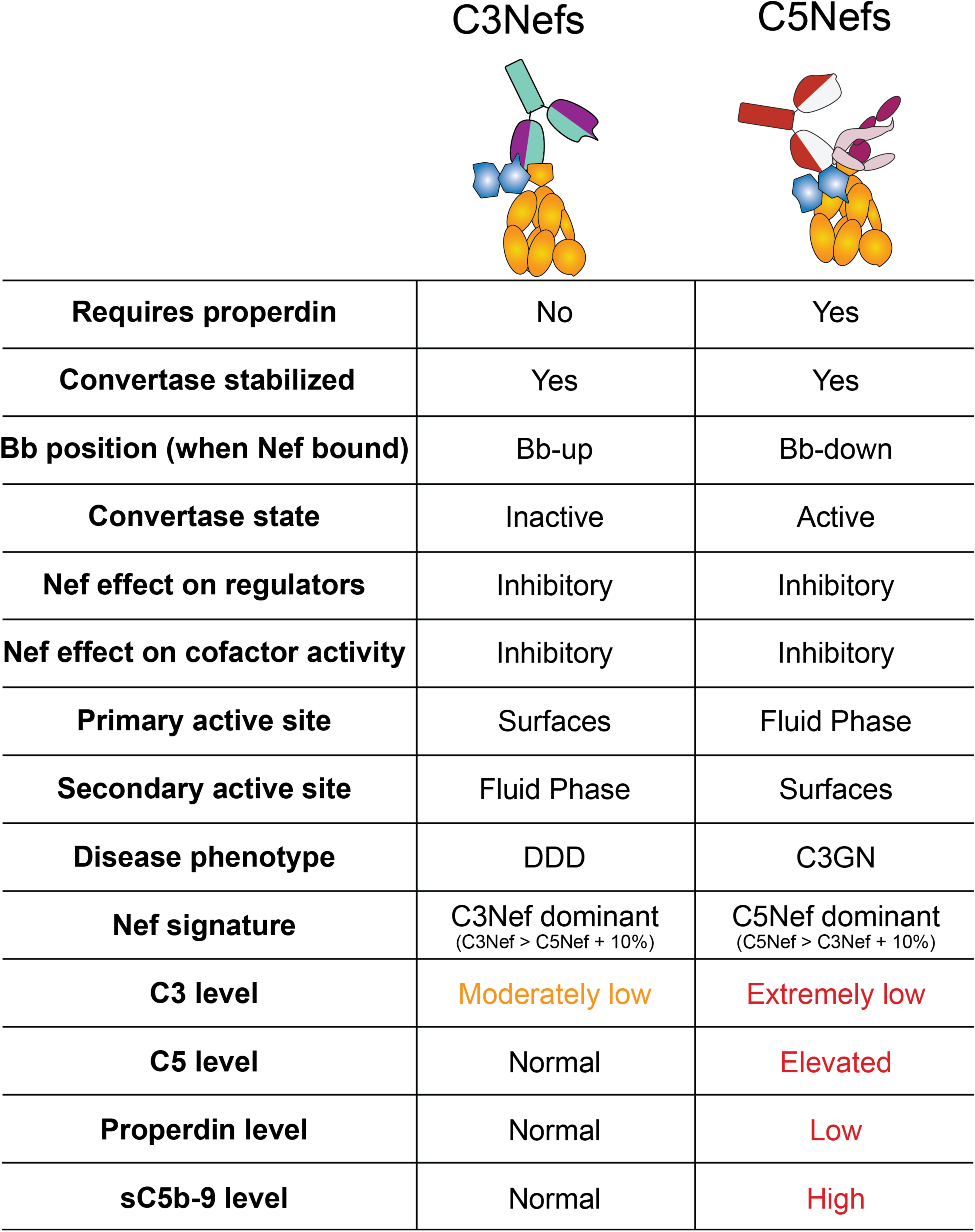
Comparison of C3Nefs to C5Nefs. Biochemical, structural, functional, and clinical biomarker profiles associated with C3Nefs (left) as compared with C5Nefs (right).

**Table S1.** Reagents used for FACS enrichment of B lineage PBMCs.

| **Antigen or Marker** | **Fluorophore** | **Dilution** | **Catalog no.** | **Clone** | **Isotype** | **Supplier** | **Description** |
| --- | --- | --- | --- | --- | --- | --- | --- |
| FC block | NA | 1:40 | 5642219 | Fc1 | human IgG1 | BD Pharmingen | FC Block |
| CD2 | FITC | 1:50 | 300206 | RPA-2.10 | mouse IgG1, k | Biolegend | Dump: T cells, NK cells |
| CD3 | FITC | 1:100 | 317305 | OKT3 | mouse IgG2a, k | Biolegend | Dump: T cells |
| CD14 | FITC | 1:100 | 301803 | M5E2 | mouse IgG2a, k | Biolegend | Dump: monocytes, macrophages |
| CD16 | FITC | 1:100 | 302005 | 3G8 | mouse IgG1, k | Biolegend | Dump: NK, monocytes, macrophages |
| CD36 | FITC | 1:100 | 336203 | 5-271 | mouse IgG2a, k | Biolegend | Dump: erythrocytes, platelets, monocytes, macrophages |
| Annexin V | eFluor450 | 1:20 | 48-8006-69 | NA | NA | Invitrogen | Live Dead |
| 7AAD | NA | 1:50 | 00-6993-42 | NA | NA | Invitrogen | Live Dead |
| CD19 | BV711 | 1:100 | 302245 | HIB19 | mouse IgG1, k | Biolegend | B cells |
| CD27 | BV605 | 1:100 | 302829 | 323 | mouse IgG1, k | Biolegend | Activated B cells |
| CD38 | APC | 1:100 | 356605 | HB-7 | mouse IgG1, k | Biolegend | Plasma cells, Plasmablast subset |
| CD21 | PE | 1:100 | 354903 | Bu32 | mouse IgG1, k | Biolegend | B cell subset |
| IgD | APCFire750 | 1:100 | 348237 | IA6-2 | mouse IgG2a, k | Biolegend | B cell subset |

**Table S2.**
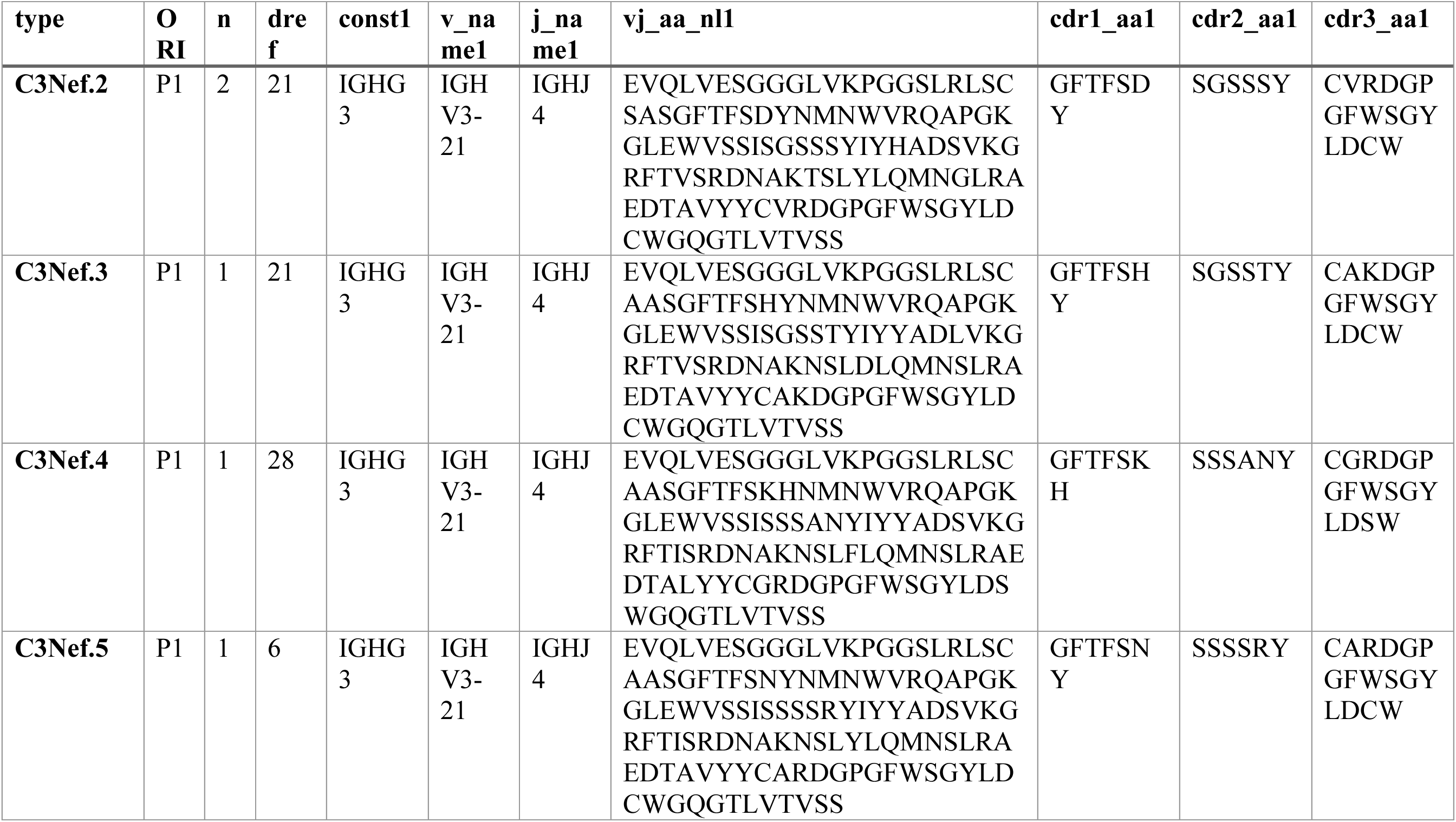
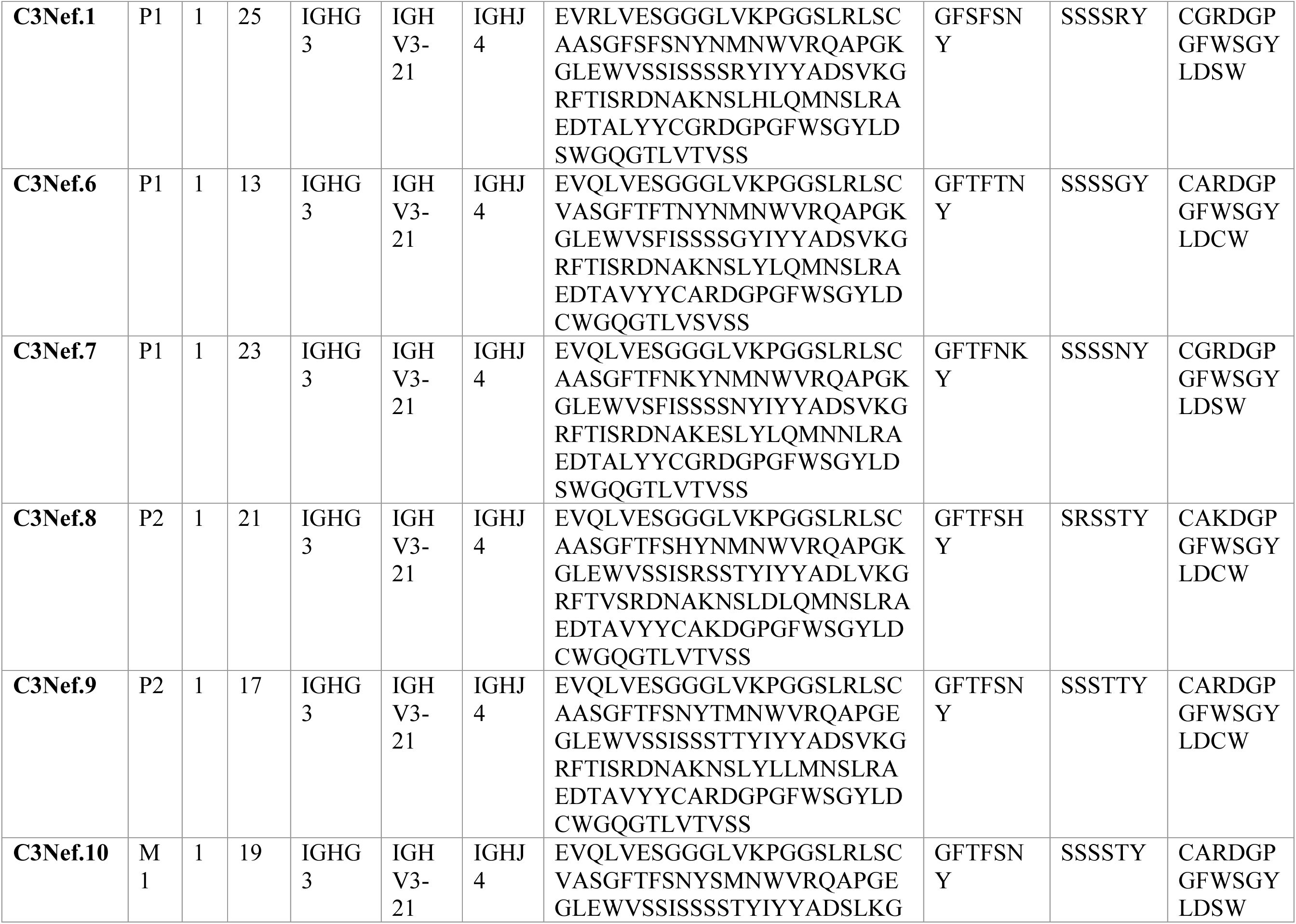
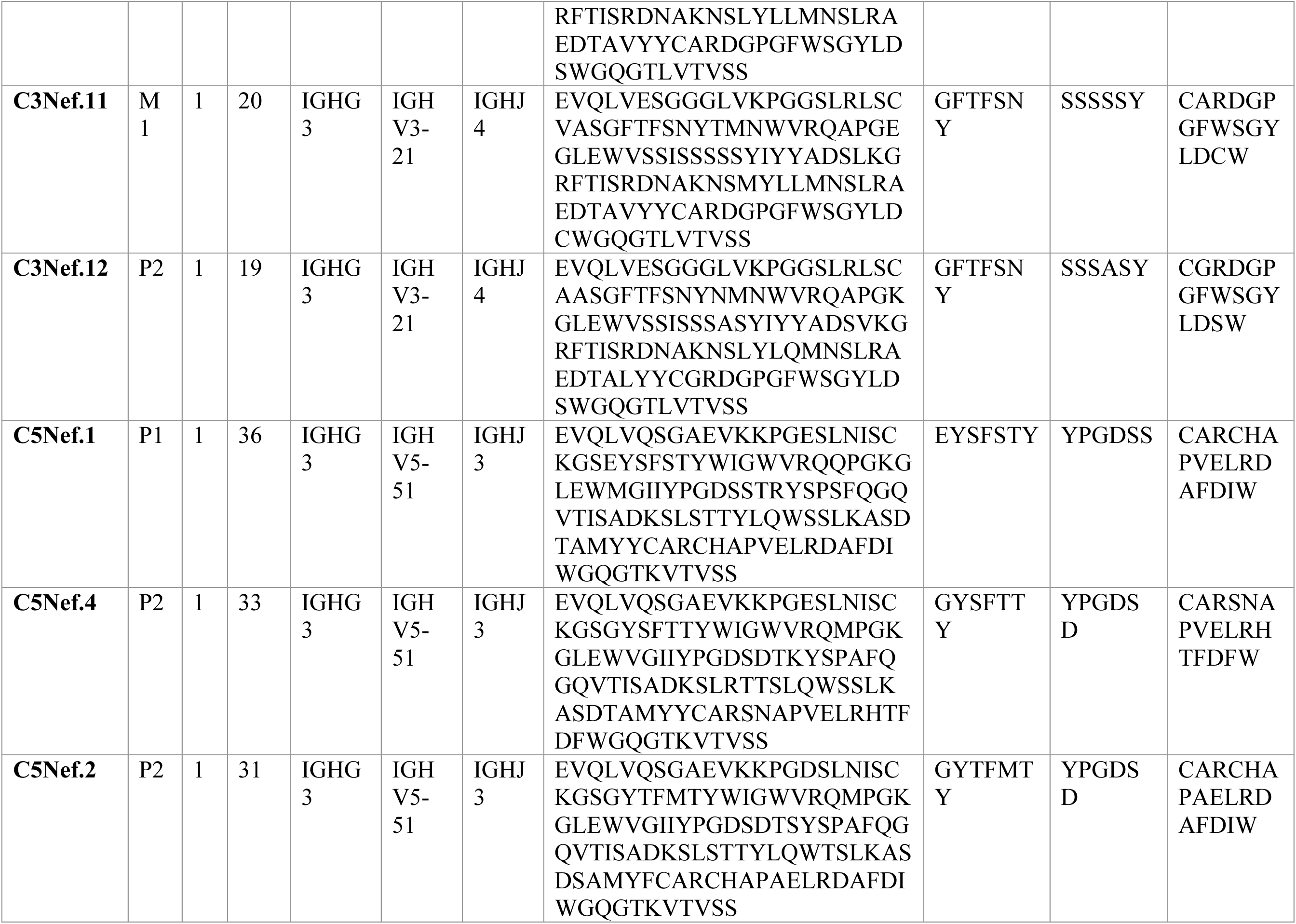
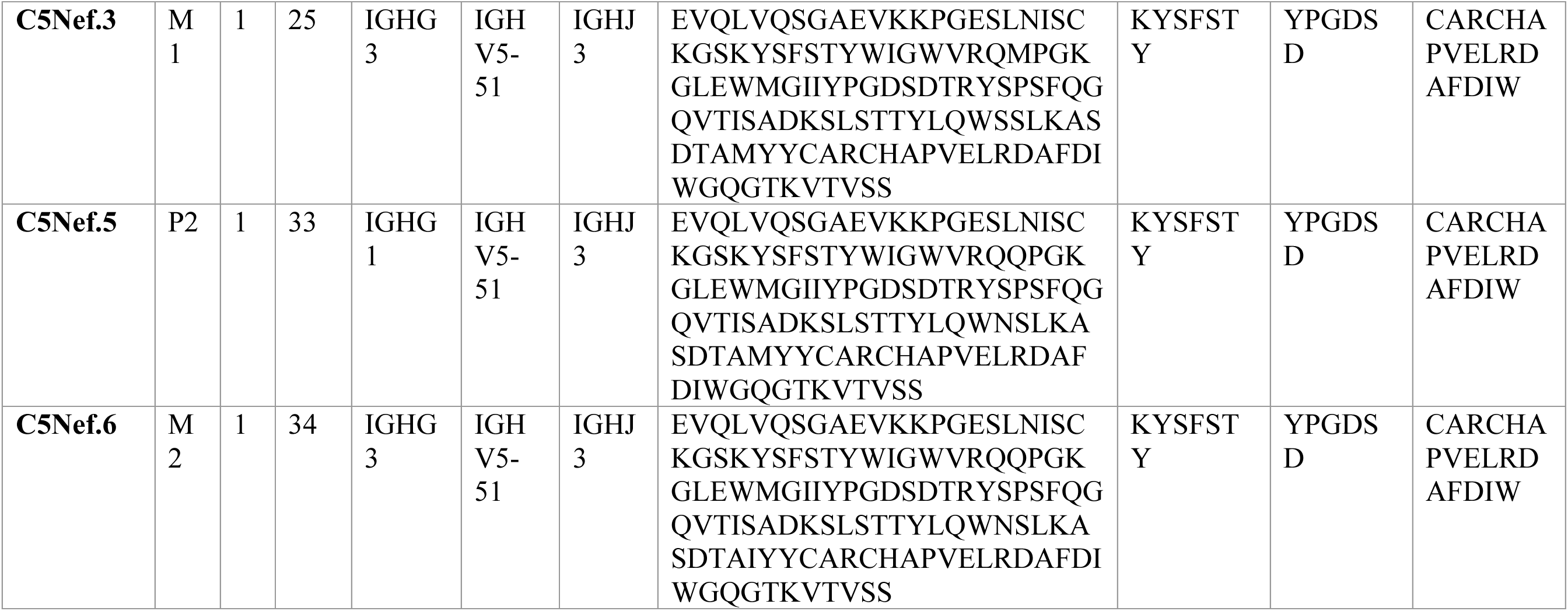
C3Nef and C5Nef V(D)J sequence information; heavy chain.

**Table S3.**
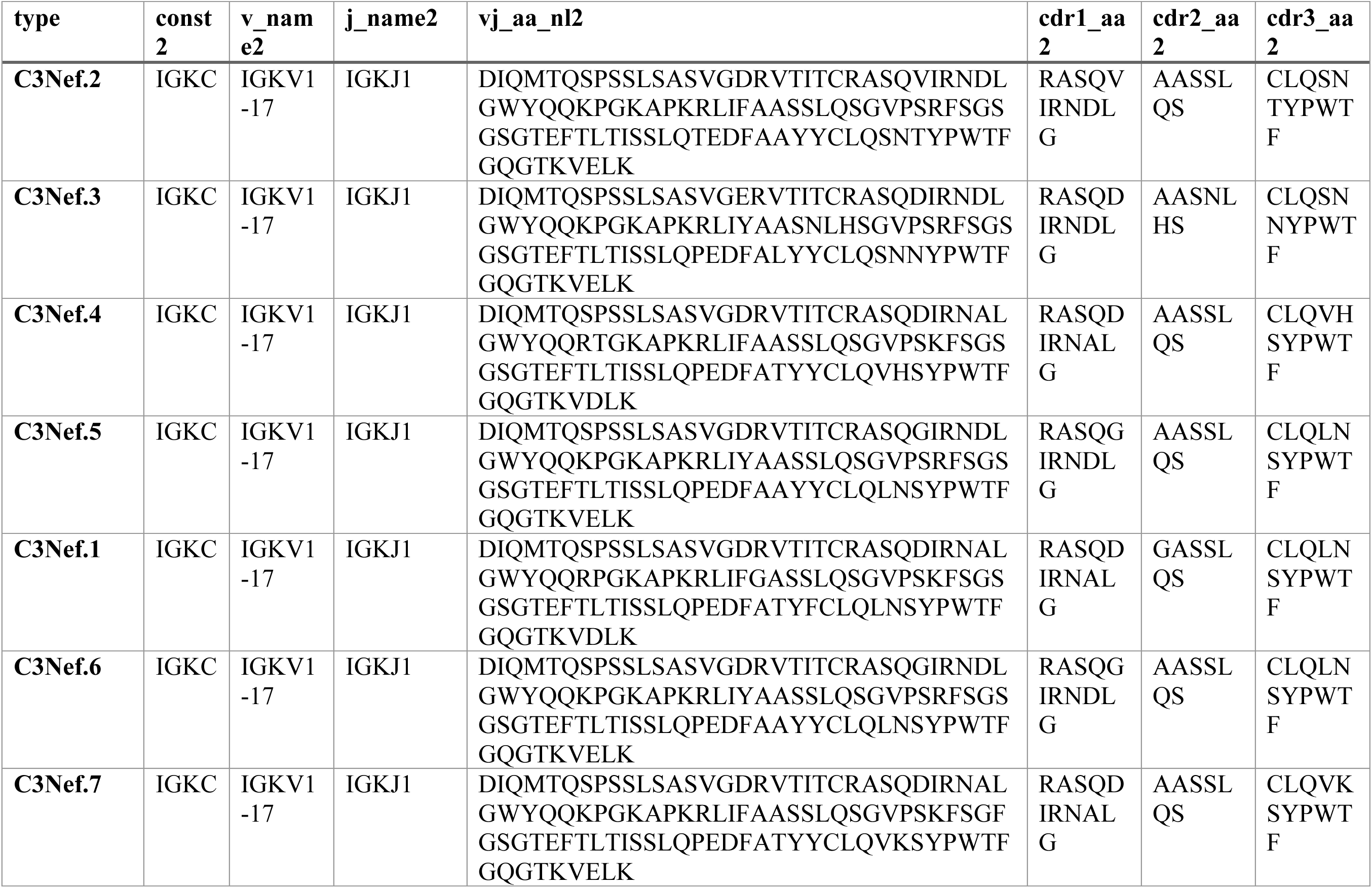
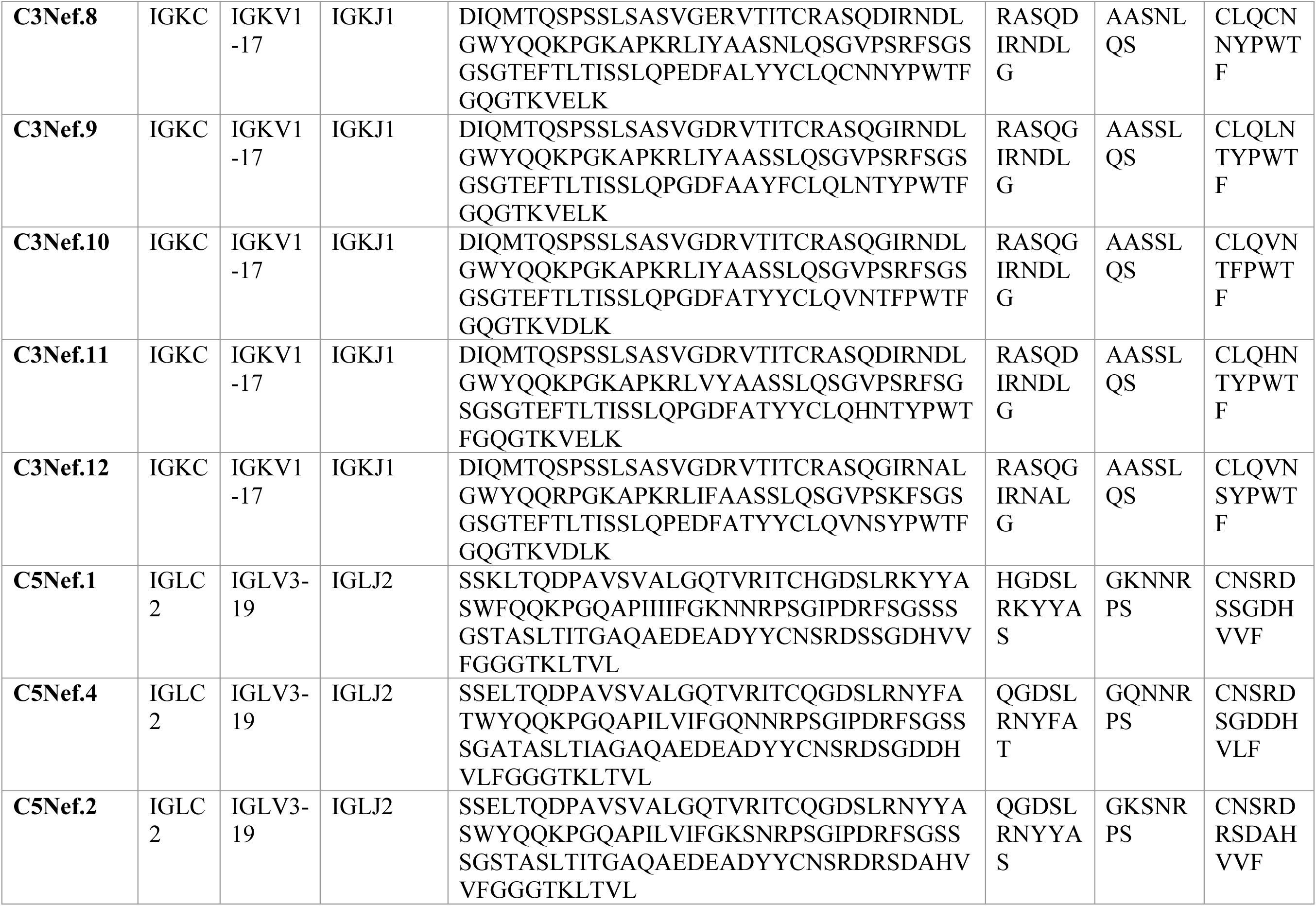
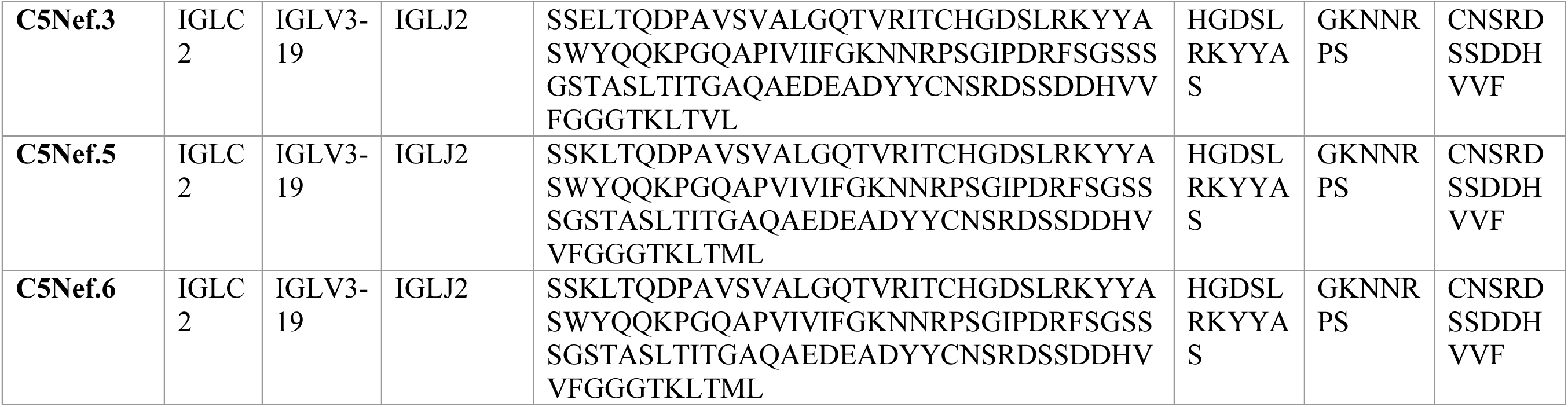
C3Nef and C5Nef V(D)J sequence information; light chain.

**Table S4.** Cryo-EM data collection and processing.

| **Data collection and processing** |  |  |
| --- | --- | --- |
| **Dataset** | **Dataset 1 (no tilt)** | **Dataset 2 (30° tilt)** |
| Microscope | Titan Krios | Titan Krios |
| Detector | Falcon 4i | K3 |
| Magnification (nominal) | 165,000x | 105,000x |
| Voltage (kV) | 300 | 300 |
| Spherical aberration (mm) | 2.7 | 2.7 |
| Total electron dose (e-/Å<sup>2</sup>) | 55 | 50 |
| Pixel size (Å) | 0.74 | 0.413 (super resolution) |
| Defocus range (um) | -2.4 to -0.9 | -2.2 to -1.0 |
| Energy filter | Selectris | Bioquantum |
| Energy filter slit width (eV) | 10 | 20 |
| Stage tilt | 0° | 30° |
| Number of Movies | 11,128 | 6,301 |
| Initial particle images (no.) | 971,220 | 194,086 |

**Table S5.**
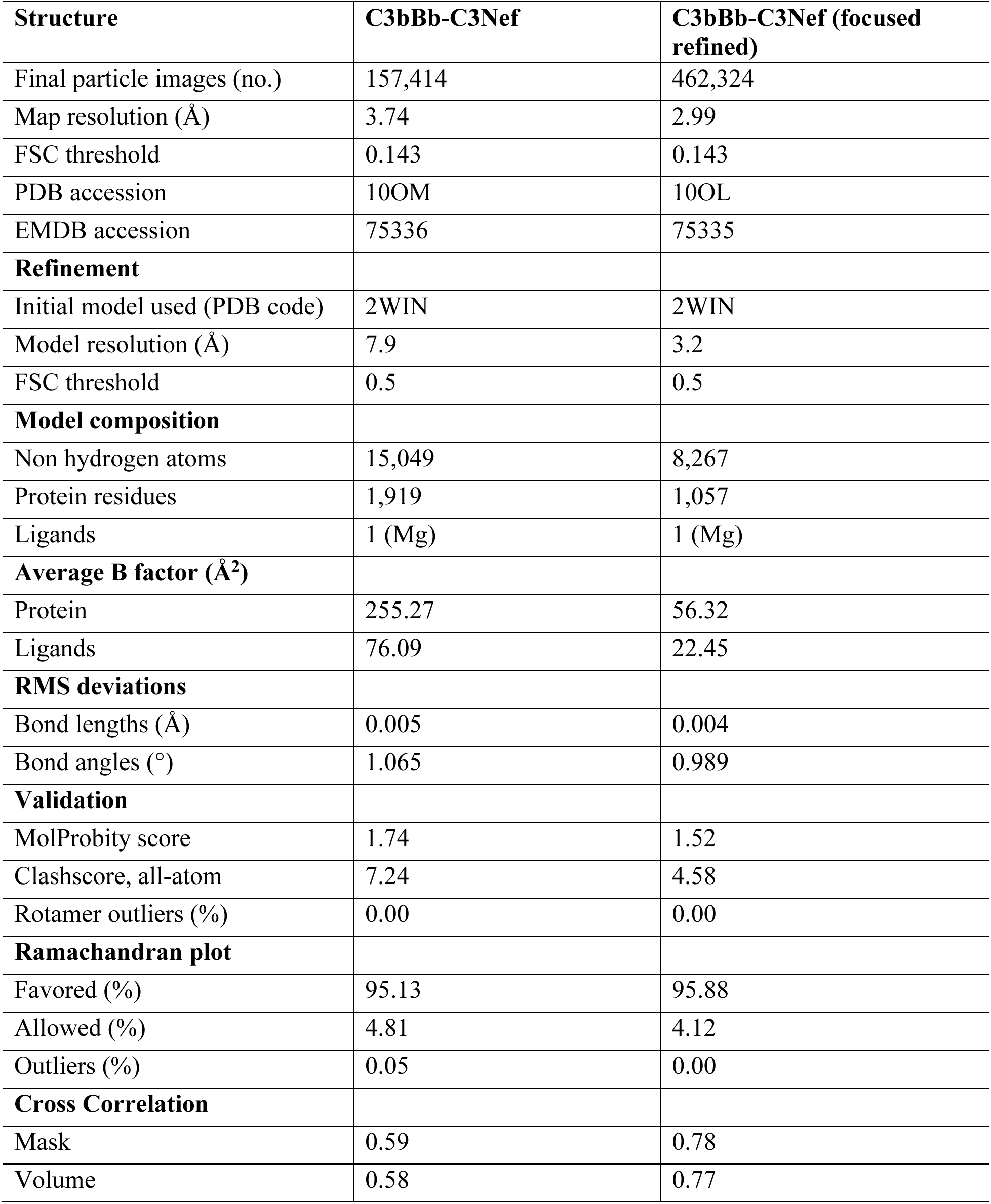
Cryo-EM model refinement and validation statistics.

| **Structure** | **C3bBb-C3Nef** | **C3bBb-C3Nef (focused refined)** |
| --- | --- | --- |
| Final particle images (no.) | 157,414 | 462,324 |
| Map resolution (Å) | 3.74 | 2.99 |
| FSC threshold | 0.143 | 0.143 |
| PDB accession | 10OM | 10OL |
| EMDB accession | 75336 | 75335 |
| **Refinement** |  |  |
| Initial model used (PDB code) | 2WIN | 2WIN |
| Model resolution (Å) | 7.9 | 3.2 |
| FSC threshold | 0.5 | 0.5 |
| **Model composition** |  |  |
| Non hydrogen atoms | 15,049 | 8,267 |
| Protein residues | 1,919 | 1,057 |
| Ligands | 1 (Mg) | 1 (Mg) |
| **Average B factor (Å<sup>2</sup>)** |  |  |
| Protein | 255.27 | 56.32 |
| Ligands | 76.09 | 22.45 |
| **RMS deviations** |  |  |
| Bond lengths (Å) | 0.005 | 0.004 |
| Bond angles (°) | 1.065 | 0.989 |
| **Validation** |  |  |
| MolProbity score | 1.74 | 1.52 |
| Clashscore, all-atom | 7.24 | 4.58 |
| Rotamer outliers (%) | 0.00 | 0.00 |
| **Ramachandran plot** |  |  |
| Favored (%) | 95.13 | 95.88 |
| Allowed (%) | 4.81 | 4.12 |
| Outliers (%) | 0.05 | 0.00 |
| **Cross Correlation** |  |  |
| Mask | 0.59 | 0.78 |
| Volume | 0.58 | 0.77 |

**Table S6.** Supplementary Table 5: Analysis of the interfaces between C3bBb convertase and C3Nef.

| Interface buried area |  |  |  |
| --- | --- | --- | --- |
| Interface |  |  | Buried area (Å²) |
| Bb | C3b (C345c) |  | 332.7 |
| Bb | C3Nef Heavy chain |  | 68.4 |
| Bb | C3Nef Light chain |  | 244 |
| C3b (C345c) | C3Nef Heavy chain |  | 538.3 |
| C3b (C345c) | C3Nef Light chain |  | 228.9 |
| C3Nef Heavy chain | C3Nef Light chain |  | 1701.2 |
| Residues on Interfaces |  |  |  |
| C3Nef residue | Regions | C3b (C345c) residue | Bb residue |
| N50 | CHRH1 | V1658, F1659 |  |
| S71 | CDRH2 | E1654 |  |
| S73 | CDRH2 | E1654 |  |
| Y76 | CDRH2 | G1650, E1654 |  |
| P120 | CDRH3 | F1659 |  |
| G121 | CDRH3 | F1659 |  |
| F122 | CDRH3 | S1655, L1529 |  |
| W123 | CDRH3 | P1662 |  |
| R50 | CDRL1 | E1530 |  |
| Y69 | CDRL2 |  | G395, G396 |
| S72 | CDRL2 |  | K356 |
| S73 | CDRL2 |  | K356 |
| L74 | CDRL2 |  | G396 |
| S80 | CDRL2 |  | I399, D403 |
| N112 | CDRL3 | K1644 |  |
| T113 | CDRL3 | Q1647, K1644 |  |
| Y114 | CDRL3 | Q1647 |  |

**Movie S1. Hypothetical Bb-rotation model of properdin-mediated C3bBb stabilization and C3Nef binding.** In this video, we use PDB structures 9U62, 10OL, and 10OM to demonstrate that Bb has a greater range of rotational motion relative to C3b than previously realized. In our model, properdin stabilizes the C3bBb convertase in the active Bb-down state by restricting the natural rotation of Bb centered around the MIDAS region. The absence of properdin permits rotation to the Bb-up position, resulting in a repositioning of the Mg^2+^ coordinating residues while maintaining the overall MIDAS architecture. The Bb-up position sterically clashes with properdin, exposes the full composite C3Nef epitope. Nef binding both stabilizes and inactivates the convertase. Bb in blue, C3b in white, C345c domain of C3b in orange, monomeric properdin (TSR5/6 shown) in yellow, C3Nef (Fab shown) in teal and aquamarine.

**Movie S2. Hypothetical Bb-rotation model of RCA-mediated C3bBb decay.** In this video, we use PDB structures 9U62, 2WIN, 5O35, 10OL, 10OM, and an AlphaFold3-generated structure of a predicted binding site between FH(1-6) and Bb to propose a Bb-rotational model of RCA-mediated decay. This model demonstrates how C345c pivots relative to C3b and swings Bb outward thus exposing the MG ring and “opening” the C3bBb convertase to factor H binding. Once factor H is bound, Bb “closes” and rotates upward and toward factor H, positioning known convertase-stabilizing variants of factor H and Bb in direct proximity. Contact of factor H with Bb disrupts the MIDAS region and irreversibly separates the vWA domain of Bb from the C345c domain of C3b resulting in complete convertase decay. C3b in white, factor H in orange, Bb in blue, factor H variants [FH(p.R53C), FH(p.Q81P)] and factor B [FB(p.K323E), FB(p.K323Q), FB(p.S367R), FB(p.D371G)] in red.

## References

1 Merle, N. S., Church, S. E., Fremeaux-Bacchi, V. & Roumenina, L. T. Complement System Part I - Molecular Mechanisms of Activation and Regulation. Front Immunol 6, 262 (2015). 10.3389/fimmu.2015.00262

2 Merle, N. S., Noe, R., Halbwachs-Mecarelli, L., Fremeaux-Bacchi, V. & Roumenina, L. T. Complement System Part II: Role in Immunity. Front Immunol 6, 257 (2015). 10.3389/fimmu.2015.00257

3 Mastellos, D. C., Hajishengallis, G. & Lambris, J. D. A guide to complement biology, pathology and therapeutic opportunity. Nat Rev Immunol 24, 118–141 (2024). 10.1038/s41577-023-00926-1

4 Pal, P., Wahi, P., Sahu, A. & Lal, G. Pro- and Anti-Inflammatory Role of Complement in Cancer. Eur J Immunol 55, e51767 (2025). 10.1002/eji.202451767

5 Nimmo, J. et al. The complement system in neurodegenerative diseases. Clin Sci (Lond) 138, 387–412 (2024). 10.1042/CS20230513

6 El Sissy, C., Rosain, J., Puel, M., Gonnin, C. & Fremeaux-Bacchi, V. Complement deficiencies and infections. Curr Opin Immunol 98, 102711 (2025). 10.1016/j.coi.2025.102711

7 Ricklin, D., Reis, E. S., Mastellos, D. C., Gros, P. & Lambris, J. D. Complement component C3 - The “Swiss Army Knife” of innate immunity and host defense. Immunol Rev 274, 33–58 (2016). 10.1111/imr.12500

8 Berends, E. T. et al. Molecular insights into the surface-specific arrangement of complement C5 convertase enzymes. BMC Biol 13, 93 (2015). 10.1186/s12915-015-0203-8

9 Mannes, M. et al. Complement inhibition at the level of C3 or C5: mechanistic reasons for ongoing terminal pathway activity. Blood 137, 443–455 (2021). 10.1182/blood.2020005959

10 Laursen, N. S., Magnani, F., Gottfredsen, R. H., Petersen, S. V. & Andersen, G. R. Structure, function and control of complement C5 and its proteolytic fragments. Curr Mol Med 12, 1083–1097 (2012). 10.2174/156652412802480925

11 Pickering, M. C. et al. C3 glomerulopathy: consensus report. Kidney Int 84, 1079–1089 (2013). 10.1038/ki.2013.377

12 Meuleman, M. S. et al. Acquired and genetic determinants of disease phenotype and therapeutic strategies in C3 glomerulopathy and immunoglobulin-associated MPGN. Nephrol Dial Transplant (2024). 10.1093/ndt/gfae245

13 Roquigny, J. et al. Acquired and genetic drivers of C3 and C5 convertase dysregulation in C3 glomerulopathy and immunoglobulin-associated MPGN. Nephrol Dial Transplant (2024). 10.1093/ndt/gfae243

14 Smith, R. J. H. et al. C3 glomerulopathy - understanding a rare complement-driven renal disease. Nat Rev Nephrol 15, 129–143 (2019). 10.1038/s41581-018-0107-2

15 Heidenreich, K. et al. C3 glomerulopathy: a kidney disease mediated by alternative pathway deregulation. Front Nephrol 4, 1460146 (2024). 10.3389/fneph.2024.1460146

16 Roquigny, J. et al. Functional Characterization of Anti-C3bBb Autoantibodies and C3 Glomerulopathy Phenotype. J Am Soc Nephrol (2024). 10.1681/ASN.0000000000000499

17 Daha, M. R., Fearon, D. T. & Austen, K. F. C3 nephritic factor (C3NeF): stabilization of fluid phase and cell-bound alternative pathway convertase. J Immunol 116, 1–7 (1976).

18 Gigli, I., Sorvillo, J., Mecarelli-Halbwachs, L. & Leibowitch, J. Mechanism of action of the C4 nephritic factor. Deregulation of the classical pathway of C3 convertase. J Exp Med 154, 1–12 (1981). 10.1084/jem.154.1.1

19 Halbwachs, L., Leveille, M., Lesavre, P., Wattel, S. & Leibowitch, J. Nephritic factor of the classical pathway of complement: immunoglobulin G autoantibody directed against the classical pathway C3 convetase enzyme. J Clin Invest 65, 1249–1256 (1980). 10.1172/jci109787

20 Daha, M. R., Kok, D. J. & Van Es, L. A. Regulation of the C3 nephritic factor stabilized C3/C5 convertase of complement by purified human erythrocyte C3b receptor. Clin Exp Immunol 50, 209–214 (1982).

21 Welsh, S. J., Zhang, Y. & Smith, R. J. H. Acquired drivers of C3 glomerulopathy. Clin Kidney J 18, sfaf022 (2025). 10.1093/ckj/sfaf022

22 Pangburn, M. K. & Muller-Eberhard, H. J. The C3 convertase of the alternative pathway of human complement. Enzymic properties of the bimolecular proteinase. Biochem J 235, 723–730 (1986). 10.1042/bj2350723

23 Michels, M., Volokhina, E. B., van de Kar, N. & van den Heuvel, L. Challenges in diagnostic testing of nephritic factors. Front Immunol 13, 1036136 (2022). 10.3389/fimmu.2022.1036136

24 Davis, A. E., 3rd, Ziegler, J. B., Gelfand, E. W., Rosen, F. S. & Alper, C. A. Heterogeneity of nephritic factor and its identification as an immunoglobulin. Proc Natl Acad Sci U S A 74, 3980–3983 (1977). 10.1073/pnas.74.9.3980

25 Williams, D. G., Bartlett, A. & Duffus, P. Identification of nephritic factor as an immunoglobulin. Clin Exp Immunol 33, 425–429 (1978).

26 Spitzer, R. E. et al. Serum C’3 lytic system in patients with glomerulonephritis. Science 164, 436–437 (1969). 10.1126/science.164.3878.436

27 Fakhouri, F. et al. Trial of Pegcetacoplan in C3 Glomerulopathy and Immune-Complex MPGN. N Engl J Med 393, 2210–2220 (2025). 10.1056/NEJMoa2501510

28 Syed, Y. Y. Iptacopan: First Approval. Drugs 84, 599–606 (2024). 10.1007/s40265-024-02009-4

29 Jaffe, D. B. et al. enclone: precision clonotyping and analysis of immune receptors. bioRxiv, 2022.2004.2021.489084 (2022). 10.1101/2022.04.21.489084

30 Stevens, K. H., Baas, L. M., van de Kar, N., van den Heuvel, L. & Michels, M. IdeZ protease does not prevent convertase stabilization by C3 nephritic factors in C3 glomerulopathy. Nephrol Dial Transplant 40, 1628–1631 (2025). 10.1093/ndt/gfaf065

31 Scott, D. M., Amos, N., Sissons, J. G., Lachmann, P. J. & Peters, D. K. The immunogloblin nature of nephritic factor (NeF). Clin Exp Immunol 32, 12–24 (1978).

32 Victor, K. D. et al. Nucleotide sequence of a human autoantibody to the alternative pathway C3/C5 convertase (C3NeF). Hybridoma 12, 231–237 (1993). 10.1089/hyb.1993.12.231

33 Rooijakkers, S. H. et al. Immune evasion by a staphylococcal complement inhibitor that acts on C3 convertases. Nat Immunol 6, 920–927 (2005). 10.1038/ni1235

34 Rooijakkers, S. H. et al. Structural and functional implications of the alternative complement pathway C3 convertase stabilized by a staphylococcal inhibitor. Nat Immunol 10, 721–727 (2009). 10.1038/ni.1756

35 Hourcade, D. E., Mitchell, L. M. & Oglesby, T. J. Mutations of the type A domain of complement factor B that promote high-affinity C3b-binding. J Immunol 162, 2906–2911 (1999).

36 Fishelson, Z. & Muller-Eberhard, H. J. C3 convertase of human complement: enhanced formation and stability of the enzyme generated with nickel instead of magnesium. J Immunol 129, 2603–2607 (1982).

37 Roumenina, L. T. et al. Hyperfunctional C3 convertase leads to complement deposition on endothelial cells and contributes to atypical hemolytic uremic syndrome. Blood 114, 2837–2845 (2009). 10.1182/blood-2009-01-197640

38 Pedersen, D. V. et al. Functional and structural insight into properdin control of complement alternative pathway amplification. EMBO J 36, 1084–1099 (2017). 10.15252/embj.201696173

39 Alcorlo, M., Tortajada, A., Rodriguez de Cordoba, S. & Llorca, O. Structural basis for the stabilization of the complement alternative pathway C3 convertase by properdin. Proc Natl Acad Sci U S A 110, 13504–13509 (2013). 10.1073/pnas.1309618110

40 Pedersen, D. V., Lorentzen, J. & Andersen, G. R. Structural studies offer a framework for understanding the role of properdin in the alternative pathway and beyond. Immunol Rev 313, 46–59 (2023). 10.1111/imr.13129

41 van den Bos, R. M., Pearce, N. M., Granneman, J., Brondijk, T. H. C. & Gros, P. Insights Into Enhanced Complement Activation by Structures of Properdin and Its Complex With the C-Terminal Domain of C3b. Front Immunol 10, 2097 (2019). 10.3389/fimmu.2019.02097

42 De la, O. B. K. I., Brondijk, T. H. C., Serna Martin, I. & Gros, P. Structural insights into C3 convertase activity of the classical pathway of complement. Nat Commun (2025). 10.1038/s41467-025-67730-4

43 Jia, C., Yang, X., Zhao, M. H., Tan, Y. & Xiao, J. Complement C3 recognition by C3 convertases. Sci Adv 12, eadz5404 (2026). 10.1126/sciadv.adz5404

44 Pedersen, D. V. et al. Structural Basis for Properdin Oligomerization and Convertase Stimulation in the Human Complement System. Front Immunol 10, 2007 (2019). 10.3389/fimmu.2019.02007

45 Pedersen, H. et al. A Complement C3-Specific Nanobody for Modulation of the Alternative Cascade Identifies the C-Terminal Domain of C3b as Functional in C5 Convertase Activity. J Immunol 205, 2287–2300 (2020). 10.4049/jimmunol.2000752

46 Donadelli, R. et al. Unraveling the Molecular Mechanisms Underlying Complement Dysregulation by Nephritic Factors in C3G and IC-MPGN. Front Immunol 9, 2329 (2018). 10.3389/fimmu.2018.02329

47 Marinozzi, M. C. et al. C5 nephritic factors drive the biological phenotype of C3 glomerulopathies. Kidney Int 92, 1232–1241 (2017). 10.1016/j.kint.2017.04.017

48 Zhang, Y. et al. Defining the complement biomarker profile of C3 glomerulopathy. Clin J Am Soc Nephrol 9, 1876–1882 (2014). 10.2215/CJN.01820214

49 Corvillo, F. et al. Serum properdin consumption as a biomarker of C5 convertase dysregulation in C3 glomerulopathy. Clin Exp Immunol 184, 118–125 (2016). 10.1111/cei.12754

50 Michels, M. et al. Long-term follow-up including extensive complement analysis of a pediatric C3 glomerulopathy cohort. Pediatr Nephrol 37, 601–612 (2022). 10.1007/s00467-021-05221-6

51 Michels, M. et al. The Role of Properdin in C5 Convertase Activity and C5b-9 Formation in the Complement Alternative Pathway. J Immunol 207, 2465–2472 (2021). 10.4049/jimmunol.2100238

52 Martín Merinero, H., et al. Functional characterization of 105 factor H variants associated with aHUS: lessons for variant classification. Blood 138, 2185–2201 (2021). 10.1182/blood.2021012037

53 Marinozzi, M. C. et al. Complement Factor B Mutations in Atypical Hemolytic Uremic Syndrome—Disease-Relevant or Benign? Journal of the American Society of Nephrology 25, 2053–2065 (2014). 10.1681/asn.2013070796

54 de Jorge, E. G. et al. Gain-of-function mutations in complement factor B are associated with atypical hemolytic uremic syndrome. Proceedings of the National Academy of Sciences 104, 240–245 (2007). doi:10.1073/pnas.0603420103

55 Roumenina, L. T. et al. Hyperfunctional C3 convertase leads to complement deposition on endothelial cells and contributes to atypical hemolytic uremic syndrome. Blood 114, 2837–2845 (2009). 10.1182/blood-2009-01-197640

56 Zhang, Y. et al. Mutation of complement factor B causing massive fluid-phase dysregulation of the alternative complement pathway can result in atypical hemolytic uremic syndrome. Kidney Int 98, 1265–1274 (2020). 10.1016/j.kint.2020.05.028

57 Madden, B. et al. Apolipoprotein E is enriched in dense deposits and is a marker for dense deposit disease in C3 glomerulopathy. Kidney Int 105, 1077–1087 (2024). 10.1016/j.kint.2024.02.013

58 Ponticelli, C., Calatroni, M. & Moroni, G. C3 glomerulopathies: dense deposit disease and C3 glomerulonephritis. Front Med (Lausanne) 10, 1289812 (2023). 10.3389/fmed.2023.1289812

59 Tarlinton, D. M., Ding, Z., Tellier, J. & Nutt, S. L. Making sense of plasma cell heterogeneity. Curr Opin Immunol 81, 102297 (2023). 10.1016/j.coi.2023.102297

60 Spitzer, R. E., Stitzel, A. E. & Tsokos, G. C. Evidence that production of autoantibody to the alternative pathway C3 convertase is a normal physiologic event. J Pediatr 116, S103–108 (1990). 10.1016/s0022-3476(05)82711-8

61 Thompson, R. A. IgG3 levels in patients with chronic membranoproliferative glomerulonephritis. Br Med J 1, 282–284 (1972). 10.1136/bmj.1.5795.282

62 Davis, A. E., 3rd, Gelfand, E. W., Schur, P. H., Rosen, F. S. & Alper, C. A. IgG subclass studies of C3 nephritic factor. Clin Immunol Immunopathol 11, 98–101 (1978). 10.1016/0090-1229(78)90207-6

63 Ambegaonkar, A. A., Holla, P., Dizon, B. L., Sohn, H. & Pierce, S. K. Atypical B cells in chronic infectious diseases and systemic autoimmunity: puzzles with many missing pieces. Curr Opin Immunol 77, 102227 (2022). 10.1016/j.coi.2022.102227

64 Dugan, H. L. et al. Profiling B cell immunodominance after SARS-CoV-2 infection reveals antibody evolution to non-neutralizing viral targets. Immunity 54, 1290–1303 e1297 (2021). 10.1016/j.immuni.2021.05.001

65 Spitzer, R. E., Stitzel, A. E. & Tsokos, G. C. Production of IgG and IgM autoantibody to the alternative pathway C3 convertase in normal individuals and patients with membranoproliferative glomerulonephritis. Clin Immunol Immunopathol 57, 10–18 (1990). 10.1016/0090-1229(90)90018-l

66 Zhang, Y. et al. Causes of alternative pathway dysregulation in dense deposit disease. Clin J Am Soc Nephrol 7, 265–274 (2012). 10.2215/CJN.07900811

67 Culek, C. T. & Nester, C. M. A Nef Precursor or Benign Counterpart: High Prevalence of Healthy Subjects with Antibody Reactivity to the C3-Convertase. Kidney360 4, 1615–1622 (2023). 10.34067/KID.0000000000000260

68 R: A language and environment for statistical computing. (Vienna, Austria, 2021).

69 RStudio: Integrated Development Environment for R (Posit Software, PBC, Boston, MA., 2025).

70 Edgar, R., Domrachev, M. & Lash, A. E. Gene Expression Omnibus: NCBI gene expression and hybridization array data repository. Nucleic Acids Res 30, 207–210 (2002). 10.1093/nar/30.1.207

71 Clough, E. et al. NCBI GEO: archive for gene expression and epigenomics data sets: 23-year update. Nucleic Acids Res 52, D138–D144 (2024). 10.1093/nar/gkad965

